# DeepHalo: Deep Learning-Powered Exploration of Halogenated Metabolites Uncovering Antibacterial Depsipeptides

**DOI:** 10.1101/2025.08.11.669588

**Authors:** Shanshan Chang, Xin Qi, Mengyuan Wang, Xinyue Huang, Ning He, Mingxu Chen, Jiahan Wang, Yu Du, Shuchen Wang, Yihong Li, Quanxiu Gao, Xinran Chen, Qing Lv, Xingxing Li, Bin Hong, Yunying Xie

## Abstract

In the omics era, confident high-throughput analytical tools are crucial for the efficient identification of metabolites. Here, we present DeepHalo, a deep learning-integrated and hierarchically optimized workflow designed for high-throughput exploration of halogenated metabolites from high-resolution mass spectrometry-based metabolomics. DeepHalo leverages deep learning models combined with a comprehensive scoring to enhance the reliability of halogen predictions. It integrates PyOpenMS for fast isotope pattern detection and incorporates a halogen-based dereplication algorithm with GNPS molecular networking to efficiently exploit and annotate halogenates from complex biological matrices. To validate its performance, DeepHalo was applied to explore halogenated metabolites from 1,296 microbial culture crude, leading to the discovery of six families of structurally diverse halogenated molecules. This included a new class of cyclic depsipeptides, aglomycins A-E, featuring rare 3-chloroanthranilic acid and/or epoxyvaline blocks. Additionally, a plausible biosynthetic pathway of aglomycins was proposed through bioinformatics analyses and targeted gene knockout experiments. Bioassays revealed that aglomycin A exhibits synergistic antibacterial activity with linezolid against vancomycin-resistant *Enterococcus faecium* (VRE) both *in vitro* and *in vivo*. We envision that DeepHalo (freely available at https://github.com/xieyying/DeepHalo) will become a powerful tool for accelerating the discovery of halogenated “dark matter”.

## Introduction

Halogenated natural products represent an important class of pharmacologically relevant compounds, as the incorporation of halogen atoms often significantly enhances their biological activity^1^. Additionally, halogens can also improve a substance’s metabolic stability and membrane permeability^1–2^. These combined advantages have made halogen substituents a common feature in over 25% of approved drugs^2^, highlighting their immense potential in drug development.

Nature, particularly microorganisms, harbors a wealth of halogenated metabolites awaiting exploration. Biogenically, halogens, primarily chlorine and bromine, are incorporated into substrates by diverse enzymes known as halogenases through electrophilic, radical, or, less commonly, nucleophilic mechanisms^3–4^. Flavin-Dependent Halogenase (FDHs) and iron(II)-*α*-Ketoglutarate-Dependent Halogenases (KDHs) are the most prevalent types, while new classes of halogenases continually being uncovered^3–5^. Genomic studies have revealed that microorganisms possess a vast repertoire of halogenase-encoding genes^6–8^. However, a significant gap exists between the prevalence of halogenase candidates and the number of identified halogenated metabolites^6–7^, suggesting that many such natural products remain undiscovered. Traditional natural product discovery approaches are hindered by high rediscovery rates and the frequent silencing of biosynthetic gene clusters (BGCs), necessitating the need for more efficient strategies. While genomics has advanced the exploration of halogenated compounds in the omics era^9^, its reliance on known halogenases limits the discovery of structurally diverse metabolites. In contrast, metabolomics provides a powerful alternative by directly profiling halogenated molecules, circumventing enzyme-specific constraints and enabling the identification of a broader range of halogenated natural products.

High-Resolution Mass Spectrometry (HRMS)-based metabolomics is a powerful tool for exploring halogenated metabolites. Statistical analysis reveals that chlorinated and brominated derivatives collectively account for over 98.5% of characterized halogenated natural products^10^. The unique isotopic distributions of chlorine (^35^Cl/^37^Cl) and bromine (^79^Br/^81^Br) atoms provide a valuable chemical signature for their identification based HRMS. Although specialized tools, such as MeHaloCoA^11^, Dynamic Cluster Analysis (DCAnalysis)^12^, HaloSeeker^13^, and ChloroDBPFinder^14^, have been developed for the targeted detection of halogenated compounds from MS data, it remains far beyond to achieve high-throughput and efficient discovery of these metabolites. The challenges primarily arise from three major obstacles. First, the confidence of halogen prediction has not yet met the requirements of high-throughput screening. Each culture extract typically yields hundreds to thousands of features^15^, requiring a analysis method with an exceptionally low false positive rate (ideally below 0.1%) to mitigate subsequent laborious investigations on erroneous detections. Technically, the inaccuracy mainly stems from inherent limitations in current isotope-based prediction models^11–14, 16^, and is further exacerbated by errors in the extracted isotope patterns, a critical flaw overlooked by all existing algorithm. Second, the speed of analysis is also a significant issue. Many existing approaches require several to dozens of minutes per sample^11–14^ and depend on heavily semi-automated^11, 13^ or manual interventions^12^ (Table S1), rendering them impractical for analysis of large-scale MS data. Finally, efficient dereplication is also critical to distinguish genuinely new halogenated natural compounds from known entities in complex chemical backgrounds. Unfortunately, most current methods fall short in this regard^11–12^, impeding the prioritization of truly novel compounds. Therefore, there is an urgent need for a more accurate, rapid, and automated computational approach to overcome these limitations and enable the efficient discovery of novel halogenated metabolites.

Herein, we present DeepHalo, a deep learning-integrated and fully automated workflow that seamlessly integrates the PyOpenMS^17^ and GNPS^18^ platforms, enabling the efficient and high-throughput discovery of novel halogenated natural products from complex biological matrices (Figure 1a). To demonstrate its capabilities, we applied DeepHalo to explore halogenated metabolites from 1,296 microbial culture crude (Figure 1b), resulting in the identification of a new class of halogenated cyclic peptides, aglomycins, along with five families of known halogenated molecules. Subsequent bioinformatics analyses and targeted gene knockout experiments disclosed a plausible biosynthetic pathway of aglomycins, while bioactivity assays revealed that algomycin A exhibits potent antibacterial activity against clinic multidrug-resistant *Enterococcus* strains and demonstrates synergistic effects with linezolid both *in vitro* and *in vivo*.

**Figure 1.**
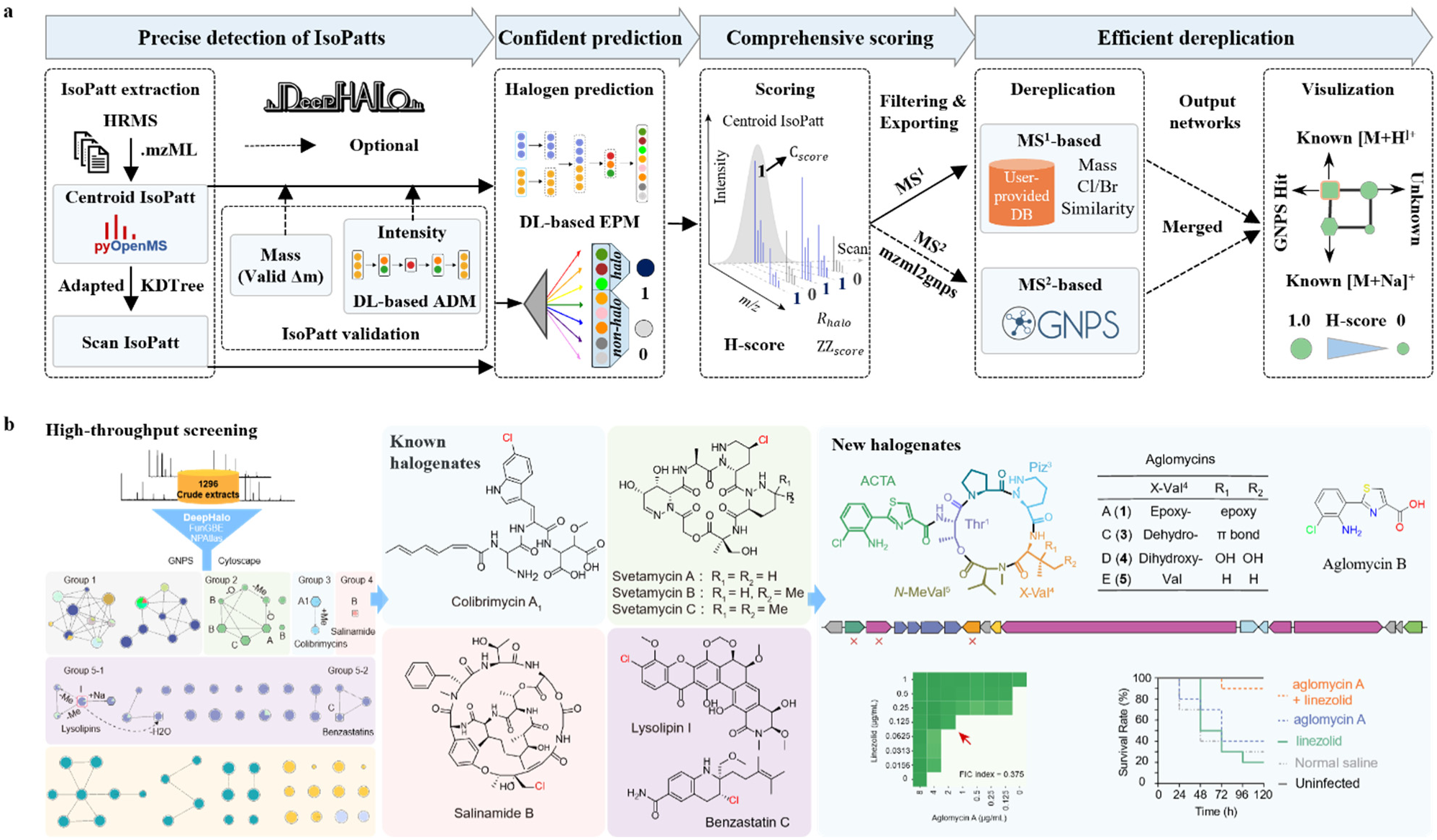
DeepHalo-assisted discovery of antibacterial depsipeptides. **a**, Schematic diagram of DeepHalo workflow. **b**, DeepHalo was applied for high-throughput screening of halogenated metabolites from 1,296 microbial culture crude, leading to the identification of five classes of structurally diverse halogenates, along with a new group of halogenated cyclic peptides, aglomycins.

## Results

### Overview of DeepHalo

DeepHalo was designed for automatic and efficient high-throughput mining of new halogenates from complex samples. To achieve this, we integrated four main functions into DeepHalo: precise detection of isotope patterns, confident halogen prediction, comprehensive scoring, and efficient dereplication (Figure 1a).

The core function of DeepHalo is to accurately predict the presence of halogens (Cl/Br) based on the isotope patterns of precursor ions. Toward this goal, DeepHalo employs a deep learning-based Element Prediction Model (EPM), trained on ∼109 million isotope patterns (Suppporting Information, Figure 2a-2b). To ensure the reliability of EPM, we implemented a series of optimization strategies. First, we generated synthetic isotope patterns to simulate co-eluting dehydrogen isomers (Figure S1) to enhance the model’s ability to filter out this common type of false signals. Second, we incorporated uncommon elements that frequently appear in natural products (Figure S2), not only to reduce false positive predictions for halogens but also to extend the model’s capability to detect other uncommon elements. Third, we developed a dual-branch Isotope Neural Network (IsoNN) that processes mass and intensity features of isotope peaks in parallel, to effectively address scale differences and allow for independent noise injection and scaling adjustments (Figure 2b). Finally, by combining manual tuning with Bayesian optimization, we efficiently optimized the model in just six iterations. The optimized EPM took relative intensities of the first five isotope peaks, along with the mass differences, Δm1 and Δm2, as input features, and output an 8-dimensional vector representing the predicted probabilities for the eight target categories (Figure 2b). These systematic enhancements enabled the EPM to achieve state-of-the-art performance in halogen prediction, as demonstrated by comparative evaluations on both simulated and real isotopic datasets (Table 1). Notably, our EPM was the only method to achieve 100% precision and recall, with zero FPR, for the binary classification of halogenated versus non-halogenated compounds on the tested real-world dataset. Even for the more detailed 8-category classification, EPM achieved 99.8% accuracy (622/623 correct), with only one Cl-type molecule misclassified as a Br-type (Figure 2d). Additionally, the comparative evaluation revealed a clear performance hierarchy (Table 1): deep learning-based models (our EPM and SIRIUS) significantly outperformed traditional machine learning-based tools (ChloroDBPFinder and DCAnalysis) and rule-based methods (HaloSeeker and MeHaloCoA), highlighting the superior capability of deep learning in resolving the complex, element-specific isotope patterns.

**Figure 2.**
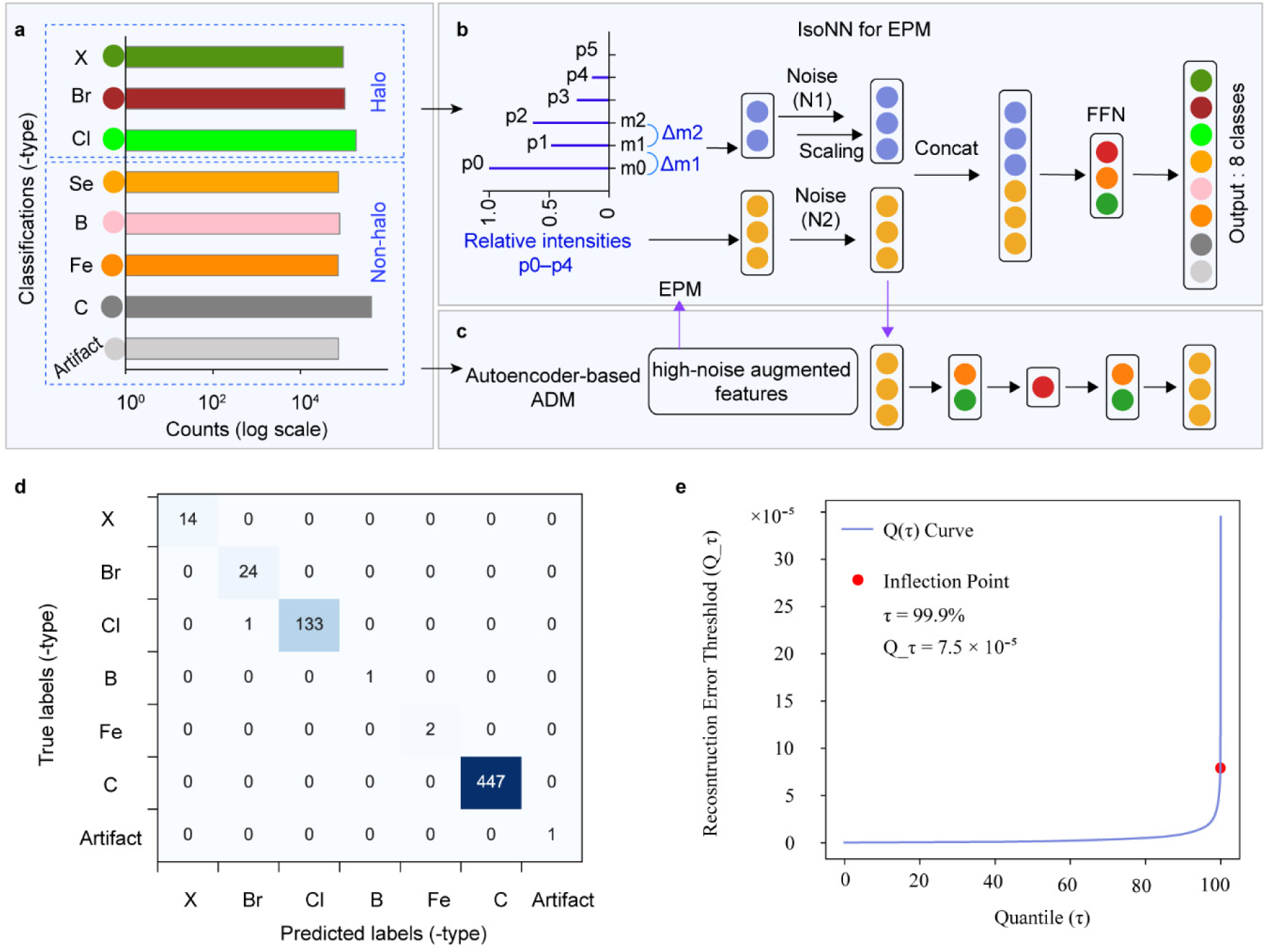
Establishment of deep learning models in the DeepHalo pipeline. **a**, Class distributions of isotope patterns used for training models (all types for the Element Prediction Model (EPM); X/Cl/Br/C-types for Anomalous Isotope Patterns Detection Model (ADM). Cl-type: Cl/Cl2; Br-type: Br/Cl3; X-type: mixed/polyhalogenated; B-type: boron-containing; Se-type: selenium-containing, and Fe-type: iron-containing; C-type: others; Artifact-type: simulated isotope patterns for co-eluting dyhydro isomers, (**b**-**c**) Network architectures: IsoNN for EPM (**b**) and autoencoder for ADM. **d**, Eight-category classification results of EPM tested on the CASMI2016_Myxo_plus dataset. **e**, Determination of ADM anomaly detection threshold (7.915×10^-5^). The optimal threshold was determined from the inflection point of the reconstruction error threshold-versus-percentile curve generated by ADM using the IsoHN dataset.

**Table 1.**
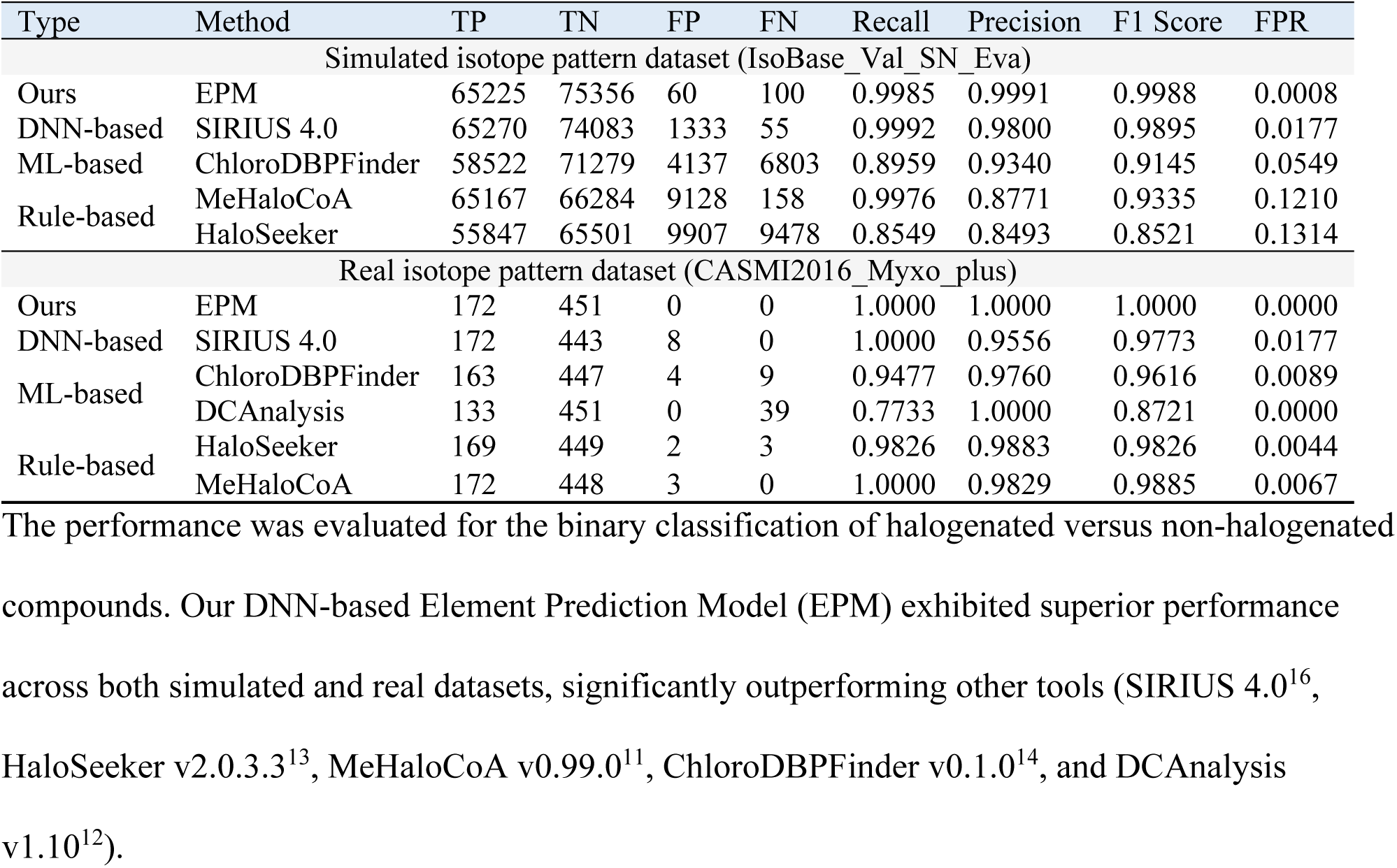
Performance comparison of methods for halogen prediction.

On the other hand, accurately detecting isotopic features from untargeted mass spectrometry data is a complex yet crucial step in the analytical workflow^19^. In the DeepHalo workflow, we employed pyOpenMS^17^ for feature detection and then applied a robust two-dimensional validation process that independently verifies both the mass and intensity dimensions. First, isotope patterns are corrected and validated by checking the mass differences between the first three peaks (Supporting Information). Only those with Δm1 and Δm2 within the 0.05th to 95.5th percentiles of the IsoHN dataset (Figrue S3), a high-noise augmented dataset containing halogenated (Cl/Br/X-type) and non-halogenated (C-type only) patterns, are retained for further analysis. Subsequently, for validation of isotopic intensities, we established a deep autoencoder-based^20^ Anomalous isotope pattern Detection Model (ADM) (Supporting Information, Figure 2c). Using this model, we computed reconstruction errors for intensities of the first five isotope peaks, which served to distinguish normal from anomalous distributions. The optimal discrimination threshold was set at 7.915×10^-5^, determined from the inflection point on the reconstruction error threshold-versus-percentile curve generated by ADM using the IsoHN dataset (Figure 2e).

Additionally, we introduced a halogen confidence score (H-score) as an indicator of the likelihood that a feature contains halogens. This metric comprehensively evaluates predictions at both the centroid and scan levels. To achieve this, we adapted pyOpenMS to extract isotopic patterns not only for each feature (centroid form) but also for each scan (Figure 1a). Subsequently, predictions were performed based on isotopic patterns of both levels, and the results were integrated for the calculation of H-score (Suppporting Information). This hierarchical assessment approach is a key distinguishing feature of DeepHalo, which not only enhances the reliability of halogen predictions but also allows flexible filtering of target features by adjusting the H-score threshold during analysis. Coupled with visualization tools, the H-score enables rapid differentiation and targeted annotation of halogenated candidates, even in highly complex datasets (Figure S4).

Automatic dereplication is essential for efficiently annotating known compounds and discovering novel ones from complex samples. In the DeepHalo pipeline, dereplication is achieved through two complementary methods (Figure 1a). 1) MS^1^-based dereplication: If the user provides a chemistry database, the pipeline will utilize MS^1^ data for dereplication by verifying the exact *m/z* of precursor ions, assessing the presence of Cl/Br, and calculating the cosine similarity between experimental and theoretical intensities. 2) MS^2^-based dereplication: If LC-HRMS data are acquired in DDA mode, the pipeline optionally exports the MS^2^ spectra of halogenated features based on a specified H-score threshold for further dereplication via molecular networking on the GNPS platform^18^. By leveraging the propagation of molecular networks, this method not only utilizes reference MS^2^ spectra from GNPS for dereplication but also eliminates false positives caused by the biotransformation of halogenates in blank matrices. Finally, all results are compiled into the network file output by GNPS and visualized using Cytoscape^21^, enabling intuitive and effective analysis of all outcomes (Figure S4). By combining these two methods, known compounds can be efficiently and accurately annotated, while potential new halogenated compounds are identified for further investigation.

Computational efficiency is a critical consideration for large-scale dataset analysis. To address this challenge, we employed a two-tier optimization strategy. First, we integrated pyOpenMS with KD-tree spatial indexing and concurrent.futures-based parallel processing, achieving an order-of-magnitude reduction in per-sample analysis time. Second, we unified all analytical steps (except dereplication) within a single “detect” function, thereby enabling fully automated processing of arbitrarily large LC-HRMS datasets, a capability essential for high-throughput applications.

Following the establishment of DeepHalo, we systematically evaluated the workflow using both simulated and real-world LC-HRMS datasets. First, we established a simulated LC-MS dataset SM1820-Base (Supporting Information) and used it to evaluate the effects of mass spectrometry parameters and isotope pattern validation on DeepHalo’s performance. The results (Figure 3) indicate that higher resolution and accuracy yield higher performance, with a resolution of 20,000 and an accuracy of 3 ppm, typically achieved by high-resolution QTof mass spectrometers, providing acceptable results. Although isotope pattern validation slightly reduces recalls, it substantially improves precisions and, consequently, the F1 scores. Next, we compared the performance of DeepHalo with state-of-the-art comparable tools (Table 2). Across all benchmarks, DeepHalo demonstrated consistent superiority, achieving the highest precision and recall, the lowest FPR, and significantly reduced computational time. Furthermore, its efficiency advantage becomes more pronounced as dataset size increases. For example, with the 145-sample CASMI 2022 dataset, competing methods failed to process the data within 12 hours, whereas DeepHalo completed the analysis in just 16.7 minutes with perfect accuracy and recall (100%).

**Figure 3.**
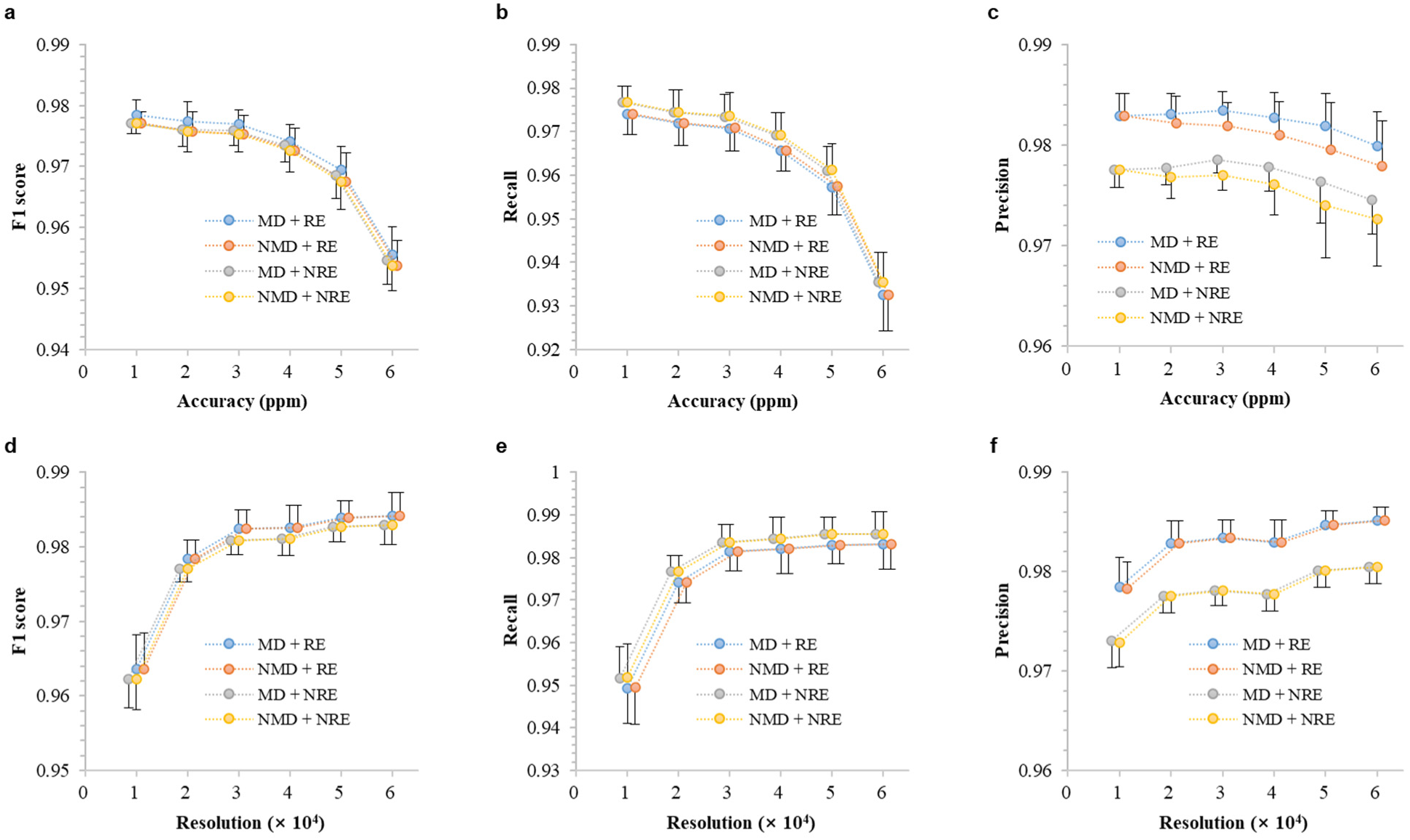
DeepHalo’s performance sensitivity to mass spectrometry parameters. The effects of MS accuracies (**a**–**c**) and resolutions (**d**–**f**) on Recalls, Precisions, and F1 scores. Key settings for DeepHalo analysis: MD/NMD: Mass Difference validation applied *vs*. not applied. RE/NRE: Reconstruction error (ADM-based) used *vs.* not used. *n* = 10 for each assay. Isotope pattern validation slightly reduces recalls but significantly improves precisions and F1 scores.

**Table 2.**
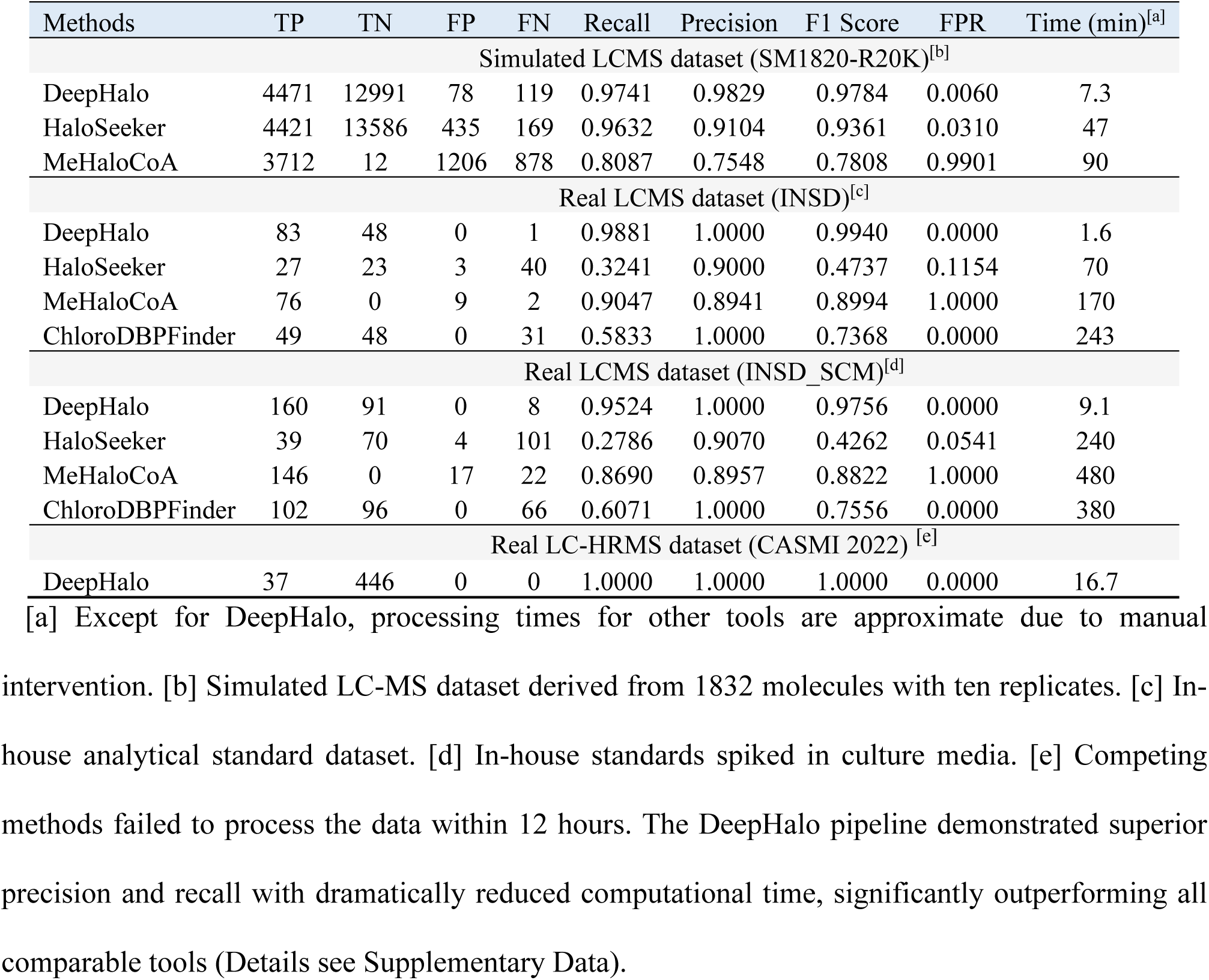
Performance comparison of our DeepHalo workflow with other counterpart methods.

### DeepHalo-assisted mining reveals a wealth of halogenated metabolites in *Streptomyces*

To demonstrate the efficiency of DeepHalo, we applied it to explore new halogenated compounds from 1296 HRMS-based metabolomic datasets derived from 53 *Streptomyces* and one *Micromonospora* strain cultures (Supporting Information). Initially, using the fully automatic ‘detect’ function of DeepHalo with H-score threshold set to 0.4, we identified and extracted potential features with halogen and their corresponding MS^2^ spectra from untargeted UPLC-HRMS/MS data of the crude extracts and 60 blank media controls. Following this, the similarity of the obtained MS^2^ spectra was analyzed using the molecular networking tool on the GNPS platform^18^. Subsequently, a deeper dereplication process was performed using the ‘dereplicate’ function of DeepHalo, employing NPAtlas^10^ (bacterial-derived data only) and MIBiG^22^ as dereplication databases. Ultimately, the results were visualized using Cytoscape for further analysis.

Based on the resulting molecular networks (MNs) (Figure 4a), blank media-derived variants (Group 1) were readily excluded according to the propagating nature of MNs. Singletons in Group 6 were then also recognized as media-derived variants based on careful examination of their precursors’ *m/z* and retention times. Concurrently, potential known compounds annotated using the user-provided database, highlighted as rectangles ([M+H]^+^) or hexagons ([M+Na]^+^) in Figure 4a, were efficiently identified. Consequently, five groups of known halogenated metabolites, svetamycins A-C, colibrimycin A1, salinamide B, lysolipin I, and benzastatin C, were detected from five strains, with one group (lysolipin I) confirmed via GNPS MS^2^ spectral matching. The structures of all these halogenates were subsequently confirmed through bioinformatics analysis combined with MS^2^ spectra or UV spectroscopy (Supporting Information, Figure S7-S10). Intriguingly, four of these five class compounds exhibit antibacterial or antiviral activities, with the chloride atom in benzastatin C proven to be critical for its antiviral activity^23^. As expected, these compounds display significant structural diversity while undergoing halogenation via distinct mechanisms. For example, the lipopeptide colibrimycin A1^24^ and the aromatic polyketide antibiotic lysolipin I^25^ are halogenated through FDH-mediated electrophilic attack on activated *sp*^2^ carbons, while depsipeptide antibiotic svetamycins^26–27^ feature KDH-catalyzed radical halogenation of unactivated *sp*^3^ carbons. In contrast, the chloride atom in the depsipeptide antibiotic salinamide B^28^ is introduced via non-enzymatic epoxide opening, and most intriguingly, the halogenation of benzastatin derivative benzastatin C^29^ involves an elusive cytochrome P450-catalyzed mechanism. Collectively, these results demonstrate DeepHalo’s efficacy in identifying structurally diverse halogenated metabolites, as well as its unique capability to uncover atypical halogenation pathways.

**Figure 4.**
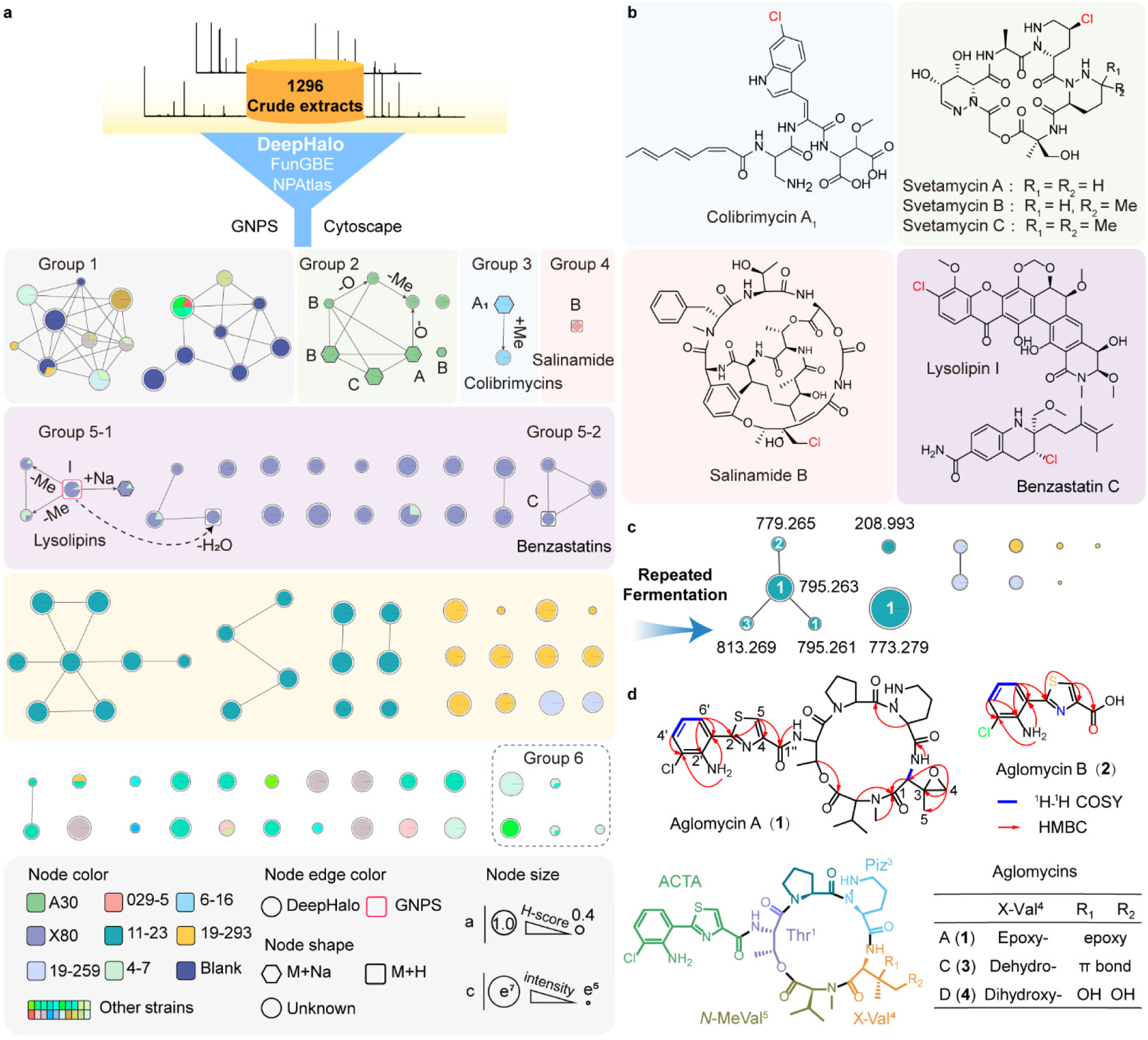
High-throughput mining of halogenated metabolites using DeepHalo. **a**, Molecular networks of potential halogenated compounds generated by mining 1296 crude extracts using DeepHalo, revealing that sixteen strains harbor halogenated features (Table S4). **b**, The structures of five groups of known halogenated compounds identified by DeepHalo. **c**, Molecular networks constructed from the potential halogenated compounds of three strains (11-23, 19-293, and 19-259) in flask-scale cultures reveals a dominant compound, aglomycin A, with an [M+H]^+^ ion at *m/z* 773.279 and an [M+Na]^+^ ion at *m/z* 795.263. **d**, The structure and key 2D NMR correlations of aglomycins A (**1**) and B (**2**) isolated from *Streptomyces* sp.11-23, together with aglomycins C (**3**) and D (**4**) identified through MS^2^ fragmentation (Figure. S34, Supplementary Information).

Further analysis revealed 11 additional strains producing halogenated metabolites (Figure 4a, Table S4), bringing the total to 16 strains, 30% of the 54 strains examined, which is considerably higher than previously believed^7-8^. Earlier studies, relying on PCR-based screening of flavin-dependent halogenases in actinomycetes strains, reported a detection rate of ∼20%^7-8^. Since most of our tested strains are *Streptomyces* spp., this discrepancy suggests that the potential for halogenated metabolite production in this genus may have been significantly underestimated. To further explore this hidden biosynthetic capacity, we sequenced the genomes of all 53 *Streptomyces* strains analyzed in this study. Using these sequences, we systematically investigated the distribution of halogenases by probing for two major classes: FDHs and KDHs using HMMer^30^. Strikingly, more than 79% (42 out of 53) of the strains encoded either FDHs or KDHs, with 36 strains harboring 82 FDHs and 16 strains containing 16 KDHs (Supplementary Data). Sequence similarity network (SSN) analysis revealed significant diversity among these halogenases, exhibiting broad substrate specificities (Figure. S11). To ascertain whether the high recovery rate of halogenases was universal or unique to our Tibetan strains, we analyzed potential halogenases in 2,110 *Streptomyces* genomic sequences downloaded from NCBI (as of January 14, 2021). The results showed that approximately 70% (1476 out of 2110) of the strains harbored halogenases, with 1324 strains containing 2297 FDHs and 388 containing 430 KDHs (Supplementary Data), consistent with the discovery rate observed in our in-house strains. These findings underscore the widespread presence of halogenases in the genus *Streptomyces* and point to a vast, untapped reservoir of halogenated natural products.

### Identification of a new class of halogenated depsipeptides

We prioritized the remaining potential halogenated natural products based on their yields and H-scores. Consequently, three strains (11-23, 19-293, and 19-259) were selected for repeated fermentation. While all three strains produced the target compounds (Figure 4c), only strain 11-23 yielded enough for further study. After optimizing the fermentation conditions (Figure S12), a scale-up fermentation combined with MS-guided targeted isolation yielded aglomycins A (**1**) and B (**2**).

Aglomycin A (**1**) was evidenced to have a molecular formula of C_35_H_45_ClN_8_O_8_S from HRESI(+)MS and ^13^C NMR data, requiring 17 double bond equivalents (DBE). Acquisition of 1D and 2D NMR data of **1** (Table S5) and detailed analysis suggested the presence of one threonine (Thr^1^), one proline (Pro^2^), one piperazic acid (Piz^3^), and one *N*-methylvaline (*N*-MeVal^5^). Additionally, a 3,4-epoxy-valine (epoxy-Val^4^) residue was deduced by the ^1^H-^1^H COSY correlation of its NH and H-2 (*δ*_H_ 5.22), combined with the key HMBC correlations from H-2 of epoxy-Val^4^ to its carbonyl carbon (C-1), H_2_-4 to C-2, C-3 (quaternary), and C-5 (isolated methyl), as well as characteristic chemical shifts of epoxy group (*δ*_C-3_ 57.4 and *δ*_C-4_ 49.6). Moreover, based on the ^1^H-^1^H COSY correlation of aromatic H-4’/H-5’/H-6’, and the key HMBC correlations from highly downfield-shifted H-5 (*δ*_H_ 8.37, s) to aromatic C-2 and C-4 as well as the carbonyl carbon (C-1’’), from NH_2_ to C-1’ and C-3’, from H-6’ to C-2, C-1’, C-2’, and from H-4’ to C-3’, combined with characteristic isotope distributions of the chlorine atom in MS (Figure S14), one 2-(2-amino-3-chlorophenyl)-4-thiazolecarboxylic acid (ACTA) residue was characterized. The connectivity among these six moieties was established by the key HMBC correlations (Figure 4d) and the MS^2^ fragmentations of the sodium adduct ion (Figure S15), ultimately confirming the planar structure of **1** (Figure 4d). The amide bond between Pro^2^ and Thr^1^ was determined to have a *trans* geometry based on the small Δ*δ*_C*β*-C*γ*_ value of 4.2 ppm (Table S5)^31–32^, whereas the amide bond between Pro^2^ and Piz^3^ was inferred to have a *cis* geometry based on the ROESY correlation of H-2-Pro^2^ and H-2-Piz^3^ (Figure S13)^33–34^. The absolute configuration of **1** was determined by combining the Marfey’s analysis^35^ with GIAO NMR calculations^36^ and DP4+ analysis^37^ (Supporting Information), ultimately confirming its amino acid residues as *S*-Thr^1^, *S*-Pro^2^, *S*-Piz^3^, *S*, *S*-epoxy-Val^4^ and *S*-*N*-Me-Val^5^.

Compound **2** with a much lower molecular weight ([M+H]⁺ *m/z* 254.9995) was determined by NMR ananlysis (Supporting Information) as a dipeptide corresponding to the truncated side chain of **1**. Additionally, two other minor components, aglomycins C (**3**) and D (**4**), were tentatively identified by MS^2^ due to their extremely low yields (Supporting Information and Figure S34). Both resemble **1**, except that epoxy-Val^4^ in **1** is replaced by dehydro-Val^4^ in **3** and dihydroxy-Val^4^ in **4** (Figure 4d).

### Deciphering the biosynthesis of algomycin A

Structurally, aglomycin A (**1**) is characterized by a cyclic pentapeptide core framework incorporating an exceptionally rare epoxyvaline residue, as well as a dipeptide side chain formed by the condensation of 3-chloroanthranilic acid (3-Cl-AA) and cysteine. Given the distinctive structure of **1**, the producer strain, *Streptomyces* sp. 11-23, was sequenced to elucidate its biosynthetic pathway. Biogenetically, the 3-Cl-AA moiety is predicted to be derived from tryptophan, which subsequently undergoes halogenation catalyzed by a tryptophan halogenase. Therefore, we utilized the tryptophan halogenase, PrnA^38^, as a probe, together with comprehensive bioinformatics analysis, revealing a putative *agl* cluster (Figure 5a). This cluster encodes two discrete giant NRPSs, AlgN and AglJ, holding two and five modules, respectively, expected to be responsible for the construction of the dipeptide side chain and the cyclic pentapeptide core scaffold (Figure 5b). The CAL domain of the first module of AlgN was expected to activate the precursor of 3-Cl-AA and transfer it to the adjacent T domain^39^. However, detailed bioinformatic analysis revealed that CAL harbors a mutation in the A10 motif, where the conserved lysine, crucial for adenylation activity^40^, is replaced by proline (Figure S35). Concurrently, *aglM* encoding an NPRS with a stand-alone A domain was found to be adjacent to *algN*, possibly compensating the adenylation function of the CAL and activate 3-Cl-AA. Intriguingly, a C_starter_ domain, typically found in lipid peptides to condense a fatty acid and the first amino acid residue for initiating the elongation of the peptide chain^41^, is located in the first module of AglJ (Figure 5b and Figure S36) and proposed to condense dipeptide acyl and threonine acyl to initiate the biosynthesis of cyclic pentapeptide. Notably, the fourth module of NRPS AglJ lacks an A domain and comprises only a C-T didomain, whereas a separate A-T didomain NPRS AglB and a free-standing type II thioesterase (TEII) AglI, were present in the *agl* cluster (Figure 5b and Figure S37). This modular configuration, reminiscent of the assembly lines of WS9326A^42^, legonmycin^43^, and rotihibin^44^, is predicted to cooperatively incorporate the unusual amino acid moiety, epoxy-Val^4^. This hypothesis was further verified through an in-frame deletion experiment targeting *aglB*. The *ΔaglB* knockout strain completely abolished production of the cyclic peptide **1** and its cyclic analogues, while leading to accumulation of its truncated dipeptide derivative **2** (Figure 5c). Moreover, metabolite analysis of the complemented strain (C-Δ*aglB*) revealed the restoration of **1** production concurrent with the disappearance of **2**, indicating that *aglB* is involved in the biosynthesis of the core pentapeptide scaffold of **1**.

**Figure 5.**
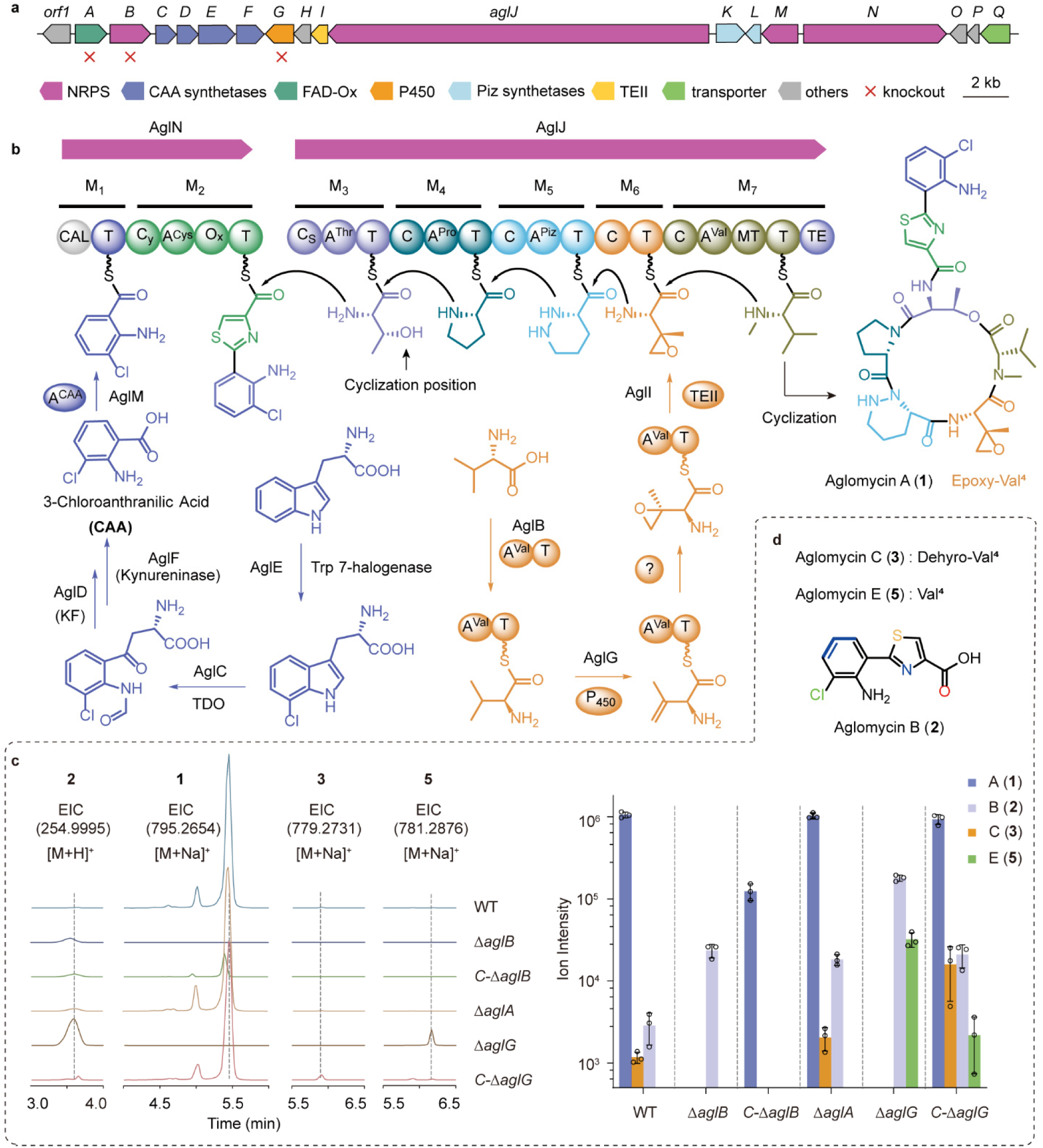
Proposed and identification of aglomycin A biosynthesis. **a**, Proposed aglomycin A biosynthetic gene cluster, confirmed by *aglA/B/G* knockout experiments. **b**, A plausible aglomycin A biosynthesis pathway. TDO: tryptophan 2,3-dioxygenase, KF: kynurenine formamidase. **c**, Extracted ion chromatograms (EICs) of compounds **1**, **2**, **3**, and **5** from wild type (WT), knockout, and complemented strains (C-strain) and their relative yields (n=3) **d**, Chemical structures of aglomycin derivatives.

Apart from the NRPSs responsible for core-scaffold biosynthesis, the genes encoding enzymes for the unusual building blocks are also identified in *agl.* AglK and AglL, exhibiting 62% and 51% amino acid sequence identities, respectively, to their homologues KtzI and KtzT^45–46^, are proposed to catalyze Piz biosynthesis. The precusor 3-Cl-AA is presumed to be derived from tryptophan through sequential catalysis by AglE (tryptophan halogenase), AglC (tryptophan 2,3-dioxygenase,), AglD (kynurenine formamidase), and AglF (kynureninase), following a route partly analogous to kynurenine pathway of tryptophan metabolism and that for antidepressant prodrug *L*-4-chlorokynurenine (*L*-4-Cl-Kyn) in taromycins^47^ (Figure 5b). Intriguingly, bioinformatics analysis of *Streptomyces* sp. 11-23 genome sequence disclosed a second triad of homologues of AglC/D/F, which are likely to involved in the *L*-Trp metabolism^48^ (Figure S38). AglC/D/F appear to have diverged from their canonical counterparts after gene duplication and coevolved with the tryptophan 7-halogenase, AglE, to specifically process 7-Cl-Trp. To our best knowledge, AglC represents the first reported TDO preferring 7-Cl-Trp, which complements the recently identified TDO acting on 6-Cl-Trp^47^ and expands the biosynthetic toolbox for halogenated tryptophan-derived molecules^49^.

Given the extreme rarity of epoxy valine in natural products, reported only recently in a patent for lipopeptide streptocinnamide B (Scm B)^50^, we conducted a comprehensive analysis alongside targeted gene knockout experiments to interrogate its biosynthesis pathway. Epoxidation of alkyl carbons, a process rarely observed in nature^51–53^, is hypothesized to occur either through a stepwise dehydrogenation–epoxidation cascade^51^ or via a hydroxylation–epoxidation route^53^. In the wild-type strain, we detected **3**, featuring dehydro-Val^4^, while no analogues containing hydroxy-Val^4^ were observed, suggesting that the epoxy group might be formed via a dehydrogenation–epoxidation cascade. In the aglomycin pathway, only two oxygenases remain without assigned functions, AlgA (a FAD-dependent oxidase) and AlgG (a cytochrome P450 monooxygenase), which are proposed to jointly mediate the two required oxygenation steps. To test this hypothesis, we individually inactivated *aglA* and *aglG*, through in-frame deletion in *Streptomyces* sp. 11-23 wild-type strain. Unexpectedly, the production of **1** was almost unaffected in Δ*aglA* mutant. In contrast, the Δ*aglG* mutant completely abolished the production of **1** and **3**, while leading to prominent production of **2** and a new halogenated compound **5** (named aglomycin E) with *m/z* 781.2876 [M+Na]^+^. Subsequent MS^2^ fragmentation analysis identified **5** as an analogue of **1**, featuring Val^4^ instead of epoxy-Val^4^ (Figure 5d and Figure. S34). Reintroduction of *aglG* into its mutant strain (C-Δ*aglG*) restored the production of **1** and **3** while dramatically reducing the levels of **2** and **5** (Figure. 5c). These results indicate that AglG is involved in forming the epoxide moiety of epoxy-valine, likely through a dehydrogenation step. However, the detailed biosynthetic mechanism remains elusive and warrants further investigation.

### The synergistic antibacterial effects of aglomycin A (1) and linezolid

We evaluated the antibacterial activity of purified aglomycins A (**1**) and B (**2**) and find them displaying potent antibacterial effects against Gram-positive bacteria with minimum inhibitory concentrations (MICs) ranging from 4 to 8 *μ*g/mL (Figure 6a). Subsequently, we further investigated their antibacterial activity against nine multidrug-resistant *Enterococcus* strains isolated from clinical samples. Both aglomycins A and B exhibited potent activity against clinical isolates of *E. faecium* and *E. faecalis* (Figure 6b). Notably, aglomycins A and B were effective against VRE regardless of the *van*A or *van*B genotype, suggesting their potential as a novel therapeutic option for combating drug-resistant enterococcal infections.

**Figure 6.**
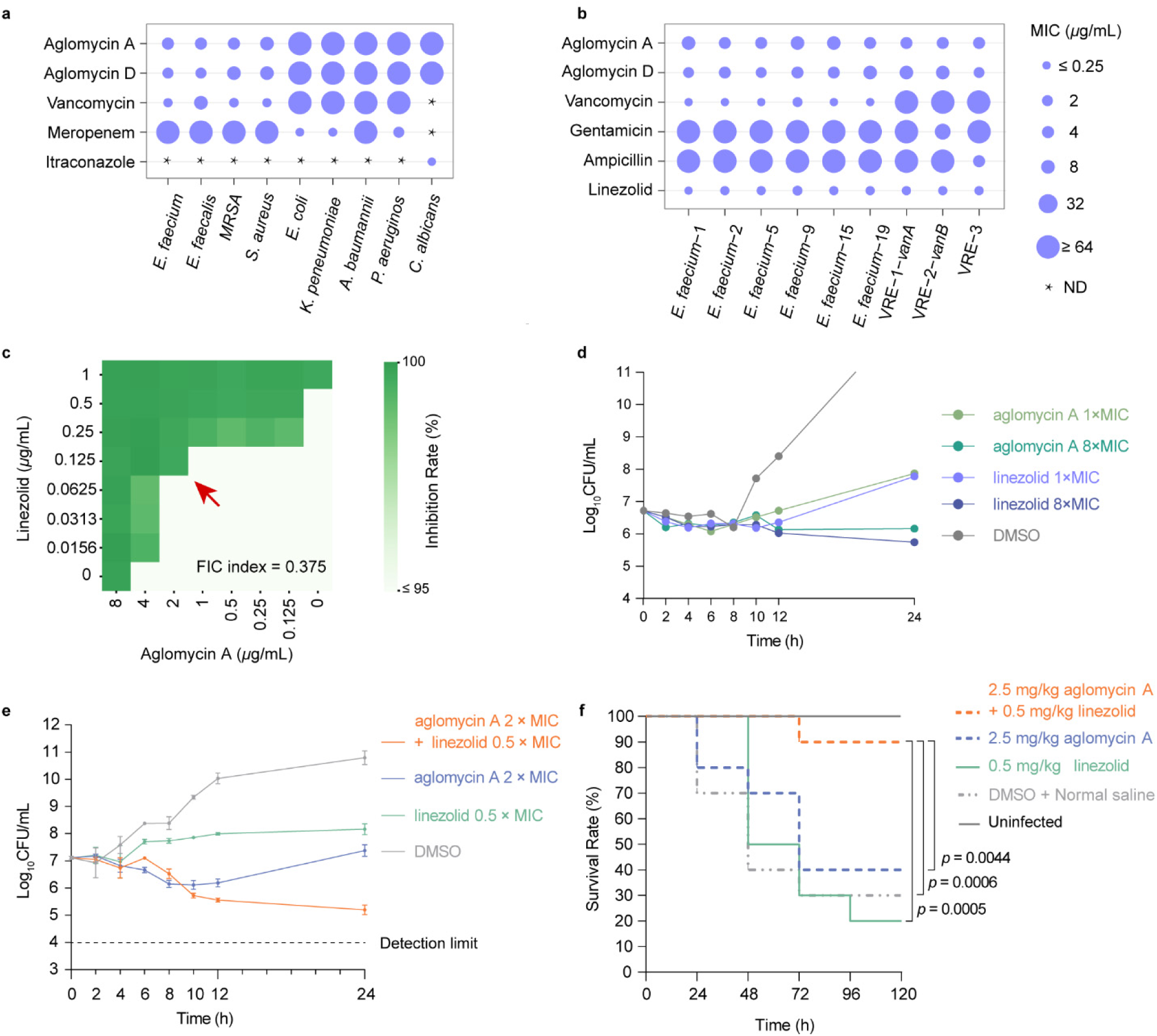
**a**, MIC values against various microbes. ND: not detect. **b**, MIC values against various drug-resistant *Enterococci* clinical isolates. **c**, Checkerboard assay of aglomycin A with linezolid against *E. faecium*; dark green indicates higher inhibition. **d**, Time-kill curve of aglomycin A and linezolid alone on *E. faecium*. Data represent the mean of two technical replicates with similar results. **e**, Time-kill curve of aglomycin A in combination with linezolid on *E. faecium*. Data represent mean of three technical replicates. **f**, The synergistic antibacterial effects of aglomycin A and linezolid in a *G. mellonella* infection model. Cox regression analysis was performed using R software (version 4.2.2), along with MSTATA software (https://www.mstata.com/).

Next, we investigated the potential synergistic or addictive effects of **1** in combination with clinically used antibiotics against *Enterococci*, including linezolid, quinupristin, dalfopristin, and daptomycin (Figure 6c and Figure S40), with **2** excluded for significant cytotoxicity (Figure S41a). Results disclosed that only the combination of **1** and linezolid demonstrated a fractional inhibitory concentration index below 0.5, specifically 0.375, indicating a synergistic effect against VRE. Furthermore, time-kill kinetics assay demonstrated that **1** exerted time-dependent bacteriostatic effects comparable to those of linezolid (Figure 6d and Figure S41b). When combined with linezolid,

- significantly enhanced bactericidal activity at 24 h, reducing bacterial counts by approximately 1–
- log_10_CFU/mL (Figure 6e and Figure S41c). These findings confirm that **1** and linezolid exhibit synergistic effects, and their combination significantly enhances antimicrobial efficacy against *E. faecium*.

Then, we employed a *Galleria mellonella* infection model^54^ to evaluate **1** *in vivo* synergistic antibacterial activity in combination with linezolid. As shown in Figure 6f, co-administration of **1** (2.5 mg/kg) and linezolid (0.5 mg/kg) to *E. faecium*-infected larvae significantly improved survival rate compared to the DMSO in saline vehicle (*p* = 0.0006) and to monotherapy with either aglomycin A (*p* = 0.0044) or linezolid (*p* = 0.0005), both of which provided only limited protection. This synergistic effect was further confirmed by co-administration of both drugs at 1 mg/kg each (Figure S41d). These results further reinforce the therapeutic potential of aglomycin A in combination with linezolid for combating infections with drug-resistant *Enterococci*.

## Discussion

Halogenated compounds constitute an important class of drug-like molecules^2^, with microorganisms representing a major natural source^10, 55^. In biological systems, halogenation is generally catalyzed by diverse halogenases, which are widely distributed in nature^3-4^. However, a significant discrepancy exists between the prevalence of halogenases and the number of identified natural halogenated compounds, suggesting that many such molecules remain undiscovered^6-7^. Traditional approaches to their exploration are hampered by high rediscovery rates and the frequent inactivity of biosynthetic gene clusters (BGCs)^56^, highlighting the need for efficient, high-throughput methods for systematic analysis.

To address these challenges, we developed DeepHalo, a deep learning-powered workflow optimized for high-throughput mining of halogenated molecules in large-scale LC-HRMS data. The workflow features five key advancements that collectively enhance both the confidence and efficiency of halogenated molecule identification. First, we developed a deep learning-based element prediction model (EPM) that achieves state-of-the-art performance in detecting halogenated compounds by analyzing their distinctive isotopic patterns. Second, we implemented an optional isotope pattern validation step that significantly improves prediction accuracy by mitigating errors in isotope peak extraction. Although this type of validation has been used for detecting compounds with common elements, DeepHalo is the first to extend this approach to the detection of halogenated compounds. Third, DeepHalo introduces a hierarchical halogen confidence score (H-score) that integrates centroid- and scan-level isotopic pattern analyses to reliably assess halogenation likelihood. This innovative metric enables adjustable threshold filtering and visualization-guided prioritization of halogenated features in complex metabolomic datasets. Fourth, by integrating MS^1^ dereplication (optimized for halogenated compounds) with GNPS-compatible MS^2^ molecular networking, our workflow achieves both high accuracy and efficiency in annotating halogenated molecules. Fifth, through computational optimizations, DeepHalo achieves unmatched efficiency, processing large-scale LC-HRMS datasets orders of magnitude faster than conventional tools. Collectively, these advancements establish DeepHalo as a transformative workflow for halogenate metabolomics, enabling high-throughput discovery and identification of halogenated compounds with unprecedented speed and accuracy.

Following the development of DeepHalo, we employed this workflow to systematically investigate halogenated metabolites in 1,296 LC-HRMS/MS datasets obtained from 54 actinomycete strains. The analysis revealed an unexpectedly high prevalence of halometabolite production, with 16 strains (30%) yielding detectable halogenated compounds. DeepHalo successfully identified five structurally distinct classes of known halogenated metabolites in these strains, all verified through HRMS, MS^2^, UV spectroscopy, and bioinformatics analyses. Moreover, genome mining targeting halogenases corroborated the high detection rate of halogenated metabolites and revealed a vast, untapped reservoir within the genus *Streptomyces*. These findings further demonstrate DeepHalo as an effective tool for halogenated compound discovery.

Additionally, from prioritized candidates, we characterized a novel cyclic depsipeptide, aglomycin A, along with its three derivatives from *Streptomyces* sp. 11-23. Aglomycin A features two rare structural blocks: 3-chloroanthranilic acid, first identified in natural products despite its common use in chemical synthesis, and an epoxide valine moiety, an exceptionally rare residue documented only recently in a patent^50^. Given the characteristic structure of aglomycin A, we subsequently investigated its biosynthesis using bioinformatics analyses and targeted gene knockout experiments. Bioinformatics analysis clearly disclosed that 3-chloroanthranilic acid is derived from tryptophan via the catalysis of the enzyme quartet AglE/C/D/F, with AglC representing the first reported TDO preferring 7-Cl-Trp. In addition, gene knockout experiments indicated that the epoxide moiety is formed through a dehydrogenation–epoxidation cascade mediated by AglG, a cytochrome P450 enzyme rarely reported to catalyze such a reaction^57^. Notably, using AglG as a probe, we identified a series of diverse biosynthetic gene clusters characterized by a conserved arrangement of genes encoding an A-T didomain NPRS (most predicted to selectively activate valine), a cytochrome P450, and a TEII (Fig. S37). These findings suggest that the occurrence of epoxidized valine in natural products may be underestimated, while the AglG offers a practical genetic probe for their systematic discovery.

Finally, we evaluated the antibacterial activity of aglomycins and found that aglomycin A exhibits potent efficacy against clinically relevant multidrug-resistant *Enterococcus* strains. Notably, aglomycin A acts synergistically with linezolid in both *in vitro* and *in vivo* models. Given that linezolid treatment is associated with serious adverse effects, such as myelosuppression, lactic acidosis, and neuropathies^58^, and has a narrow therapeutic window^59^, the combination of aglomycin A and linezolid might represent a promising therapeutic strategy against multidrug-resistant enterococcal infections.

Overall, our study demonstrates how integrating computational methods, particularly deep learning, with metabolomics can significantly accelerate the discovery of natural products. The DeepHalo workflow enables rapid, systematic mining of halogenated compounds from complex LC-HRMS datasets with markedly higher efficiency than traditional approaches. As a proof of concept, this strategy led to the identification of a novel antibiotic aglomycin A, which features rare structural building blocks, unique biosynthetic logic, and synergistic antibacterial activity with linezolid. We envision that this approach will boost the discovery of halogenated “dark matter” by efficiently analyzing vast LC-HRMS/MS datasets. Importantly, the DeepHalo framework is readily adaptable for discovering other classes of rare-element-containing metabolites. Beyond halogenation, similar computational strategies can be deployed to mine bioactive compounds with diverse ecological and pharmacological functions, thereby substantially expanding the current toolkit for natural product research.

## ASSOCIATED CONTENT

### Supporting Information

The Supporting Information is available free of charge at Experimental details for materials, methods, and spectroscopic data. AUTHOR INFORMATION

### Author Contributions

X.Y., H.B., and L.X. conceptualized and managed this study. Q.X. established DeepHalo workflow. C.S., W.M., H.X., H.N., C.M., W.J., D.Y., C.X., L.Q., and G.Q. carried out the wet experiments. Q.X., C.S., W.M., H.X., D.Y., W.S., and L.Y. analyzed the data. Q.X., C.S., W.M., H.X., and X.Y. drafted the manuscript. W.S., L.X., H.B. and X.Y. edited the manuscript.

### Notes

The authors declare no competing financial interest.

## ACKNOWLEDGMENT

This research was co-funded by CAMS Innovation Fund for Medical Sciences (CIFMS, 2021-I2M-1028) and the National Natural Science Foundation of China (81973219 and 82104047) and supported by Biomedical High Performance Computing Platform, Chinese Academy of Medical Sciences. We sincerely thank Professor Zhengyan Guo and Professor Xinyi Yang for their thorough review of our manuscript and constructive suggestions.

