## Supporting Information for "DeepHalo: Deep Learning-Powered Exploration of Halogenated Metabolites Uncovering Antibacterial Depsipeptides"

#### Comparison of available tools for targeted analysis of halogenates

Table S1. Comparison of available tools for targeted analysis of halogenates

| Software | Release Year | Maximum Batch Size / Mode | Speed[a] (~/sample) | MS/MS Support | Halogen Prediction Model | False Positive Rate (FPR) [c] |
|---|---|---|---|---|---|---|
| ChloroDBPFinder[1] | 2024 | Unlimited / Fully Automated | 16 min | Yes | Random Forest | 5.49% |
| HaloSeeker[2] | 2022 | Dozens / Semi-Automated | 10 min | No | Rule-Based | 13.14% |
| SIRIUS[3] | 2019 | Dozens / Semi-Automated | Days[b] | Yes | Deep Neural Network | 1.77% |
| MeHaloCoA[4] | 2016 | Dozens / Semi-Automated | 20 min | No | Rule-Based | 12.10% |
| DCAnalysis[5] | 2016 | Single Sample / Semi-Automated | 20 min | No | Polynomial Equation | NS |
| **DeepHalo** | 2025 | Unlimited / Fully Automated | 20 s | Yes | Deep Neural Network (IsoNN) | 0.08% |

Based on actual testing [a] or estimation [b] on the INST_SCM standard dataset developed in this project. [c] False positive rate of the halogen prediction models used in specific softwares, evaluated on IsoBase_Val_SN_Eva dataset. NS: Not Suitable for large-scale analysis.

#### General experiments

General Experimental Procedure. Optical rotations were recorded at 25 °C using a Rudolph Autopol® IV automatic polarimeter in MeOH. UV spectra were measured on a SHIMADZU UV-1800 spectrophotometer with MeOH as the solvent. For NMR spectra, both 1D and 2D were acquired at 600 MHz for $^{1}H$ NMR and 150 MHz for $^{13}C$ NMR using a Varian VNS-600 spectrometer in DMSO-$d_6$ ($\delta$ in ppm, $J$ in Hz). HRESIMS data collection was performed on a Waters Xevo G2-XS QTof mass spectrometer, operated by MassLynx v 4.1 software. Microporous adsorption resin (Diaion HP20; Mitsubishi, Japan) was utilized for column chromatography. Flash chromatography was carried out on a Combi Flash® Rf system equipped with an ODS flash column (RediSep Rf $C_{18}$ flash column, 12 g). Semi-preparative HPLC experiments were executed on an Agilent1200 series HPLC system with a diode array detector (DAD), employing a YMC-Pack $C_8$ semi-preparative column (10 mm × 250 mm, 5 $\mu$m).

### Establishment of DeepHalo

#### Datasets and preprocessing

*Datasets for model building*

For the training and evaluation of the models of Element Prediction (EPM) and Anormal isotope pattern Detection (ADM), we use simulated isotope spectra based on chemical formulas of the molecules from six compound databases: COCONUT[6], NPAtlas V2021_08[7], ChEBI[8], HMDB5.0[9], DSSTox[10], ZINC20[11] (Figure S1a). All databases were pooled together and filtered by the following rules: (1) the duplications of molecular formula were excluded; (2) only the compounds with molecular weight (MW) between 50 and 2000 Da were retained; (3) the compound contains elements from CHONFPSClBrINaBFeSe (Figure S2) and contains at least three hydrogen and one carbon atoms and at most four sulfur atoms. The molecules were divided into two primary groups: halo and non-halo. The halo group comprises three subcategories: Cl-type ($Cl/Cl_2$), Br-type ($Br/Cl_3$), and X-type (mixed/polyhalogenated). The non-halo group is further divided into four subcategories: B-type (B), Se-type (Se), Fe-type (Fe), and C-type (all remaining molecules). To balance the data, the extremely high numbers of molecules between 350 and 500 was done 1/2 random sampling. For augmenting the data with high molecular weights, dimerization strategies used for molecular with MW 500 – 1000 Da in Cl-, Br-, and C-type classes (Figure S1b). Specifically, C-type molecules were directly dimerized. In contrast, Cl-type molecules underwent dimerization followed by the replacement of chlorine with hydrogen (-Cl + H), while Br-type molecules were dimerized and then modified by substituting either bromine (-Br + H) or chlorine (-Cl + H) with hydrogen depending on their composition. Additionally, due to the limited number of B-, Se-, and Fe-containing molecules in the obtained datasets, we generated additional synthetic molecules, specifically, [M+Se], [M+B-3H], and [M+Fe-3H], based on CHON-only molecules (M) extracted from the COCONUT and NPAtlas natural products databases (Figure S1c–d). Furthermore, to simulate the false isotope patterns arising from co-eluting dehydro isomers, we employed the same molecular formulas from our B-, Se-, and Fe-type data augmentation and generated composite spectra by computing a weighted sum of a molecule's isotope pattern (Iso1) and that of its dehydro isomer (Iso2):

$$\text{Overlapping Iso} = \frac{\text{Iso1} + \text{ratio} \cdot \text{Iso2}}{1 + \text{ratio}}$$

where ratio represents the relative contribution of the dehydro isomer (0.33, 0.66, 0.99, 1.32, 1.65, 2.3, and 5 used in this study). For each isotope peak, calculate the new mass and fraction as follows:

$$m_{new} = (f1 \cdot m1 + f2 \cdot m2 \cdot ratio)/(f1 + f2 \cdot ratio)$$

$$f_{new} = f1 + f2 \cdot ratio$$

where, f1 and m1 represent the fraction abundance and mass of a given isotopic peak in Iso1, while f2 and m2 correspond to those in Iso2 with the same nominal mass (with absent peaks treated as f = 0, m = 0). Finally, peaks with relative intensities below 0.0001 are removed, and the remaining intensities are normalized to 100. These newly generated fake isotope patterns were then classified into the artifact-type class.

In total, 1,093,965 isotopic patterns were generated (the IsoBase dataset) and used to build EPM. From IsoBase, data corresponding to halo-class and C-type molecules were extracted, and empirically high noise was introduced to the mass and the relative intensity of each isotope peak, as previously described[12], to create the IsoHN dataset for ADM construction. The resulting datasets were then split into training and validation sets with an 80:20 ratio for model training. Additionally, the IsoHN dataset was used to statistically determine mass difference thresholds for validating the extracted isotope patterns. Subsequently, empirically standard noise was applied to IsoBase_Val, the validation dataset for EPM training, as previously described[12], to generate IsoBase_Val_SN for model evaluation during hyperparameter optimization. Finally, to simulate real-world scenarios and enable a fair comparison with other methods, we retained all halogenated data and the non-halogenated C-type subset while downsampling the remaining non-halogenated types to 0.1% of their original counts in the IsoBase_Val_SN dataset. Additionally, molecules with masses over 1000 Da were reduced by half in the same dataset. This resulted in the IsoBase_Val_SN_Eva dataset, used for comparative evaluation of the EPM against other models.

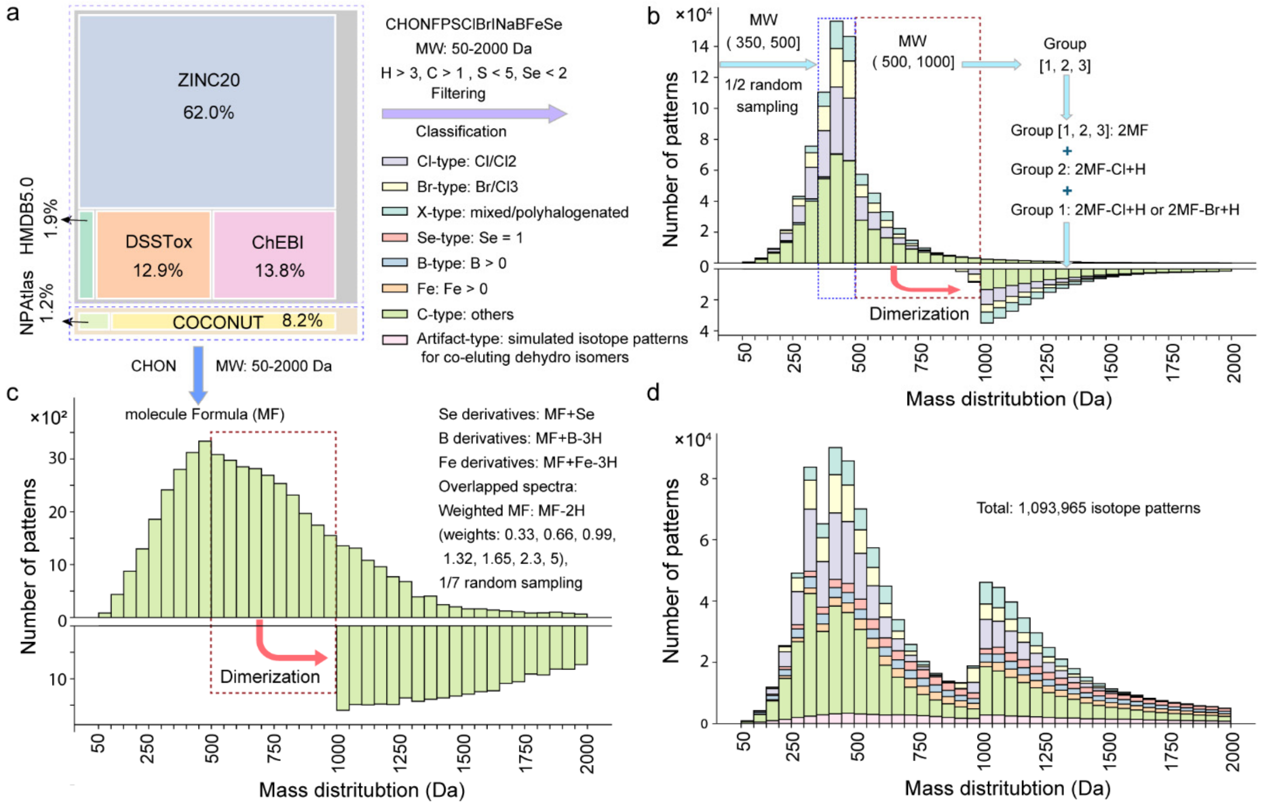

**Figure S1. Generation of the dataset for model training and evaluation**. **a**, Five public datasets were combined, dereplicated by formula, and filtered by molecular weight and elemental composition. The molecules were divided into two primary groups, halogenated (with three subcategories: Cl-, Br, and X-type, where X-type molecules contain Cl or Br atoms but that are not classified as Cl-type or Br-type.) and non-halogenated (with four subcategories: Se-, B-, Fe-, C-, and Artifact-type). **b**, Data was balanced by 1/2 random sampling, and isotope patterns with higher molecular weights were augmented using class-specific dimerization. **c**, Se-, B-, and Fe-type data were augmented, and simulated patterns from co-eluting dehydro isomers were generated based on the COCONUT and NPAtlas databases. **d**, A total of 1,093,965 patterns was generated for model training, along with their mass distribution.

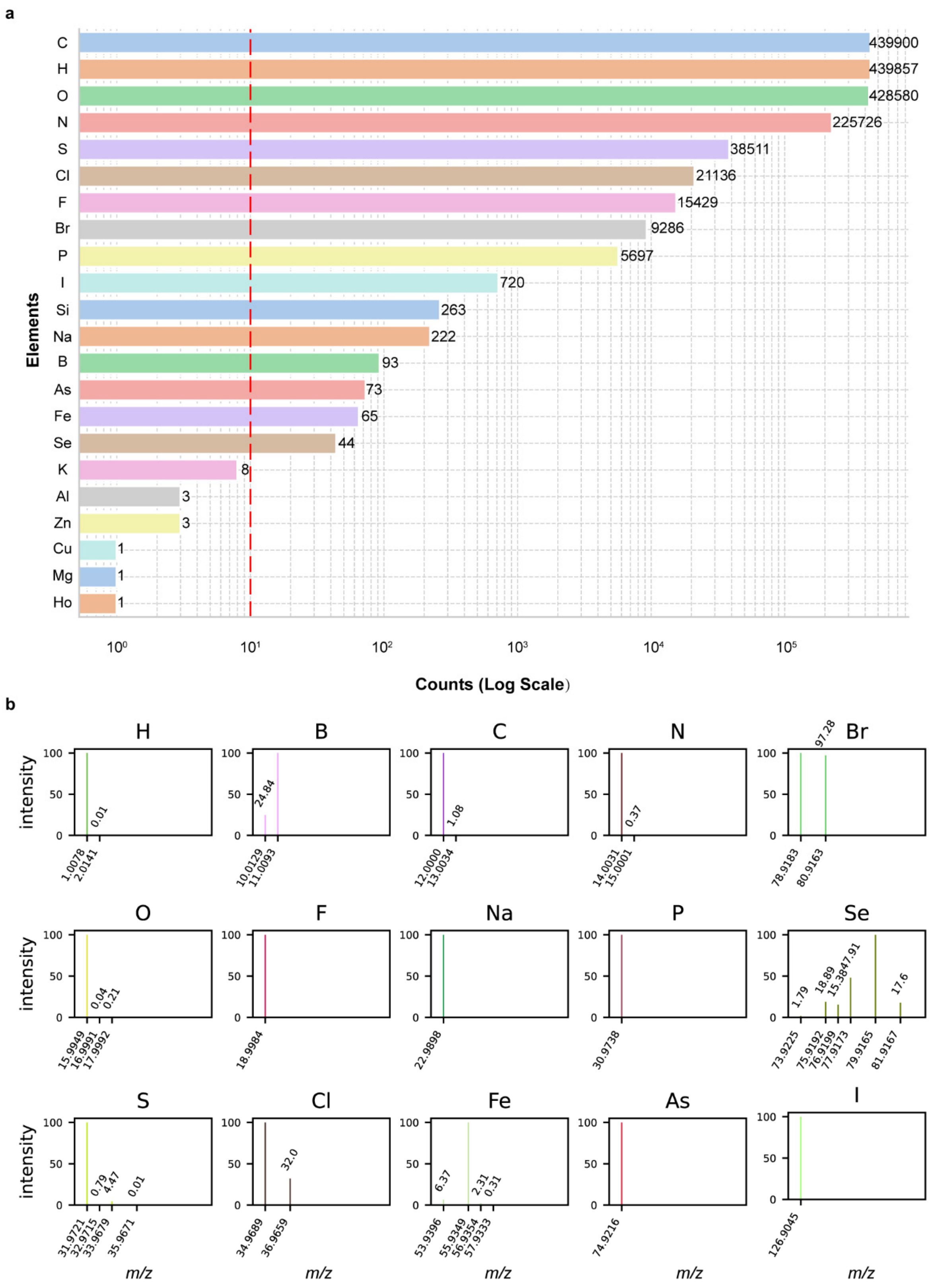

**Figure S2. Element selection for DeepHalo's Element Prediction Model (EPM). a**, Element occurrence statistics in natural products databases (COCONUT and NPAtlas). EPM development included all elements present in more than 10 molecules, excluding silicon (Si) due to its prevalence in chemically derivatized forms. **b**, Isotopic patterns of the included elements (Si excluded). In addition to bromine (Br) and chlorine (Cl), elements with distinct isotopic signatures, boron (B), selenium (Se), and iron (Fe), are grouped separately in the EPM.

*Real-world dataset for EPM model evaluation*

To rigorously evaluate the EPM under complex real-world conditions, we utilized the CASMI_Myxo_Plus dataset (Supplementary Data). This dataset integrates high-resolution isotope patterns from the *myxo* dataset [12](88 patterns acquired on a Bruker MaXis 2G qTOF spectrometer) and the CASMI 2016 dataset[13-14] (366 positive and 166 negative patterns measured on a Q Exactive Plus Orbitrap), totaling 619 isotope patterns (mass range: 67.042 – 2213.962 Da) featuring chlorine, boron, and bromine-containing compounds (with the only selenium compound excluded due to large mass error. To further challenge the model, we augmented the dataset with four edge cases: two iron adducts, one overloaded sample, and one overlapped dehydro isomers.

*Simulated dataset for deepHalo workflow benchmarking*

Gold standard datasets with verified ground truths are crucial for benchmarking algorithms. However, real-world LC-MS datasets often fall short of this ideal due to contaminants and the inherent uncertainty in instrument noise and accuracy. To circumvent these issues, we employed simulated metabolomic data with precise ground truth. Specifically, we generated simulated LC-MS datasets using SMITER_modified (https://github.com/xieyying/SMITER_modified), a customized version of the LC-MS simulator from the original SMITER tool[15]. These datasets were based on a published collection of 1820 compounds[16], including 459 halogen-containing compounds (395 with either one or two chlorine atoms, 38 with either one bromine or three chlorine atoms, and 26 with either one bromine and one chlorine, two to four bromine, or six chlorine atoms).

First, ten random input files for SMITER_modified were generated. In these files, peak widths were randomly assigned between 10 and 15 seconds, and scan start times were calculated as the documented retention times (in seconds) minus 0.3 times the respective peak width. Peak scaling factors, representing the maximum intensity of each compound, were randomly set between 1e5 and $5e^8$. Using these parameters, along with various resolutions, mass accuracies, and a scan time of 0.3 seconds (typical for QTOF mass spectrometry), a series of crowded LC-MS chromatograms spanning 1 to 13 minutes were simulated. Additionally, 1,000 random noise signals with intensities between 0 and 1e5 were introduced to each dataset to mimic real-world scenarios. The simulated datasets include conditions with fixed mass accuracy (1 ppm) and varying resolutions (from $1 \times 10^4$ to $6 \times 10^4$), as well as conditions with fixed resolution ($2 \times 10^4$) and varying mass accuracies (2 – 5 ppm). Each condition comprised ten replicate LC-MS files to form SM1820-Base. From these simulations, ten replicate LC-MS data with a mass accuracy of 1 ppm and a resolution of $2 \times 10^4$ were selected and combined to form the SM1820-R20K dataset, which was used to benchmark DeepHalo against other methods.

*Measured dataset for DeepHalo workflow benchmarking*

To evaluate the effectiveness of DeepHalo in identifying chlorine- or bromine-containing compounds in real-world complex samples, a series of LC-HRMS/MS analyses were acquired using UPLC-QTof (See below). A standard sample containing 11 purified compounds (5 with one or two chlorine atoms and 2 with a bromine atom), with masses ranging from 238.0509 to 1447.4300 Da, was analyzed at low, medium, and high concentrations using UPLC-HRMS/MS. Each concentration was analyzed with one to three replicates, resulting in 12 LC-HRMS runs, collectively referred to as the IN-house STandards dataset (INST). Additionally, to assess DeepHalo's capability in detecting halogenated compounds within complex samples, spiked sample analyses were performed. Varying amounts of the standard sample were mixed with F1 and M3 culture media (Supplementary Information) and analyzed via UPLC-HRMS/MS, generating the IN-house STandards Spiked in Culture Media dataset (INST_SCM). All resulting data were converted to .mzML format, making them suitable for DeepHalo analysis.

The details of INST and INST_SCM acquisition are as follows: the stock solution of halogenated standards included demeclocycline (0.35 mg), sansanmycin A (0.69 mg), erythromycin (0.35 mg), [4-Br-Phe]-sansanmycin (4.09 mg), [4-Cl-Phe]-sansanmycin (5.54 mg), bleomycin (2.99 mg), vancomycin (0.43 mg), 6-Cl-L-tryptophan (0.43 mg), and 5-Br-L-tryptophan (0.43 mg) dissolved in 6 mL 50% methanol. Then the stock solution was diluted with 50% methanol to 10-fold and 100-fold. The preparation of spiking solution was same as the method described above, except that F1 (2% glucose, 1% corn steep liquor, 0.4% soybean meal, 1% dextrin, 0.5% peptone, 0.2% $(NH_4)_2SO_4$, pH 5.6) and M3 (2% soluble starch, 0.5% glycerol, 1% defatted wheat germ, 0.3% meat extract, 0.3% dry yeast, and 0.3% $CaCO_3$, pH 7.0) media were used as solvents for dilution, respectively. After centrifugation, the above solutions were analyzed using a Waters ACQUITY UPLC H-Class system (ACQUITY UPLC BEH™ $C_{18}$ column, 1.7 $\mu$m, 2.1 × 100 mm, 0.3 mL/min) coupled with a Waters Xevo G2-XS QTof mass spectrometer. The samples of halogenated standards solution and the spiking solution (20 $\mu$L) were analyzed by UPLC with a linear gradient of MeCN-$H_2O$ containing 0.1% formic acid (A/B = 95/5−65/35, *v*/*v*, over 9 min; followed by A/B = 65/35−0/100, *v*/*v*, over 2 min). The fast DDA function in continuum mode were utilized for mass analysis. The specific instrumental parameters were as follows: source temperature:120 °C, cone gas flow: 30 L/h, desolvation temperature: 450 °C, desolvation gas flow: 800 L/h, capillary voltage: 3 kV, and sample cone voltage: 40 V. The reference was LE (leucine-enkephalin, 2 ng/mL), and it was sampled every 60 seconds. The Waters Masslynx (V4.1) was used to acquire and correct data. The MS survey scan covered a range of 150 - 2000 *m/z* with a scan time of 0.5 s, while the MS/MS scan ranged from 40 to 2000 *m/z* with a scan time of 0.1 s. Following the MS scan, the three ions with the highest intensity were selected sequentially for MS/MS analysis. Ramping the collision energy occurred in two stages: from 30 to 65 eV for low-mass analytes (150 Da), and from 55 to 90 eV for high-mass analytes (2000 Da).

Additionally, we also evaluated the efficiency of DeepHalo by apply it to the dataset of the Critical Assessment of Small Molecule Identification 2022 contest (CASMI 2022)[17]. This dataset consists of 145 Orbitrap LC-MS/MS data in both ESI(+) and ESI(-) mode for 500 unique compounds, with 37 precursor adducts bearing chlorine or bromine.

*Data availability*

The simulated isotope patterns derived from six open compound databases: COCONUT https://coconut.naturalproducts.net/. Version January 2022), NPAtlas (https://www.npatlas.org/. V2022_09), ChEBI (https://www.ebi.ac.uk/chebi/. Accessed 1 May 2023), HMDB 5.0 (https://hmdb.ca/downloads, Accessed 30 May 2023), DSSTox (https://figshare.com/search?q=CFM-ID_metadata_DTXCID. Accessed 25 May 2023), ZINC20 (https://zinc.docking.org/). The real isotope patterns are from open datasets: the *myxo* dataset (https://bio.informatik.uni-jena.de/software/sirius/. consisting of 88 isotope patterns measured on a Bruker MaXis 2G qTOF spectrometer). The CASMI 2016 dataset (http://casmi-contest.org/2016/).

The real LC-HRMS dataset CASMI 2022 (https://fiehnlab.ucdavis.edu/casmi), INST (MassIVE MSV000098203), and INST_SCM (MassIVE MSV000098204).

**Construction of deepHalo**

The DeepHalo framework was implemented in Python using TensorFlow, Keras, pyOpenMS, and KD-tree. It consists of four main components: a dataset processing module, a deep learning module, an LC-MS data analysis module, and a post-processing module.

The dataset processing module handles raw input data and generates training features by calculating isotope patterns from molecular formulas based on the above datasets and preprocessing methods. For each compound, the first six isotope peaks (p0-p5) are extracted. The relative intensities of these peaks, normalized to their maximum value, along with the normalized monoisotopic mass (p0) and the mass differences Δm1 (p1-p0) and Δm2 (p2-p1), are computed and prepared as input features for model training.

The deep learning module constructs two deep neural network (DNN) models: one for element prediction (EPM) and another for anomalous isotope pattern detection (ADM). DeepHalo employs isotope patterns to identify the presence of chlorine and bromine in molecules. To achieve this, we developed the Isotope Pattern Neural Network (IsoNN) architecture and trained an Element Prediction Model (EPM) using approximately 109 million simulated isotope patterns. IsoNN is a dual-branched deep neural network that processes two distinct types of input data: the relative intensities and masses of isotope peaks (including monoisotopic mass and mass differences Δm1/Δm2). The intensity branch incorporates Gaussian noise injection for regularization, followed by dense layer transformations, while the mass feature branch applies a nonlinear scaling operation, Gaussian noise injection, and dense processing. The processed features from both branches are concatenated and passed through a feed-forward network, culminating in classification probabilities via a softmax output layer. Model performance was evaluated on both simulated (IsoBase_Val_SN_Eva) and real (CASMI_Myxo_Plus) datasets using standard metrics such as Precision, Recall, F1-score, and False Positive Rate (FPR), with comparisons to alternative methods. The training process and hyperparameter optimization details are provided in the Supporting Information. Following the construction of the EPM, an ADM was developed for isotope peak validation based on their intensities. Following the construction of the EPM, an ADM was developed for isotope peak validation based on their intensities. Unlike the EPM, the ADM employs a deep autoencoder neural network trained on the IsoHN dataset. It uses the feature embeddings (output from the first dense layer of the relative intensity branch in the EPM) as input, leveraging its pre-trained representations to enhance anomaly sensitivity. The network uses ReLU activation functions, two hidden layers (dimensions: [64,16]), optimized by minimizing MSE loss via Adam (learning rate: 0.0003). After training, the reconstruction error was computed on the IsoHN dataset. The anomaly threshold was determined by locating the inflection point of the reconstruction error threshold (Q_τ) versus percentile (τ) curve, defined as:

$$t_{\text{threshold}} = Q_{\tau^*}, \qquad \tau^* = \arg\max_{\tau} \left| \frac{\mathrm{d}^2 Q_\tau}{\mathrm{d}\tau^2} \right|$$

where $Q_\tau$ is the $\tau$-th percentile of reconstruction errors, and the second derivative is approximated via discrete differencing.

The LC-MS data analysis module is the core component of DeepHalo and performs four key functions: (1) isotope pattern detection and validation from raw LC-MS data, (2) halogen presence prediction, (3) comprehensive scoring, and (4) MS1/MS2 data output. For isotope pattern extraction, we developed a robust pyOpenMS-based solution, selected for its superior performance and Python compatibility. The customized workflow enables dual-level isotopic pattern extraction, simultaneously characterizing overall distributions at the centroid feature level while capturing individual patterns at the scan level. To further improve the confidence of feature detection, we established a two-dimension validation process for centroid isotope patterns. The first validation stage examines mass differences using strict criteria derived from the IsoHN dataset (0.05%-95.5% quantile ranges for Δm1 and Δm2). Specifically, starting with the first isotope peak in a detected cluster, three consecutive peaks that meet the mass difference conditions are identified as the first three isotope peaks. The second validation stage employs the trained ADM to filter isotopes based on their relative intensity patterns. Validated isotopic features are then classified using the optimized EPM. We further introduce a halogen confidence score (H-score) that integrates evidence from both centroid and scan levels to comprehensively evaluate halogen presence. The H-score is defined as:

$$H_{score} = \frac{1}{3}(ZZ_{score} + R_{halo} + C_{score})$$

where $ZZ_{score}$ represents the consistency of halo-positive across consecutive scans, calculated using a modified ZigZag index[18]:

$$ZZ_{score} = \left(1 - \frac{\sum_{i=1}^{N-2}(2S_i - S_{i-1} - S_{i+1})^2}{4N-8}\right) R_{halo}$$

$R_{halo}$ represents the proportion of halogen-positive scans:

$$R_{halo} = \frac{\sum_{i=1}^{N} S_i}{N}$$

$C_{score}$ is binary indicator (1/0) based on the prediction of the centroid isotope pattern of the feature. $S_i$ represents the EPM prediction result for scan *i* (1 for halogen presence, 0 for absence), and $N$ is the total number of scans in a feature. Subsequently, the H-score serve as a critical filter, with only features meeting the threshold proceeding to $MS^1$ data export (CSV format). For LC-MS data acquired in DDA mode, the corresponding $MS^2$ spectra of filtered features can be optionally exported via a customized mzml2gnps package.

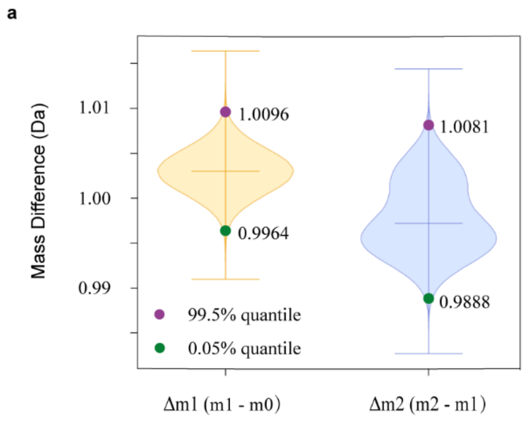

**Figure S3.** Mass difference statistics for Δm1 and Δm2 derived from the IsoHN dataset (validation criteria: 0.05%–95.5% quantile ranges)

Finally, the post-processing module is responsible for dereplication and writing DeepHalo results to GNPS output files. If a user-supplied chemistry database is available, the pipeline will perform $MS^1$-based dereplication. Candidate compounds are identified by matching precursor ions ($[M+H]^+$/$[M+Na]^+$) within a user-defined mass accuracy, detecting halogen signatures (Cl/Br count ≥1), and comparing isotopic patterns using cosine similarity between experimental and theoretical intensities for further filtering with a user-defined threshold (default: 0.96). The cosine similarity is defined as:

$$\text{cosine\_similarity} = \frac{\sum_{i=1}^{5} I_{\text{exp}_i} \cdot I_{\text{thre}_i}}{\sqrt{\sum_{i=1}^{5}\left(I_{\text{exp}_i}\right)^2} \cdot \sqrt{\sum_{i=1}^{5}\left(I_{\text{thre}_i}\right)^2}}$$

where: $I_{\exp_i}$ represents the experimental isotopic peak intensity of the (i)-th isotopic peak. $I_{\text{thre}_i}$ represents the theoretical isotopic peak intensity of the (i)-th isotope. Additionally, if molecular networking is performed via GNPS platform[19], the pipeline also writes the prediction and $MS^1$-based dereplication results into the network file output by GNPS, which subsequently can be visualized using Cytoscape[20].

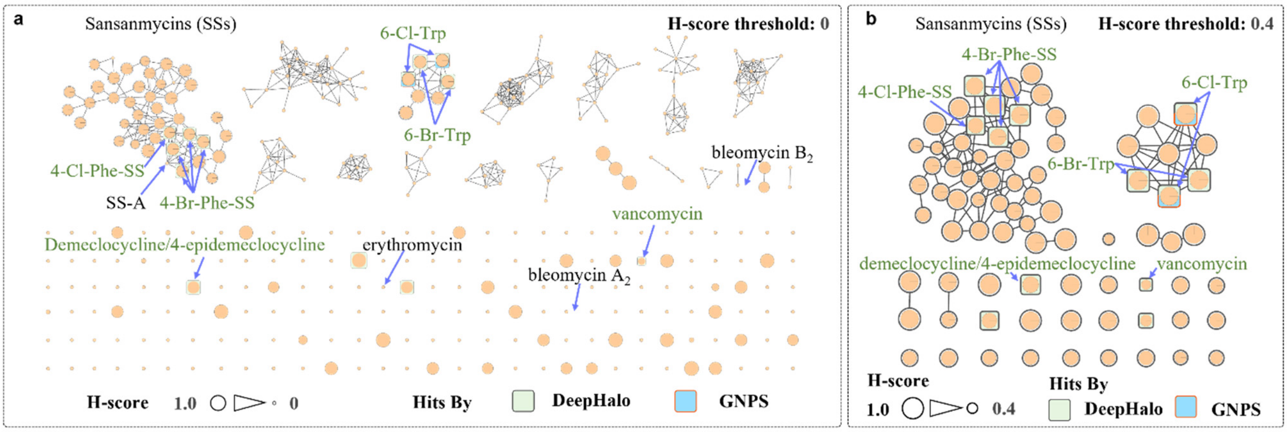

**Figure S4. DeepHalo's analysis, combined with an integrated H-score, enables halogenate dereplication and annotation in complex matrices.** (**a**) With an H-score threshold of 0 (all features retained), all seven spiked halogenated standards (highlighted in green) were automatically annotated through molecular networking with NPAtlas/INST_SCM database, while the four non-halogenated standards (displayed in blank) were manually identified, exhibiting low H-score values. (**b**) Applying an H-score threshold of 0.4 selectively retained halogenated features, significantly streamlining downstream analysis.

We showcased DeepHalo's ability to dereplicate and annotate halogenated compounds using our in-house dataset (INST_SCM) with integrated H-score analysis (Figure S4). By setting the H-score threshold to 0, DeepHalo output all detected features along with their corresponding MS2 spectra. Subsequent molecular networking, combined with automatic dereplication function of DeepHalo using the NPAtlas microbial natural products database supplemented by the INST_SCM standards, enabled automatic annotation of all seven halogenated standards spiked into two complex culture media (F1 and M3) with perfect recall@1. The only exception was vancomycin, which was matched alongside its isomer (chloro-orienticin B). Furthermore, all four spiked non-halogenated standards were manually identified and, as expected, exhibited low H-scores. The visualization aided by the H-score enabled differentiation of compounds based on their likelihood of containing halogens, facilitating targeted labeling of halogenated compounds in complex datasets (Figure S4a). By contrast, applying an H-score threshold of 0.4 exclusively retained potential halogenated features (Figure S4b). This selective filtering not only greatly streamlines downstream analysis, but also dramatically cuts the time needed for subsequent molecular networking, making DeepHalo particularly effective for large-scale MS data.

**Hyperparameter optimization**

The hyperparameters of the deep learning model were optimized using a combination of Keras Tuner's Bayesian optimization and manual tunning. The automatically search space included the following parameters: batch sizes (4, 16, 64, 128), mass difference feature scaling power (0, 10, 20), number of units in the first dense layer for each input type (16, 64, 256 for intensity features; 8, 32, 128, 512 for mass features), number of additional dense layers (1 to 5), number of units in these additional layers (32, 128, 512), dropout rate (0.0, 0.3), and learning rate (0.0001, 0.0003). The model was compiled using the Adam optimizer and the sparse categorical cross-entropy loss function, which is appropriate for multi-class classification tasks. Model performance was evaluated based on validation accuracy, and the best model was selected after 200 trials. The manually tunned parameters include Gaussian noise levels (N1 for mass and N2 intensity features), and the input features. First, five rounds of hyperparameter searches based on Bayesian Optimization were conducted to evaluate the impact of Gaussian noise levels and the number of isotopic peaks on model performance. The Gaussian noise levels (N1 and N2) tested were 0.0005 and 0.02, 0.001 and 0.03, and 0.001 and 0.04 for the first five isotopic peaks, as well as 0.001 and 0.03 for the first four and six isotopic peaks (Figure S5b), respectively. The input features included the intensities of the above defined isotopic peaks and two mass differences ($\Delta m1$ and $\Delta m1$). Other hyperparameters were optimized automatically by Bayesian Optimization. Additionally, a search was conducted using Gaussian noise level of 0.001(N1) and 0.03 (N2), incorporating five isotopic peak intensities, mass differences, $\Delta m1$ and $\Delta m2$, as well as monoisotopic mass (m0) as input features to evaluate the impact of monoisotopic mass on model performance. Model performance was evaluated on the IsoBase_Val_SN dataset using two metrics: the area under the micro-average precision-recall curve (AUPRC) and the area under the micro-average ROC curve (AUROC) for halogenated class. The hyperparameters was optimized on a server with 12 cores and 1 NVIDIA A100 GPU.

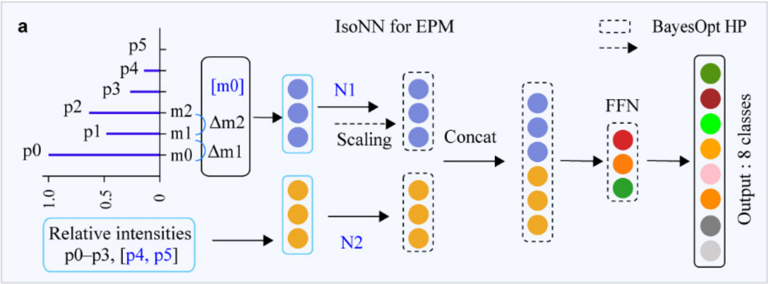

b

| Hyperparameter (HP) | Model 1 | Model 2 | Model 3 | Model 4 | Model 5 | Model 6 |
|---|---|---|---|---|---|---|
| Mass Gaussian Noise (N1) | 0.001 | 0.001 | 0.001 | 0.001 | 0.0005 | 0.001 |
| Intensity Gaussian Noise (N2) | 0.03 | 0.03 | 0.03 | 0.03 | 0.02 | 0.04 |
| Relative Intensities (p0–p3, [p4, p5]) | p0–p4 | p0–p3 | p0–p5 | p0–p4 | p0–p4 | p0–p4 |
| Include monoisotope mass (m0)? | × | × | × | √ | × | × |

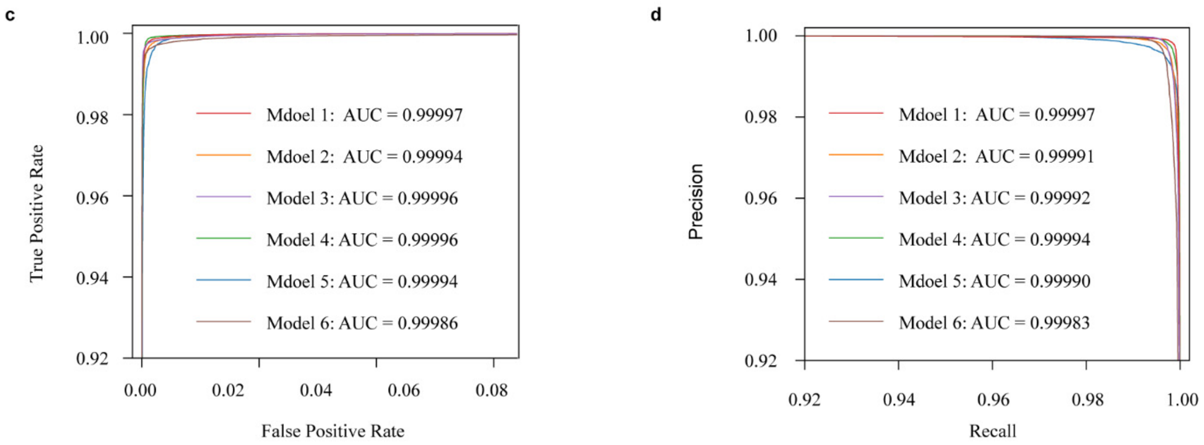

**Figure S5. Hyperparameter optimization**. **a**, Network architectures (IsoNN for EPM) along with the manual hyperparameters highlighted in blue, and **b**, Six EPM variants with different manually tuned hyperparameters. **c**–**d**, Evaluation of the EPM models for binary classification of halogenated versus non-halogenated compounds, using AUROC (**c**) and AUPRC d) metrics. All achieved AUROCs and AUPRCs > 0.9998, with Model 1 performing best (AUROC = 0.99997; AUPRC = 0.99997).

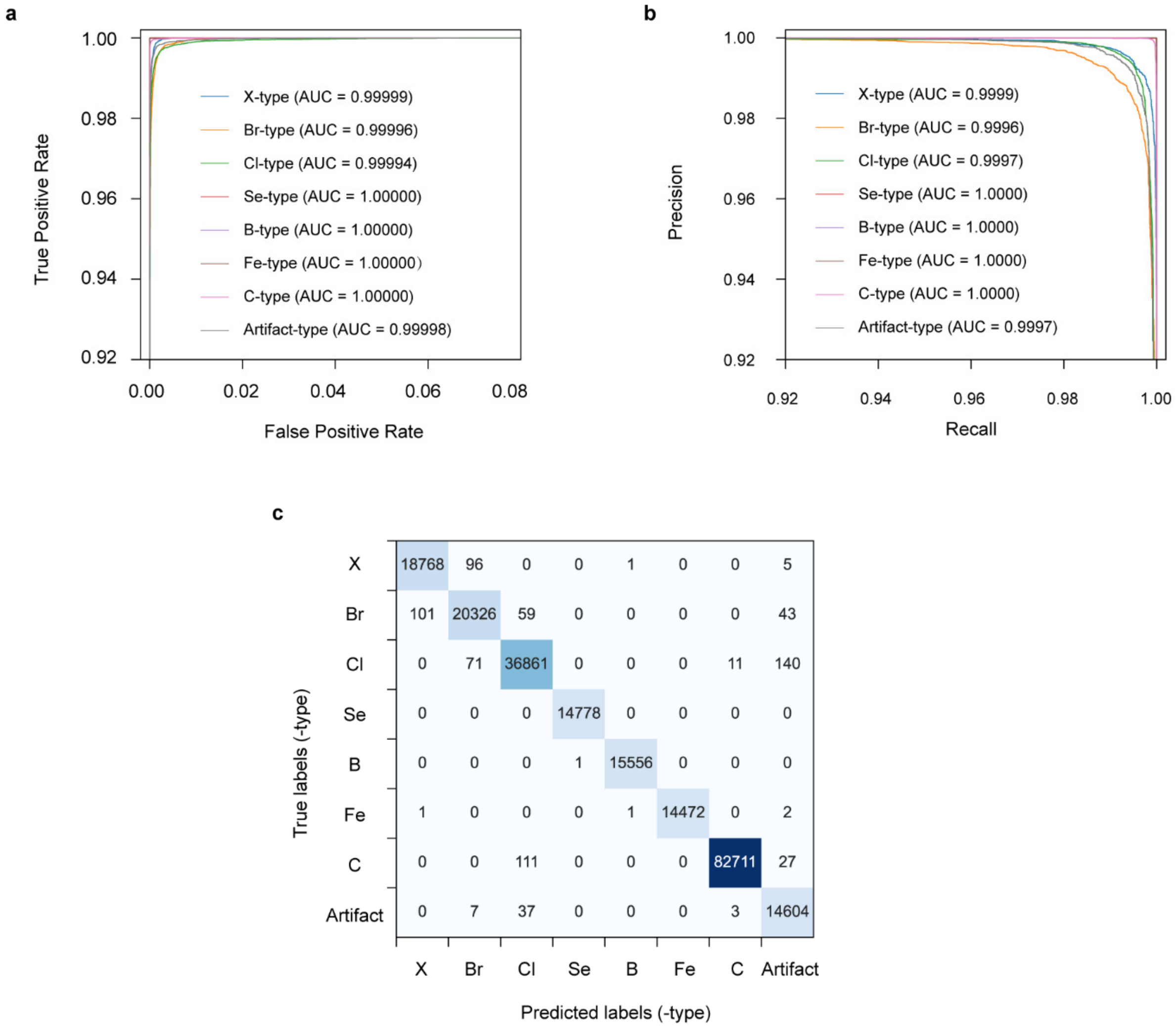

**Figure S6. Classification capability of EPM for all eight classes. a**–**b**, The performance of the EPM is evaluated across all eight classes using AUROC (**a**) and AUPRC (**b**) metrics, along with a confusion matrix (**c**) based on a simulated validation dataset (IsoBase_Val_SN_Eva). All classes achieved high AUROC and AUPRC values, with the Se, B, Fe, and C classes attaining perfect scores (1.0000). While the Cl-type class performed the worst due to some misclassifications with the C-type and Artifact-type, it is still reach high AUROC (0.99994) and AUPRC values (0.9997).

**Comparison of EPM performance with other methods.**

The performance of EPM was compared with SIRIUS 4.0 (DNNRegressionPredictor), HaloSeeker v2.0.3.3, MeHaloCoA v0.99.0, ChloroDBPFinder v0.1.0 (binary model), and DCAnalysis v1.10 on both simulated and real isotope patterns. DCAnalysis was excluded from simulated data evaluation due to its inability to process large datasets. For MeHaloCoA, compounds were classified as halogenated when meeting all of the following reported criteria[4]: 1) Isotope peak intensity ratios: $0.30 < p_2/p_0 \leq 6.00$ and $p_1/p_0 < 1.50$; 2) Mass difference constraints: $(m/z_1 - m/z_0) \geq (1.003 - \varepsilon)$ and $(m/z_2 - m/z_0) \leq (1.997 + \varepsilon)$; where p for peak intensity, *m/z* for mass-to-charge, and $\varepsilon = 0.05$ Da. HaloSeeker identified halogenated compounds according to published criteria[2], classifying a spectrum as positive when any of these conditions was satisfied:1) $(A\text{-}2/A = 0) \cap (A\text{+}2/A \geq 0.25)$; 2) $(A\text{-}2/A \geq 0.60) \cap (A\text{+}2/A \geq 0.20)$; 3) $(A\text{-}2/A \geq 0.27) \cap (A\text{+}2/A \geq 0.36)$; where A represents intensity of the most intense peak in the pattern.

**Performance evaluation of DeepHalo workflow**

We systematically evaluated the DeepHalo workflow using both simulated and real-world datasets. For the simulated data (SM1820-Base and SM1820-R20K) with complete ground truth, we defined true positives (TP) as correctly identified halogenates, false positives (FP) as both misclassified non-halogenates and artificial features predicted as positive, false negatives (FN) as undetected or misclassified halogenates, and true negatives (TN) as correctly classified non-halogenates (excluding false features). For three real-world LC-HRMS datasets, metrics were computed on curated subsets with confirmed ground truth (n = 132 for INST, n = 264 for INST_SCM, and n = 500 for CASMI 2022), where TP represented correctly identified validated halogenated features, FP denoted misclassified validated non-halogenates, FN included undetected or misclassified validated halogenates, and TN comprised properly classified validated non-halogenates. From these values, we computed recall, precision, F1-score, and false positive rate (FPR) using the standard formulas provided in the scikit-learn documentation. ChloroDBPFinder was not included in the SM1820-R20K analysis due to its inability to automatically process LC–MS data without tandem mass spectrometry. Additionally, due to the extremely long running time of competing methods (> 12 h), only DeepHalo was evaluated on the CASMI 2022 dataset (n = 145 LC-HRMS data). The evaluations were performance on a Windows 10 system equipped with a 12th Gen Intel Core i7-12700F processor, running at 2.10 GHz with 12 cores, and 64 GB of RAM, accelerated by NVIDIA RTX A4500.

**Code availability**

The DeepHalo can be found at https://github.com/xieyying/deephalo and https://pypi.org/project/deephalo/ ; The mzml2gnps have been integrated into DeepHalo and also can be found at https://github.com/xieyying/mzml2gnps and https://pypi.org/project/mzml2gnps/ ; The modified SMITER can be found at https://github.com/xieyying/SMITER_modified.

#### DeepHalo assisted-mining of halogenated metabolites

##### Actinomycete Material and Identification

Fifty-four actinomycete strains (53 *Streptomyces* spp. and *Micromonospora* sp.) were isolated from lichen, plant, and soil samples collected in Tibet, China. All strains were taxonomically identified by 16S rRNA gene sequencing (GenBank accession No. See Table S3).

##### Miniaturized cultivation of 53 *Streptomyces* spp. and one *Monomicrospora* sp.

The strains stored at -80°C were inoculated onto MS2 medium (Table S4) and incubated statically at 28°C for 7 days. The strains were then transferred onto fresh MS2 medium for subculturing. A sterile iron spatula was used to pick a 0.5 $cm^2$ piece of mycelium, which was then transferred to an Erlenmeyer flask (250 mL) containing 50 mL of M2 medium. The flask was shaken at 220 rpm and 28°C for 48 hours to obtain the seed culture.
Miniature fermentation was conducted in a 96-well plate, with 800 $\mu$L of medium (M1-M13 and MS1-MS13, as shown in Table S4) added to each well. After adding seed culture (40 $\mu$L) to each well, the plates were sealed and incubated at 28°C for 7 days. The liquid culture media were shaken at 900 rpm, while the solid culture media were incubated statically. The blank media were treated in parallel as a control.

After freeze-drying the liquid fermentation broth for 48 hours, methanol (800 $\mu$L) was added to each well for extraction by ultrasonication (5 min) and shaking (30 min), followed by centrifugation (4500 rpm, 10 min); For the solid fermentation broth, methanol-ethyl acetate mixture (1:4, 800 $\mu$L) was added to each well for extraction by ultrasonication (5 min) and shaking (30 min). Then the supernatant was transferred to the new 96-well plate, allowed the solvent to evaporate, and redissolved in methanol (300 $\mu$L). Finally, the solution was centrifuged at 4500 rpm for 10 minutes. An ODS plate was used to adsorb 600 $\mu$L of the above supernatant, which was then eluted with 300 $\mu$L of methanol, and the effluent was collected in a deep 96-well plate. An additional 800 $\mu$L of methanol was used to further elute the ODS plate, and the combined eluate was collected in the deep

96-well plate for UPLC-MS analysis and the construction of a metabolic database.

Table S2. Information of 54 strains used in this study and the culture media used.

| No. | Strains | GenBank accession numbers | Media |
|---|---|---|---|
| 1 | *Strepomyces* sp. cmx-5-11 | PV655549 | M1-M12; MS1-MS12 |
| 2 | *Strepomyces* sp. cmx-11-25 | PV655550 | M1-M12; MS1-MS12 |
| 3 | *Strepomyces* sp. cmx-6-16 | PV655551 | M1-M12; MS1-MS12 |
| 4 | *Strepomyces* sp. cmx-11-4 | PV655552 | M1-M12; MS1-MS12 |
| 5 | *Strepomyces* sp. cmx-067-1 | PV655553 | M1-M12; MS1-MS12 |
| 6 | *Strepomyces* sp. cmx-10-25 | PV655554 | M1-M12; MS1-MS12 |
| 7 | *Strepomyces* sp. cmx-058-1L | PV655555 | M1-M12; MS1-MS12 |
| 8 | *Strepomyces* sp. X-19 | PV655556 | M1-M12; MS1-MS12 |
| 9 | *Strepomyces* sp. cmx-13-2 | PV655557 | M1-M12; MS1-MS12 |
| 10 | *Strepomyces* sp. cmx-4-9 | PV655558 | M1-M12; MS1-MS12 |
| 11 | *Strepomyces* sp. cmx-18-6 | PV655559 | M1-M12; MS1-MS12 |
| 12 | *Strepomyces* sp. A30 | PV655560 | M1-M12; MS1-MS12 |
| 13 | *Strepomyces* sp. 5-10 | PV655561 | M1-M12; MS1-MS12 |
| 14 | *Strepomyces* sp. 021-3 | PV655562 | M1-M12; MS1-MS12 |
| 15 | *Strepomyces* sp. cmx-5-18 | PV655563 | M1-M12; MS1-MS12 |
| 16 | *Strepomyces* sp. X-45 | PV655564 | M1-M12; MS1-MS12 |
| 17 | *Strepomyces* sp. cmx-13-9 | PV655565 | M1-M12; MS1-MS12 |
| 18 | *Strepomyces* sp. cmx-8-6 | PV655566 | M1-M12; MS1-MS12 |
| 19 | *Strepomyces* sp. 049-1 | PV655567 | M1-M12; MS1-MS12 |
| 20 | *Strepomyces* sp. 029-5 | PV655568 | M1-M12; MS1-MS12 |
| 21 | *Strepomyces* sp. cmx-14-8 | PV655569 | M1-M12; MS1-MS12 |
| 22 | *Strepomyces* sp. 021-4 | PV655570 | M1-M12; MS1-MS12 |
| 23 | *Strepomyces* sp. cmx-10-8 | PV655571 | M1-M12; MS1-MS12 |
| 24 | *Strepomyces* sp. cmx-10-37 | PV655572 | M1-M12; MS1-MS12 |
| 25 | *Strepomyces* sp. 061-3 | PV655573 | M1-M12; MS1-MS12 |
| 26 | *Strepomyces* sp. cmx-4-25 | PV655574 | M1-M12; MS1-MS12 |
| 27 | *Strepomyces* sp. cmx-4-7 | PV655575 | M1-M12; MS1-MS12 |
| 28 | *Strepomyces* sp. X-80 | PV655576 | M1-M12; MS1-MS12 |
| 29 | *Strepomyces* sp. cmx-8-16 | PV655577 | M1-M12; MS1-MS12 |
| 30 | *Strepomyces* sp. 039-1 | PV655578 | M1-M12; MS1-MS12 |
| 31 | *Strepomyces* sp. 020-2-3H-GM | PV655579 | M1-M12; MS1-MS12 |
| 32 | *Strepomyces* sp. C19 | PV655580 | M1-M12; MS1-MS12 |
| 33 | *Strepomyces* sp. cmx-10-19 | PV655581 | M1-M12; MS1-MS12 |
| 34 | *Strepomyces* sp. cmx-11-23 | PV655582 | M1-M12; MS1-MS12 |
| 35 | *Strepomyces* sp. cmx-10-20 | PV655583 | M1-M12; MS1-MS12 |
| 36 | *Strepomyces* sp. 030-hv | PV655584 | M1-M12; MS1-MS12 |
| 37 | *Strepomyces* sp. cmx-10-17 | PV655585 | M1-M12; MS1-MS12 |
| 38 | *Strepomyces* sp. cmx-11-39 | PV655586 | M1-M12; MS1-MS12 |
| 39 | *Streptomyces* sp. XZ19-198 | PV668804 | M1-M11, M13; MS1-MS11, MS13 |
| 40 | *Streptomyces* sp. XZ-19-091 | MW110675 | M1-M11, M13; MS1-MS11, MS13 |
| 41 | *Streptomyces* sp. XZ-19-435 | PV688581 | M1-M11, M13; MS1-MS11, MS13 |
| 42 | *Streptomyces* sp. XZ-19-081 | MW110672 | M1-M11, M13; MS1-MS11, MS13 |
| 43 | *Streptomyces* sp. XZ-19-043-3 | PV668805 | M1-M11, M13; MS1-MS11, MS13 |
| 44 | *Streptomyces* sp. XZ-19-147 | MW110693 | M1-M11, M13; MS1-MS11, MS13 |
| 45 | *Streptomyces* sp. XZ-19-152 | MW110695 | M1-M11, M13; MS1-MS11, MS13 |
| 46 | *Streptomyces* sp. XZ-19-459 | PV668806 | M1-M11, M13; MS1-MS11, MS13 |
| 47 | *Streptomyces* sp. XZ-19-316 | MW110733 | M1-M11, M13; MS1-MS11, MS13 |
| 48 | *Streptomyces* sp. XZ-20-671 | PV668802 | M1-M11, M13; MS1-MS11, MS13 |
| 49 | *Streptomyces* sp. XZ-19-034 | MW110661 | M1-M11, M13; MS1-MS11, MS13 |
| 50 | *Streptomyces* sp. XZ-19-513 | PV668801 | M1-M11, M13; MS1-MS11, MS13 |
| 51 | *Streptomyces* sp. XZ-19-136 | MW110690 | M1-M11, M13; MS1-MS11, MS13 |
| 52 | *Streptomyces* sp. XZ-19-259 | MW110722 | M1-M11, M13; MS1-MS11, MS13 |
| 53 | *Streptomyces* sp. XZ-19-359 | MW110747 | M1-M11, M13; MS1-MS11, MS13 |
| 54 | *Micromonospora* sp. XZ-19-293 | MW110725 | M1-M11, M13; MS1-MS11, MS13 |

Table S3. Thirteen Media used in OSMAC.

| Media | Components |
|---|---|
| M1 | potato 200 g, glucose 20 g, 1 L of deionized water, natural pH |
| M2 | yeast extract 4.0 g, glucose 4.0 g, malt extract 10.0 g, 1 L of deionized water, pH 7.2 |
| M3 | soluble starch 20.0 g, glycerol 5.0 g, malt extract 10.0 g, beef extract 3.0 g, yeast extract 3.0 g, $CaCO_3$ 3.0g, 1 L of deionized water, pH 7.0 |
| M4 | galactose 3.3g, dextrin 3.3g, glycerol 1.7g, soybean peptone 1.7g, corn steep liquor 0.83g, $(NH_4)_2SO_4$ 0.33g, $CaCO_3$ 2.0g, 1L deionized water, pH 7.0 |
| M5 | glucose 10.0 g, beef extract 10.0 g, peptone 1.0 g, NaCl 5.0 g, 1 L deionized water, pH 7.0 |
| M6 | glucose 20.0 g, regular starch 5.0 g, peptone 6.0 g, $(NH_4)_2SO_4$ 7.0 g, $CaCO_3$ 2.0 g, 1L deionized water, natural pH |
| M7 | regular starch 20.0 g, glucose 20.0 g, peptone 3.0 g, beef extract 3.0 g, $CaCO_3$ 2.5 g, trace elements 1 mL (trace element composition per 0.1 L: $FeSO_4$ 0.1 g, $MnCl_2$ 0.1 g, $ZnSO_4$ 0.1 g, $CuSO_4$ 0.1 g, $CoCl_2$ 0.1 g), peanut cake powder 10.0 g, 1 L deionized water, pH 7.2 |
| M8 | mannitol 40.0 g, malt extract powder 40.0 g, yeast extract powder 10.0 g, $K_2HPO_4$ 2.0 g, $MgSO_4·7H_2O$ 0.5 g, $FeSO_4·7H_2O$ 0.01 g, 1 L deionized water, pH 7.2 |
| M9 | $K_2HPO_4$ 1.0 g, $NaNO_3$ 0.3 g, KCl 0.005 g, $MgSO_4·7H_2O$ 0.005 g, $FeSO_4$ 0.001 g, sucrose 30.0 g, 1 L deionized water, pH 7.0 |
| M10 | glycerol 20.0 g, molasses 10.0 g, casein peptone 5.0 g, $CaCO_3$ 4.0 g, peptone 1.0 g, 1 L deionized water, natural pH |
| M11 | glucose 20.0 g, malt extract powder 40.0 g, yeast extract powder 4.0 g, $K_2HPO_4$ 5.0 g, NaCl 2.5 g, $ZnSO_4$ 0.04 g, $CaCO_3$ 0.4 g, 1 L deionized water, pH 6.0 |
| M12 | soybean peptone 3.0 g, NaCl 5.0 g, pancreatin peptone 17.0 g, $K_2HPO_4$ 2.5 g, glucose 2.5 g, 1 L deionized water, natural pH |
| M13 | yeast extract powder 4.0 g, malt extract powder 10.0 g, soluble starch 4.0 g, natural pH |

The addition of 2% agar powder to the above liquid medium results in solid media MS1-MS13.

**High-throughput mining of halogenated metabolites**

Fifty-three *Streptomyces* and one *Micromonospora* strains (Table S3) were preprocessed, and analyzed using UPLC-HRMS/MS as previously described[21]. Including 60 blank culture media, a total of 1356 UPLC-HRMS/MS data were collected. The raw data were converted to .mzML format using MSconvert and automatically analyzed with DeepHalo's 'detect' function. The minimum peak intensity was set to 1000, signal-to-noise ratio to 6, minimum intensity to $1e^5$, and H-score threshold to 0.4, with feature validation through mass difference and isotopic peak intensities. The analysis was performed using the same computer configuration as previously employed for DeepHalo evaluation, with the complete process requiring about 4.9 hours. The output $MS^2$ data from DeepHalo were uploaded to the GNPS server for analysis using classic molecular networking with a precursor and fragment *m/z* tolerance of 0.02, a cosine score of 0.6, and a minimum of 5 fragments, with other parameters set to default. After that, the dereplication function of DeepHalo was used to dereplicate based on NPAtlas (bacterial-derived data only) and MIBiG database, then write the predictions and dereplication results to the GNPS output network file. The final results were visualized using Cytoscape V3.8[20].

Table S4. The 16 strains with halogenated features identified by DeepHalo

| No. | Strains | Media | Confirmed compounds | Compound class |
|---|---|---|---|---|
| 1 | *Strepomyces* sp. A30 | M3-M5, M8-M10, M12; MS2, MS3, MS5, MS7, MS8, MS12 | svetamycins A-C | depsipeptide |
| 2 | *Strepomyces* sp. cmx-6-16 | M8, M12 | colibrimycin A1 | lipopeptide |
| 3 | *Strepomyces* sp. 029-5 | MS10 | salinamide B | bicyclic depsipeptide |
| 4 | *Strepomyces* sp. x-80 | M1, M2, M4, M5, M10, M11; MS1-MS7, MS10-MS12 | lysolipin I; benzastatin C | aromatic polyketide; benzastatin derivative |
| 5 | *Strepomyces* sp. cmx-11-23 | M2-M4, M6-M10; MS2-MS12 | - | - |
| 6 | *Micromonospora* sp. XZ-19-293 | M2-M4, M6, M7, M11, M12; MS2-MS10, MS12 | - | - |
| 7 | *Streptomyces* sp. XZ-19-259 | MS1-MS4, MS8 | - | - |
| 8 | *Strepomyces* sp. 020-2-3H-GM | MS10 | - | - |
| 9 | *Strepomyces* sp. cmx-10-8 | MS3 | - | - |
| 10 | *Strepomyces* sp. cmx-4-7 | MS11 | lysolipin I; | aromatic polyketide |
| 11 | *Strepomyces* sp. cmx-5-18 | M7 | - | - |
| 12 | *Strepomyces* sp. X-19 | M6 | - | - |
| 13 | *Streptomyces* sp. XZ-19-435 | MS4 | - | - |
| 14 | *Streptomyces* sp. XZ-19-034 | M11, MS3 | - | - |
| 15 | *Streptomyces* sp. XZ-19-359 | M2-M5, M10, M11, M13; MS1, MS2, MS3, MS7 | - | - |
| 16 | *Streptomyces* sp. XZ-19-459 | M2, M4, M5, M7, M13 | - | - |

**Structural elucidation of known halogenated metabolites**

Structural confirmation of five classes of DeepHalo hits. Further validation of five groups of known halogenates was performed using an integrated approach combining HRMS, MS/MS spectra, UV spectroscopy, and BGC comparison analysis (Figure S8-11) [22-26]. Gene cluster comparison: comparison and visualization of gene clusters were used clinker, a Python based tool[27]. The GenBank files of biosynthetic gene cluster and similar clusters were uploaded. The gene cluster comparison figures were analyzed with minimum alignment sequence identity of 0.3 and the number of decimals of 2.

Svetamycin A (Group 2 from strain A30), with an $[M+H]^+$ ion at *m/z* 631.2224 and an $[M+Na]^+$ ion at *m/z* 653.2059, had a molecular formula of $C_{24}H_{35}ClN_8O_{10}$ deduced from the (+)-HRESIMS and characteristic isotope distributions of the chlorine atom. Careful analysis of its $MS^2$ spectra revealed that it showed fragment ions at *m/z* 74.060, 85.076, 115.051, 119.035, 160.052, 173.054, 214.083, 272.098 and 302.100 (Figure S8), corresponding to diagnostic ions of *α*-MeSer, Pip, (*β*, *γ*-OHdPip), γ-ClPip, Haa-(*α*-MeSer), [(*β*, *γ*-OHdPip)-Haa]-CO, (*β*, *γ*-OHdPip)-Ala, Haa-(*α*-MeSer)-Pip, [Ala-(γ-ClPip)-Pip]-CO, respectively. Based on similarity analysis with known biosynthetic gene clusters, the *sve* BGC of A30 showed high similarity to the svetamycin BGC (*sve*)[28], it was confirmed as known compound, svetamycin A. The similarity of the obtained $MS^2$ spectra was analyzed in the molecular network of group 6, and provided evidence for the presence of the known compounds svetamycin B and svetamycin C.

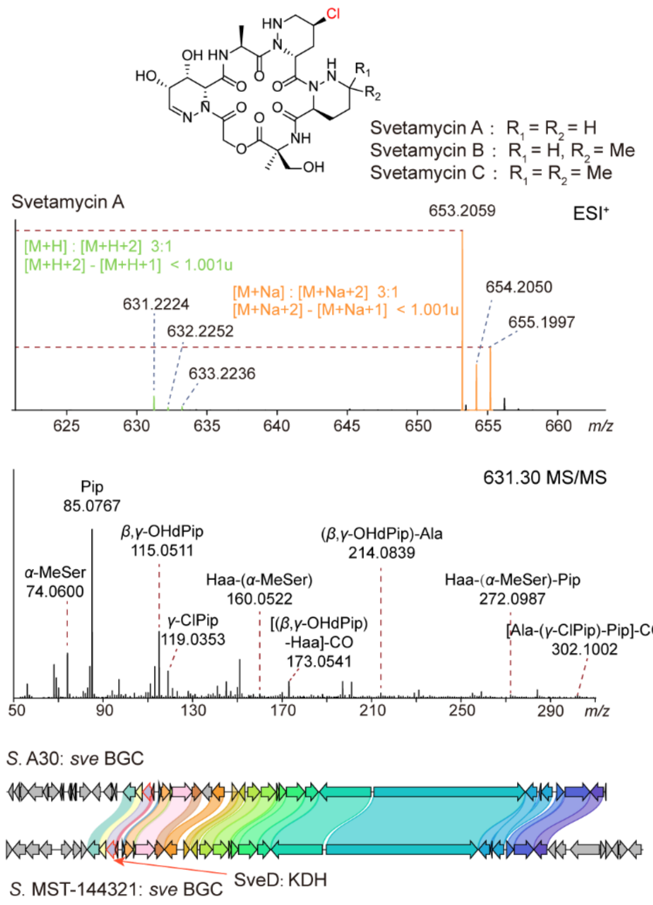

**Figure S7. Verification the group of svetamycins annotated by DeepHalo using HRMS, $MS^2$ spectra and bioinformatics.**

Colibrimycin A1, categorized as group 3 from *Streptomyces* sp. 6-16, was detected as an [M-$H_2O$+H]$^+$ ion at *m/z* 570.1794 and an [M+Na]$^+$ ion at *m/z* 610.1710. Its molecular formula, $C_{27}H_{30}ClN_5O_8$, was deduced from (+)-HRESIMS, supported by the characteristic isotopic pattern associated with the presence of a chlorine atom. Careful analysis of its MS$^2$ spectra revealed that it showed fragment ions at *m/z* 121.064, 191.036, 207.112 and 219.031 (Figure S9), corresponding to diagnostic ions of $C_8H_9O^+$, $C_{10}H_8ClN_2^+$, $C_{11}H_{15}N_2O_2^+$ and $C_{11}H_8ClN_2O^+$, respectively. Based on similarity analysis with known biosynthetic gene clusters, *cbm* BGC showed significant similarity to the colibrimycins BGC (*cbm*) from *Streptomyces* sp. CS147[23].

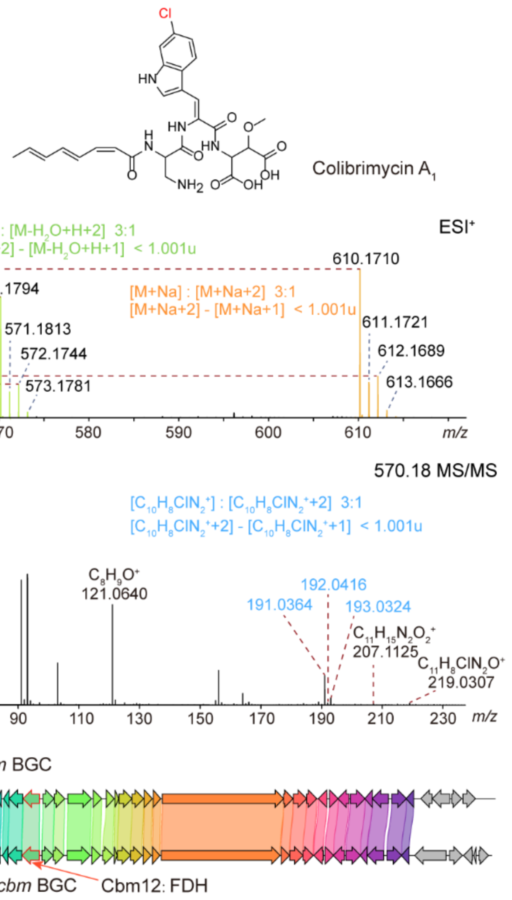

**Figure S8. Verification the group of colibrimycins annotated by DeepHalo using HRMS, MS$^2$ spectra and bioinformatics.**

The salinamide B (Group 3 from strain 29-5, Figure S10), with an $[M+Na]^+$ ion at *m/z* 1078.4480, had a molecular formula of $C_{51}H_{70}ClN_7O_{15}$ deduced from the (+)-HRESIMS and characteristic isotope distributions of the chlorine atom. Comparative analysis of biosynthetic gene clusters revealed that the *sln* BGC shares high sequence similarity with the salinamides BGC (*sln*) from *Streptomyces* sp. CNB0915[24].

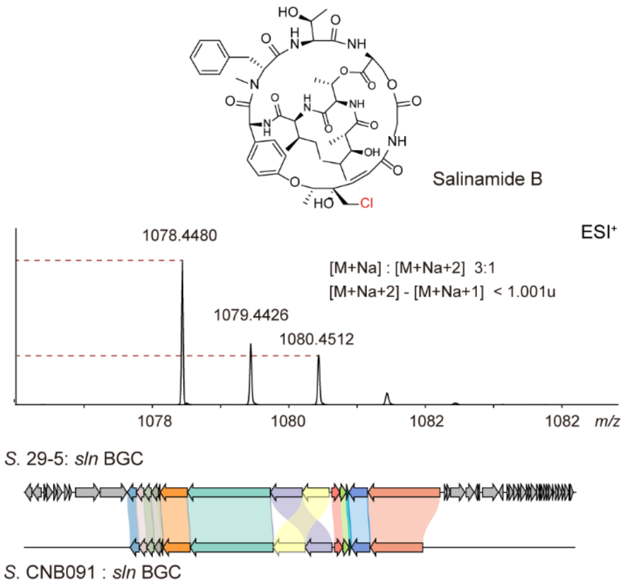

**Figure S9. Verification the group of salinamides annotated by DeepHalo using HRMS and bioinformatics.**

Both Groups 4 (lysolipins) and 5 (benzastatins) were derived from *Streptomyces* sp. X80. Lysolipin I was detected as an $[M+H]^+$ ion at *m/z* 598.1168 and an $[M+Na]^+$ ion at *m/z* 620.0966, with its molecular formula ($C_{29}H_{24}ClNO_{11}$) determined by (+)-HRESIMS, supported by the characteristic chlorine isotopic pattern (Figure S11). Its identity was further confirmed through UV spectroscopy and comparative analysis with known biosynthetic gene clusters[25]. Benzastatin C, characterized by an $[M+H]^+$ ion at *m/z* 351.1832, was assigned the molecular formula $C_{19}H_{27}ClN_2O_2$ based on (+)-HRESIMS and isotopic signatures consistent with the presence of chlorine. Genomic analysis of the associated *bez* BGC revealed high sequence similarity to the benzastatins BGC (*bez*) previously identified in *Streptomyces* sp. CNB0916, suggesting a conserved biosynthetic pathway[26].

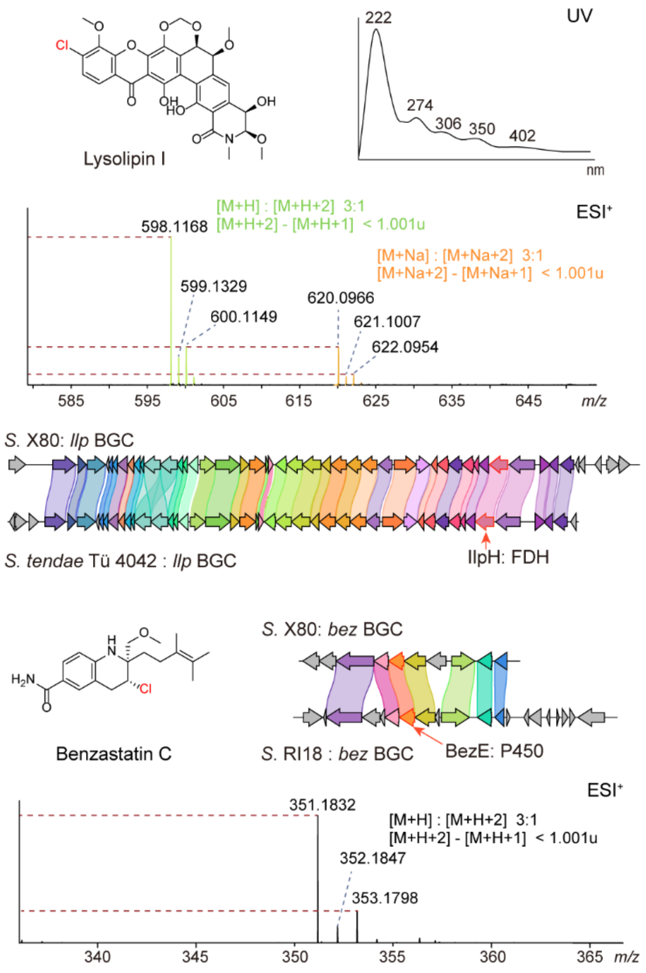

**Figure S10. Verification the groups of lysolipins and benzastatins annotated by DeepHalo using HRMS, UV and bioinformatics.**

**Investigation of halogenase distribution**

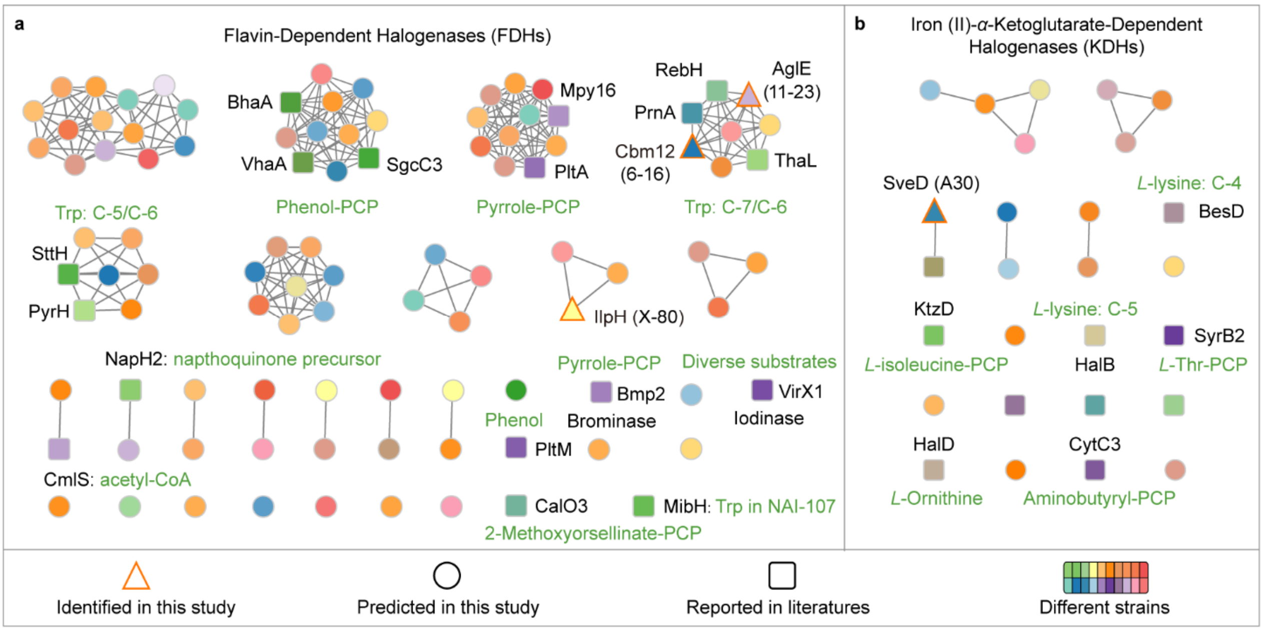

**Figure 11. Sequence similarity networks (SSNs) of predicted halogenases across our 53 Streptomyces strains.** (a) SSNs of flavin-dependent halogenases (FDHs). (b) SSNs of iron (II)/α-Ketoglutarate-Dependent Halogenases (KDHs). Candidate halogenases were detected via HMMer[29] and analyzed for sequence similarity using EFI-EST[30]. The substrates of characterized halogenases are highlighted in green.

The draft genomes of 53 *Streptomyces* strains were sequenced on an Illumina HiSeq platform (Illumina, San Diego, CA, USA) and assembled with SPAdes v3.13.1. Candidate halogenases were identified using HMMer 3.1b2. For FDHs, the queries included Pfam profiles PF04820 (Trp halogenase)[31] and PF22045 (Chloramphenicol halogenase, halogenating an alkyl group)[32] together with sequences of characterized FDH with diverse substrates (PrnA: Trp, C-7; RebH: Trp, C-7; PyrH: Trp, C-5; SttH: Trp, C-6; Thal: Trp, C-6; PltA: Pyrroly-PCP; Bmp2: Pyrroly-PCP; Mpy16: Pyrroly-PCP; NapH2: Napthoquinone precursor; PltM: diverse phenolic compounds; SgcC3: Tyrosyl-PCP; BhaA: β-hydroxy tyrosyl-PCP; VhaA Hexapeptide-peptideyl-PCP: ClmS: Acetyl-CoA, CalO3: 2-Methoxyorsellinate-PCP; VirX1: Iodinase, diverse substrates)[33], while for KDHs, the queries comprised Pfam profiles PF22814 (Carrier-protein-independent halogenase WelO5)[34] and PF23169 (Halogenase D)[35] along with reference KDH sequences (KthP: Piz-PCP, CytC3: Aminobutyryl-PCP, SyrB2: L-Thr-PCP; KtzD: L-isoleucine-PCP; BesD: L-lysine, C-4; HalB: L-lysine, C-5; and HalD: L-ornithine)[33]. A strict E-value cutoff of 1e$^{-20}$ was applied. The putative halogenases, together with known ones, were then analyzed for sequence similarity, and SS networks (SSNs) were constructed using EFI-EST[30]. The resulting SSNs were visualized with Cytoscape[20]. The amino acid sequences of the corresponding proteins for 53 in-house *Streptomyces* strains have been deposited GenBank and the accessions archived in Supplementary Data.

All 2110 *Streptomyces* genomic sequences (as of January 14, 2021) were downloaded from National Center for Biotechnology Information. Using the same method described above, potential halogenases were identified in these genomes, and their corresponding accessions have been archived in Supplementary Data.

#### Structural elucidation of aglomycins

##### Acquisition of aglomycins A and B

Three strains with potential halogenate production, based on DeepHalo analysis, were further cultured in 500 mL Erlenmeyer flasks and analyzed using UPLC-LCMS/MS and DeepHalo to confirm the presence of the target halogenates. While the selected three strains all produced the target compounds, only *Streptomyces* sp. 11-23 yielded enough for further study. Previous studies have shown that *N*-acetylglucosamine (GlcNAc) can enhance secondary metabolite yields[36], and that NaCl can boost the production of certain halogenated compounds[37-38]. Thus, various concentrations of GlcNAc and NaCl were supplemented to M3 media to test their effects on the production of major component, and the optimal media was used as culture media. The strain 11-23 was cultured on ISP2 plates at 28 °C for 7 days. For seed culture preparation, a mycelial agar plug (1 cm²) was inoculated into each of five 500 mL Erlenmeyer flasks containing 100 mL of ISP2 medium and incubated at 28 °C on a rotary shaker at 220 rpm for 2 days. For large-scale fermentation, 5 mL of the seed culture was added to each of two hundred 500 mL Erlenmeyer flasks containing 100 mL of culture medium and cultivated at room temperature for 9 days.

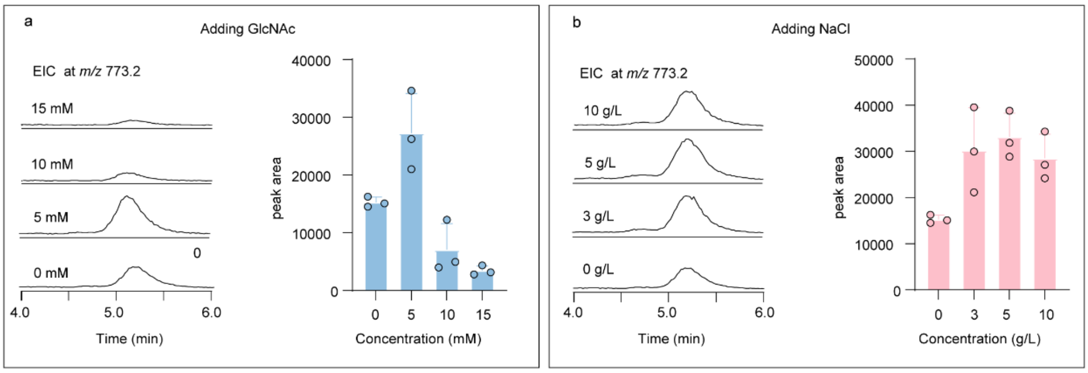

**Figure S12. Optimization of fermentation conditions for the production of aglomycin A.** Various concentrations of GlcNAc (a) and NaCl (b) were added to the M3 medium and tested to improve *Streptomyces* sp. 11-23 producing aglomycin A (n = 3). The results indicate a significant increase in aglomycin A production with 5 mM GlcNAc and 3 – 10 g/L NaCl.

The fermentation broth (19 L) was collected and adsorbed onto 1.5 L of microporous adsorption resin X1180. The column was sequentially rinsed with deionized water, and 30%, 50%, 70% and 100% ethanol–water (*v*/*v*) solutions. Each gradient was eluted until the effluent was colorless, resulting in fractions Fr.A, Fr.B, Fr.C, and Fr.D. Fr. D (260 mg) was further fractionated using reversed-phase $C_{18}$ flash column chromatography with a stepwise gradient of MeCN-$H_2O$, yielding five subfractions (Fr.D1-D5). Subfraction D3 (46 mg) was subjected to semipreparative HPLC using a YMC-Pack $C_8$ column, yielding aglomycin A (**1**) ($t_R$ 41.2 min; 7.5 mg). Fr. A and Fr. B was further fractionated using reversed-phase (RP) $C_{18}$ flash column chromatography, employing a stepwise gradient of MeCN-$H_2O$ (10:90 to 20:80, *v*/*v*, elution time 30 min; 20:80 to 50:50, elution time 30 min; 50:50 to 100:0, elution time 20 min; flow rate, 5 mL/min), resulting in five subfractions (Fr. $AB_1$-$AB_5$). Fr. $AB_3$ (52 mg) was subjected to semipreparative HPLC using a YMC-Pack $C_8$ column (10 mm × 250 mm, 5 $\mu$m; MeCN-Water, 42:58, *v*/*v*; flow rate, 2.5 mL/min), yielding aglomycin B (**2**) ($t_R$ 29.3 min; 2.9 mg).

#### Determination of the planar structure of aglomycin A (1)

Table S5. $^{1}H$ and $^{13}C$ NMR data for compound **1** in DMSO-$d_6$.

| | No. | $\delta_C$, type | $\delta_H$, multi. ($J$ in Hz) | $^{1}H$-$^{1}H$ COSY | HMBC |
|---|---|---|---|---|---|
| ACTA | 2 | 168.2, C | | | |
| | 4 | 148.7, C | | | |
| | 5 | 123.3, CH | 8.37, s | | 1'', 4, 2 |
| | 1' | 114.8, C | | | |
| | 2' | 142.1, C | | | |
| | 3' | 119.5, C | | | |
| | 4' | 131.4, CH | 7.49, d (9.0) | 5' | 2', 3', 6' |
| | 5' | 116.6, CH | 6.73, t (9.0) | 4', 6' | 1', 2', 3', 4', 6' |
| | 6' | 128.3, CH | 7.68, d (7.8) | 5' | 2, 1', 2', 3', 4' |
| | 1'' | 159.2, C | | | |
| | 2'-$NH_2$ | | 7.30, s | | 1', 3' |
| Thr[1] | 1 | 166.8, C | | | |
| | 2 | 52.5, CH | 4.71, m | 3, 2-NH | 1, 4, ACTA-1'' |
| | 3 | 70.3, CH | 4.72, m | 2, 4 | 1, 4, *N*-Me-Val-1 |
| | 4 | 18.2, $CH_3$ | 1.30, d (6.0) | 3 | 2, 3 |
| | 2-NH | | 8.17, d (9.0) | 2 | 1, ACTA-1'' |
| Pro[2] | 1 | 167.5, C | | | |
| | 2 | 53.5, CH | 4.88, dd (3.0, 7.8) | 3 | 3, 4 |
| | 3 | 29.3, $CH_2$ | 1.93, m | 2, 4 | 1, 2, 5, 4 |
| | | | 1.84, m | 2, 4 | 1, 5, 4 |
| | 4 | 25.1, $CH_2$ | 1.93, m | 3, 5 | 3, 5, 2, |
| | | | 2.33, m | 3, 5 | 3, 5, |
| | 5 | 46.7, $CH_2$ | 3.66, m | 4 | 3, 4, |
| | | | 3.61, m | 4 | 3, 4, |
| Piz[3] | 1 | 170.4, C | | | |
| | 2 | 57.9, CH | 5.10, d (6.0) | 3 | 1, 3, 4, Pro-1 |
| | 3 | 26.5, $CH_2$ | 1.74, m | 2, 4 | 1, 5 |
| | | | 2.09, d (13.8) | 2, 4 | 5 |
| | 4 | 21.0, $CH_2$ | 1.64, d (13.8) | 3, 5 | 2 |
| | | | 1.55, m | 3, 5 | 2 |
| | 5 | 46.6, $CH_2$ | 3.09, m | 4, 5-NH | 3, 4 |
| | | | 2.56, m | 4, 5-NH | 3, 4 |
| | 2-NH | | 6.78, m | 5 | 2, 4, 5, Pro-1 |
| epoxy-Val[4] | 1 | 167.0, C | | | |
| | 2 | 52.5, CH | 5.22, d (8.4) | 2-NH | 1, 3, 4, 5, Piz-1 |
| | 3 | 57.4, C | | | |
| | 4 | 49.6, $CH_2$ | 2.56, d (5.4) | | 3, 5 |
| | | | 2.51, d (5.4) | | 2, 3, 5 |
| | 5 | 20.5, $CH_3$ | 1.46, s | | 2, 3, 4 |
| | 2-NH | | 8.26, d (8.4) | 2 | 1, 2, Piz-1 |
| *N*-Me-Val[5] | 1 | 167.8, C | | | |
| | 2 | 69.3, CH | 3.28, d (9.6) | 3 | 1, 3, 4, 5, $NCH_3$, epoxy-Val-1 |
| | 3 | 27.0, CH | 2.32, m | 2, 4, 5 | 1, 2, 4, 5 |
| | 4 | 18.6, $CH_3$ | 0.76, d (6.6) | 3 | 2, 3, 5 |
| | 5 | 21.1, $CH_3$ | 0.94, d (6.6) | 3 | 2, 3, 4 |
| | $NCH_3$ | 39.4, $CH_3$ | 3.07, s | | 2, epoxy-Val-1 |

$^{1}H$ and $^{13}C$ NMR data were recorded at 600 and 150 MHz, respectively. The assignments were based on 2D NMR ($^{1}H$-$^{1}H$ COSY, HSQC, HMBC and ROESY) experiments.

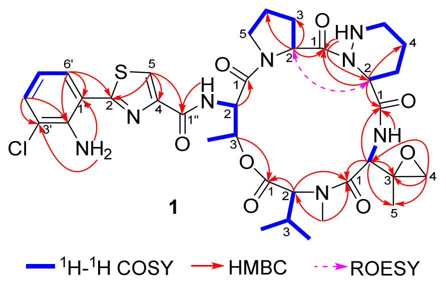

**Figure S13. The structure and the key 2D NMR correlations for aglomycin A (1).**

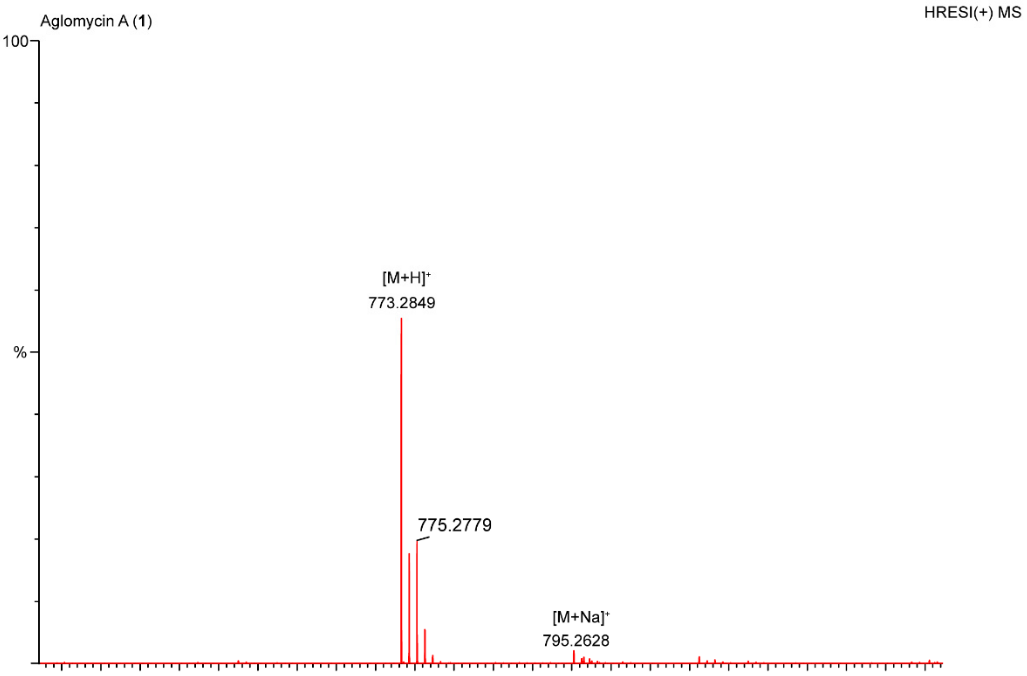

**Figure S14. HRESI(+)MS spectrum of compound 1**

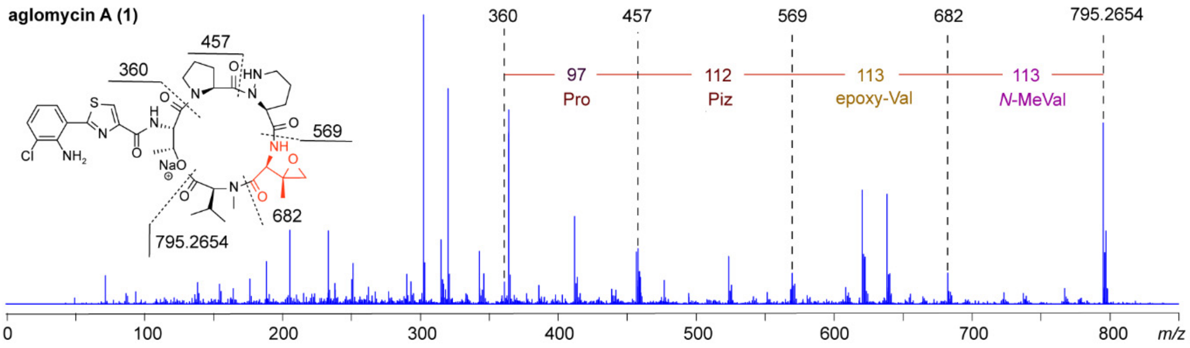

**Figure S15. MS$^2$ spectrum of sodium adduct ion of compound 1**

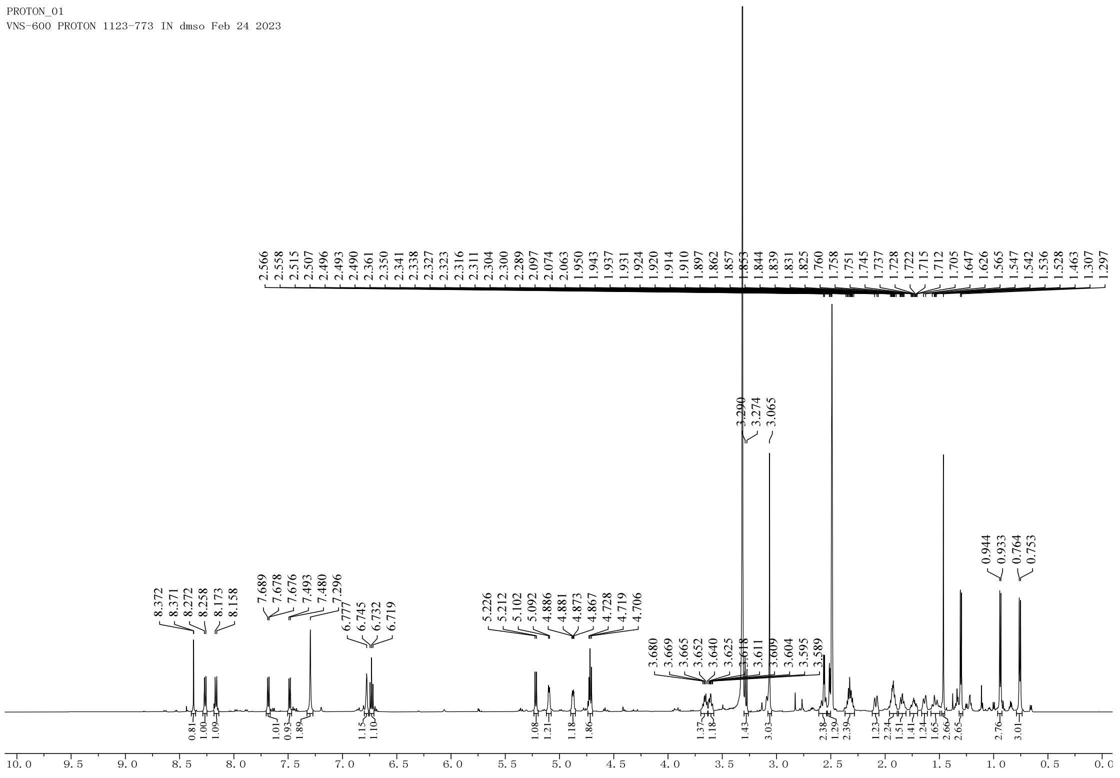

**Figure S16. $^{1}H$ NMR spectrum of compound 1 in DMSO-$d_6$ (600 MHz).**

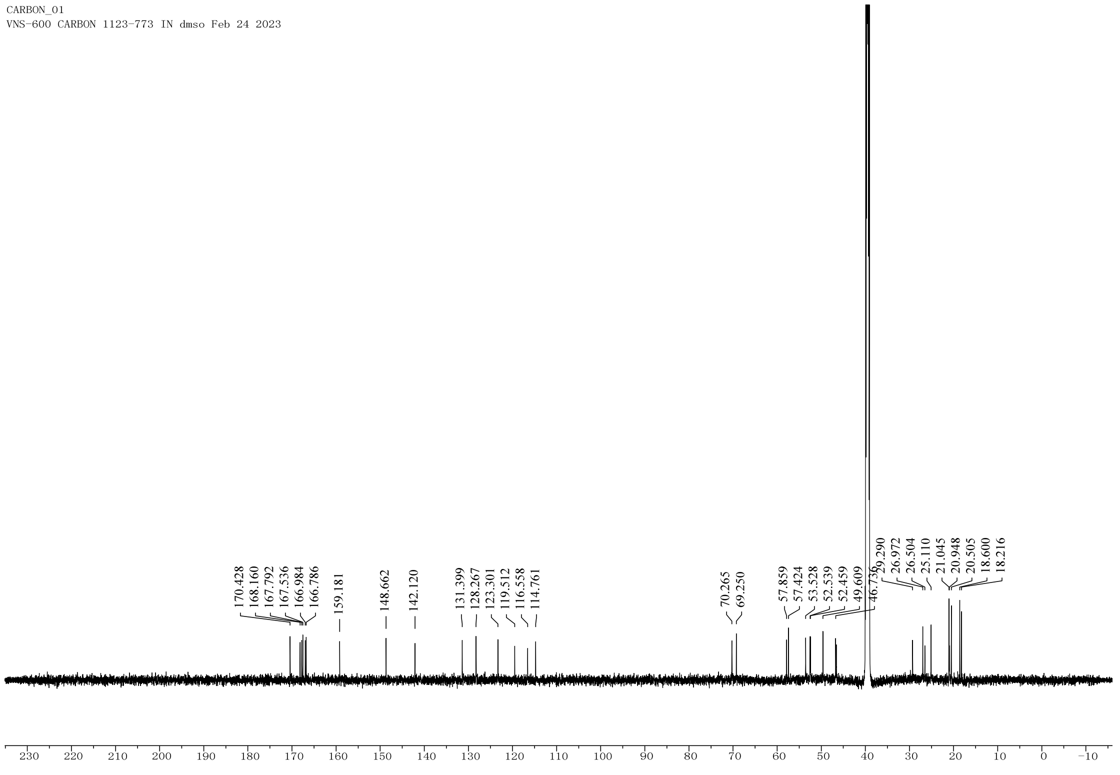

**Figure S17. $^{13}C$ NMR spectrum of compound 1 in DMSO-$d_6$ (150 MHz).**

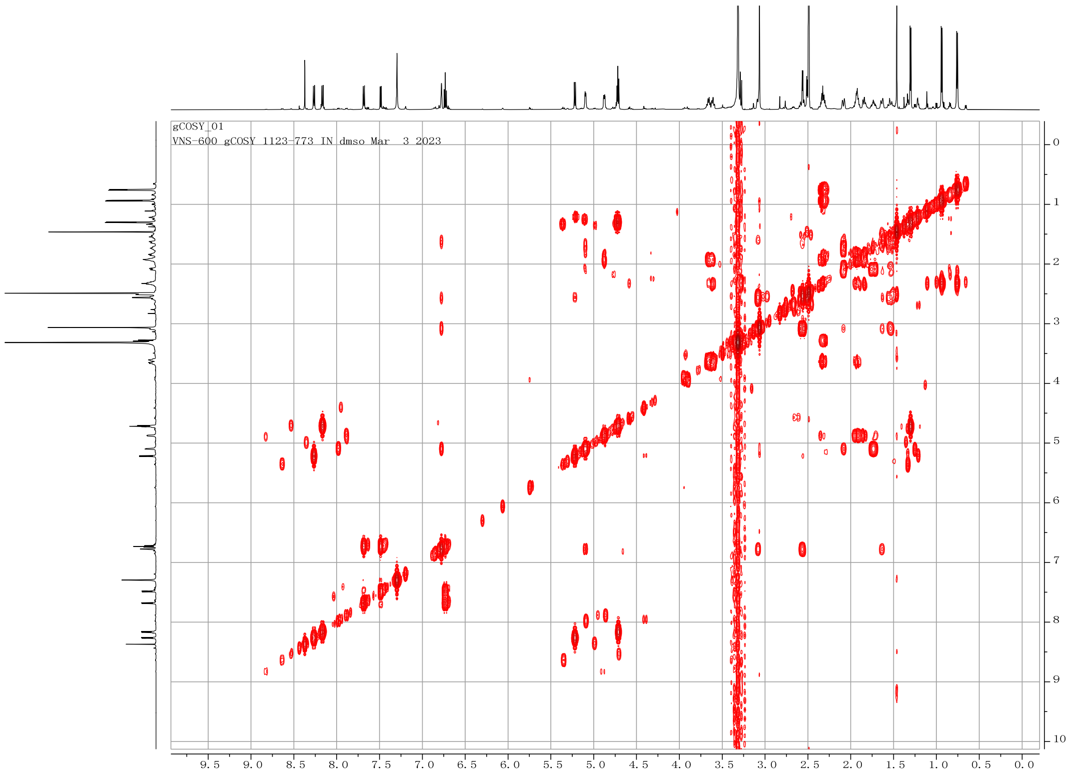

Figure S18. $^{1}H$-$^{1}H$ COSY spectrum of compound 1 in DMSO-$d_6$

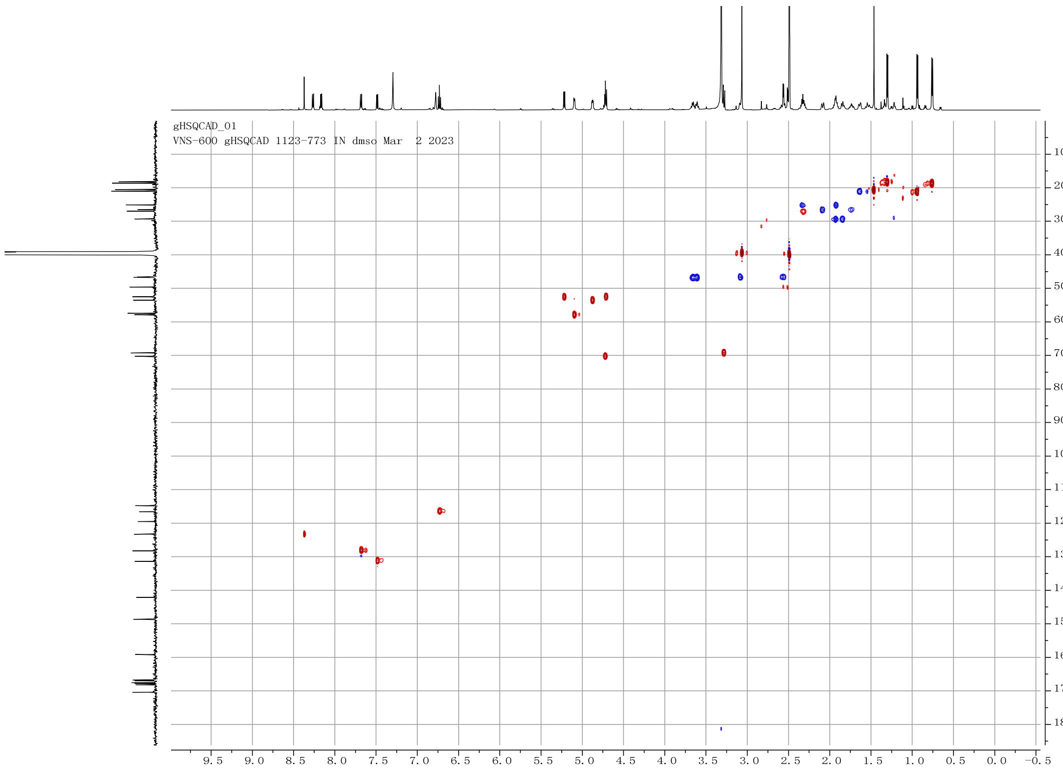

Figure S19. HSQC spectrum of compound 1 in DMSO-$d_6$

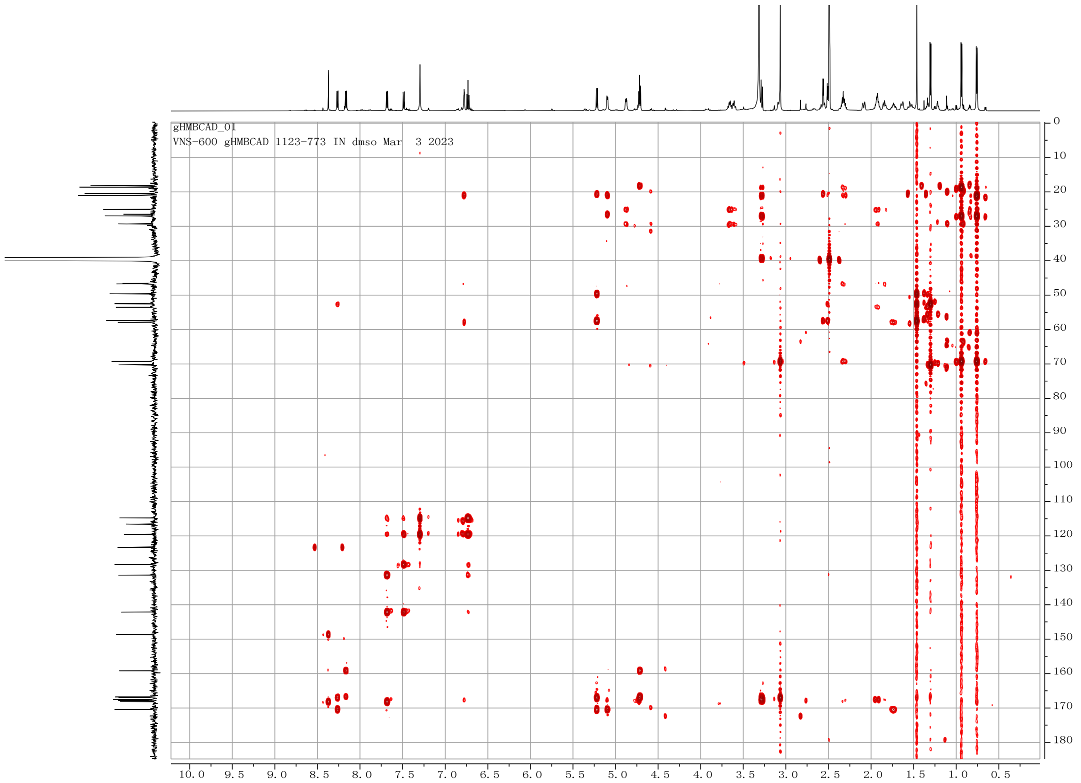

**Figure S20. HMBC spectrum of compound 1 in DMSO-$d_6$**

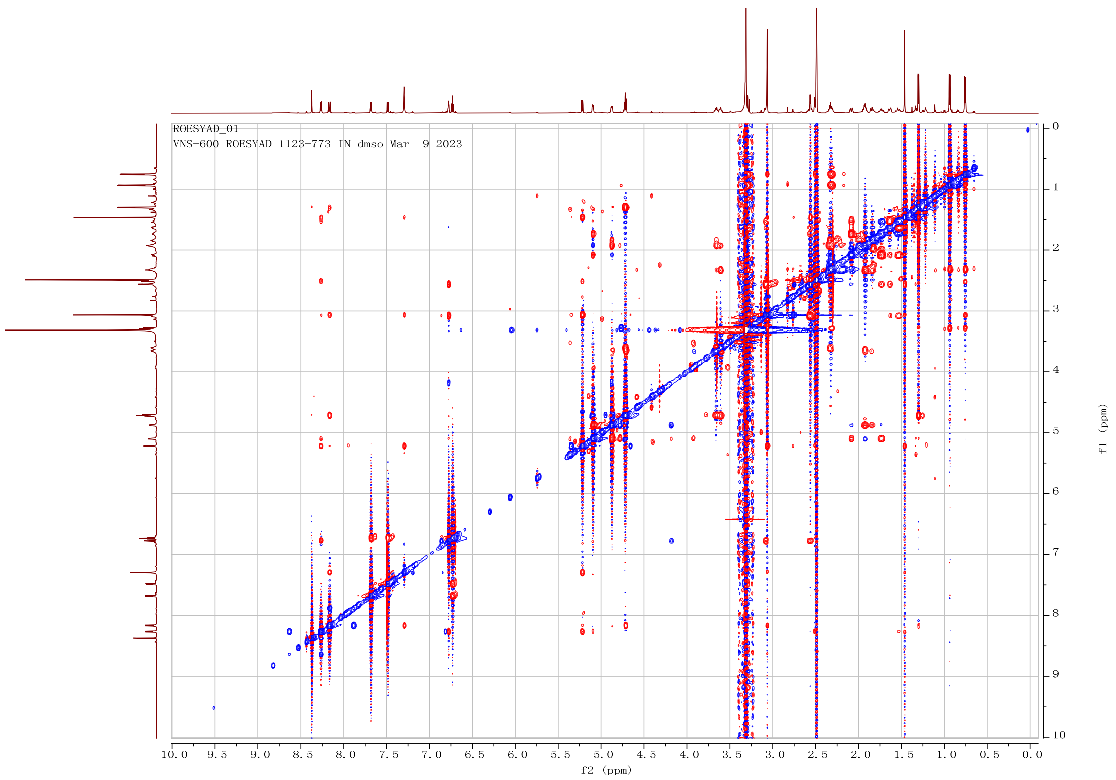

**Figure S21. ROESY spectrum of compound 1 in DMSO-$d_6$**

**Determination of the absolute configuration of aglomycin A (1)**

*Advanced Marfey's method.*

Algomycin A (1, 0.5 mg) was dissolved in 200 $\mu$L of 6 *N* HCl and hydrolyzed at 110 °C for 16 hours, after which the obtained hydrolysates were divided equally into two portions and dried under a flow of nitrogen ($N_2$). A 0.1 M $NaHCO_3$ solution (30 $\mu$L) was added to each of the dried hydrolysates of 1, as well as to the authentic standards of *N*-Me-*L*-Val, *L*-HPDA, *L*-Pro, *L*-Thr and *L*-*allo*-Thr, respectively. A solution of *L*-FDAA (1% in acetone, 30 $\mu$L) was then added to each reaction vial and kept at 40 °C for 1 hour, after which the reaction was quenched with 30 $\mu$L of 1 M HCl and subsequently diluted with 200 $\mu$L of MeCN. The derivatives of *D*-FDAA were prepared using the same procedure. The prepared derivatives were subjected to LC-MS analyses using an ACQUITY UPLC® BEH $C_{18}$ column (1.7 $\mu$m, 100 × 2.1 mm, at 30 °C) and eluted with an equant MeCN-$H_2O$ solution containing 1% formic acid (30:70, 0.3 mL/min, 13 min). The configuration of amino acid residues was determined by analyzing the retention times of the enantiomers *L*-DAA-*N*-Me-*L*-Val (6.57 min), *D*-DAA-*N*-Me-*L*-Val (6.82 min), *L*-DAA-*L*-HPDA (9.00 min), *D*-DAA-*L*-HPDA (7.90 min), *L*-DAA-*L*-Pro (8.55 min), *D*-DAA-*L*-Pro (9.22 min), *L*-DAA-*L*-Thr (6.26 min) and *D*-DAA-*L*-Thr (8.02 min) as shown Figure S22-S25. The amino acid residues of **1** were identified as *L*-*N*-Me-Val, *L*-HPDA, *L*-Pro, and *L*-Thr.

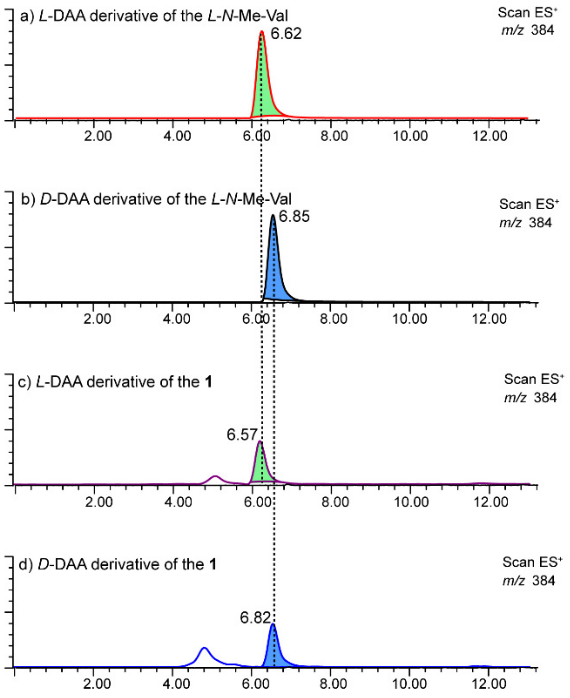

**Figure S22. The Marfey's analysis of the acid hydrolysates of compound 1, *L*-*N*-Me-Val and *D*-*N*-Me-Val.** a) Extracted ion chromatogram at *m/z* 384 for *L*-DAA-*L*-*N*-Me-Val. b) Extracted ion chromatogram at *m/z* 384 for *D*-DAA-*L*-*N*-Me-Val. c) Extracted ion chromatogram at *m/z* 384 for *L*-DAA derivatized of hydrolyzates of **1**. d) Extracted ion chromatogram at *m/z* 384 for *D*-DAA derivatized of hydrolyzates of **1**.

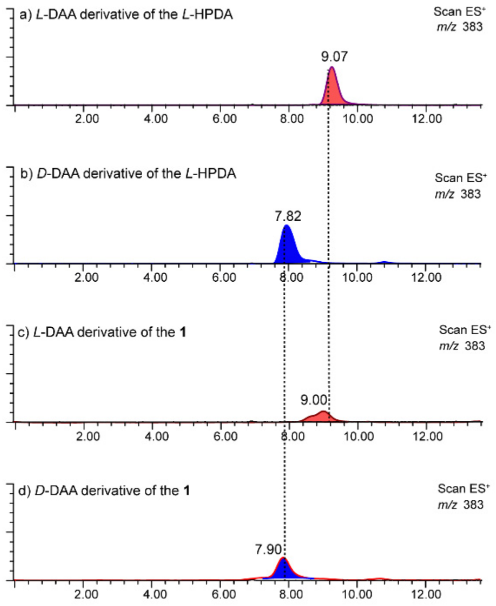

**Figure S23. The Marfey's analysis of the acid hydrolysates of compound 1, *L*-HPDA and *D*-HPDA.** a) Extracted ion chromatogram at *m/z* 383 for *L*-DAA-*L*-HPDA. b) Extracted ion chromatogram at *m/z* 383 for *D*-DAA-*L*-HPDA. c) Extracted ion chromatogram at *m/z* 383 for *L*-DAA derivatized of hydrolyzates of **1**. d) Extracted ion chromatogram at *m/z* 383 for *D*-DAA derivatized of hydrolyzates of **1**.

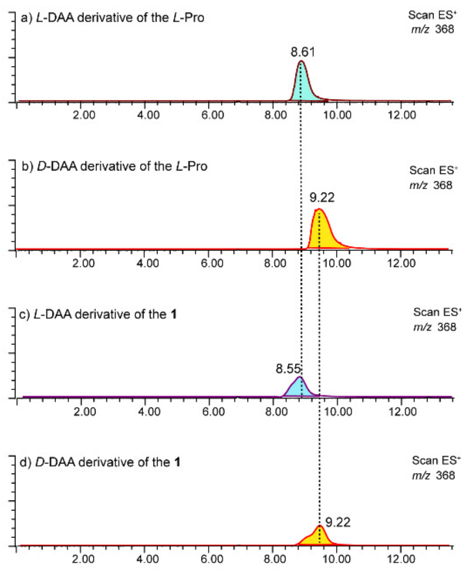

**Figure S24. The Marfey's analysis of the acid hydrolysates of compound 1, *L*-Pro and *D*-Pro.** a) Extracted ion chromatogram at *m/z* 368 for *L*-DAA-*L*-Pro. b) Extracted ion chromatogram at *m/z* 368 for *D*-DAA-*L*-Pro. c) Extracted ion chromatogram at *m/z* 368 for *L*-DAA derivatized of hydrolyzates of **1**. d) Extracted ion chromatogram at *m/z* 368 for *D*-DAA derivatized of hydrolyzates of **1**.

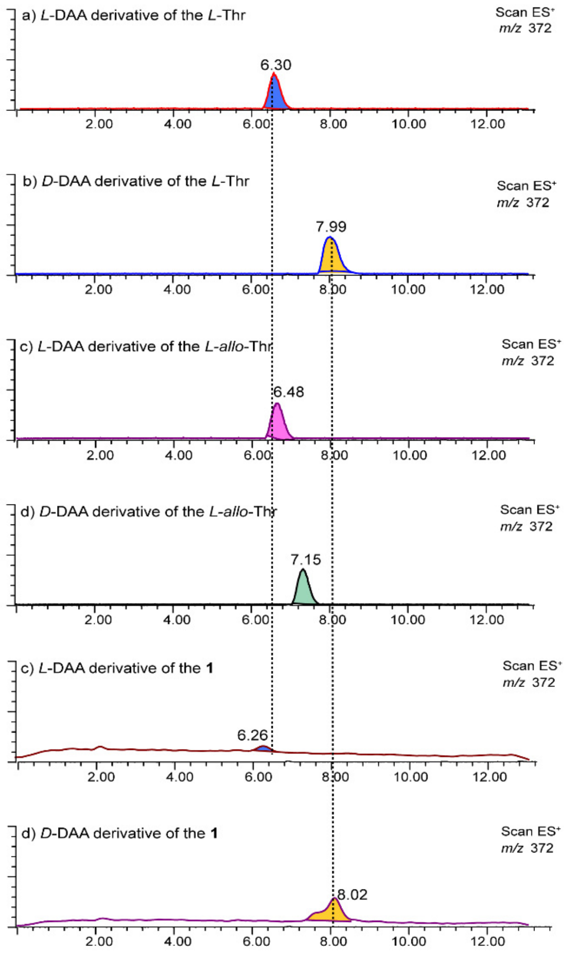

**Figure S25. The Marfey's analysis of the acid hydrolysates of compound 1, *L*-Thr and *L*-allo-Thr.** a) Extracted ion chromatogram at *m/z* 372 for *L*-DAA-*L*-Thr. b) Extracted ion chromatogram at *m/z* 372 for *D*-DAA-*L*-Thr. c) Extracted ion chromatogram at *m/z* 372 for *L*-DAA-*L*-*allo*-Thr. d) Extracted ion chromatogram at *m/z* 372 for *D*-DAA-*L*-*allo*-Thr. e) Extracted ion chromatogram at *m/z* 372 for *L*-DAA derivatized of hydrolyzates of **1**. f) Extracted ion chromatogram at *m/z* 372 for *D*-DAA derivatized of hydrolyzates of **1**.

*Theoretical NMR calculation.*

GIAO NMR calculations[39] and DP4+ analysis[40] were employed for the determination of the whole absolute configurations of **1**. To simplify the calculation, 2-(2-amino-3-chlorophenyl)-4-thiazolecarboxylic acid (ACTA) was replaced with acetic acid, which did not affect the accuracy of the result[41-42]. Conformational analysis of compound 1 was conducted using the Macromodel software with a genetic algorithm and the MMFF force field, applying an energy threshold of 2.5 kcal/mol. Geometry optimization was conducted using Gaussian 16 at the B3LYP/6-31+G(d) level, followed by NMR calculations at the PCM/mPW1PW91/6-311+G(d,p) level. For each potential diastereomer, the scaled chemical shift and the corrected mean absolute error (CMAE) were determined as previously described[43]. The DP4+ probability for each diastereomer was calculated using Boltzmann-averaged shielding tensor data through the universal and customizable DP4+ analysis method[44]. Notably, exchangeable protons data were excluded from the DP4+ analysis.

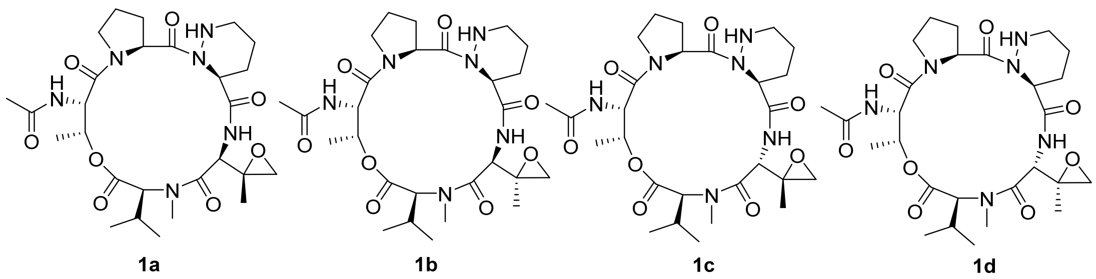

**Figure S26. The four possible diastereomers of 1 (1a-1d)**

Table S6. Experimental and calculated $^{13}$C NMR chemical shifts of 1a – 1d

| No. | $\delta_{exp}$ [a] | $\sigma^x$ (shielding constants)[b] | | | | $\delta_u$ (unscaled shifts)[c] | | | | $\delta_s$ (scaled shifts)[d] | | | |
|---|---|---|---|---|---|---|---|---|---|---|---|---|---|
| | | **1a** | **1b** | **1c** | **1d** | **1a** | **1b** | **1c** | **1d** | **1a** | **1b** | **1c** | **1d** |
| 1 | 167.8 | 12.40 | 11.75 | 12.90 | 12.89 | 180.81 | 181.46 | 180.31 | 180.32 | 166.35 | 166.14 | 167.07 | 166.90 |
| 2 | 69.3 | 115.15 | 112.60 | 124.30 | 123.95 | 78.06 | 80.61 | 68.91 | 69.27 | 68.57 | 70.65 | 61.05 | 61.22 |
| 3 | 27.0 | 158.20 | 158.52 | 157.56 | 157.53 | 35.01 | 34.70 | 35.65 | 35.68 | 27.60 | 27.19 | 29.39 | 29.25 |
| 4 | 21.1 | 166.77 | 166.42 | 168.15 | 168.10 | 26.44 | 25.98 | 23.37 | 23.43 | 19.44 | 18.94 | 17.70 | 17.59 |
| 5 | 18.6 | 166.95 | 167.23 | 169.84 | 169.79 | 26.26 | 26.79 | 25.06 | 25.11 | 19.27 | 19.71 | 19.31 | 19.20 |
| 6 | 39.4 | 146.00 | 144.43 | 154.00 | 154.26 | 47.21 | 48.78 | 39.22 | 38.95 | 39.20 | 40.52 | 32.78 | 32.37 |
| 7 | 167.0 | 13.69 | 11.14 | 13.14 | 12.26 | 179.53 | 182.07 | 180.07 | 180.95 | 165.13 | 166.71 | 166.84 | 167.50 |
| 8 | 52.5 | 129.64 | 128.66 | 127.54 | 126.72 | 63.58 | 64.55 | 65.68 | 66.49 | 54.78 | 55.46 | 57.97 | 58.58 |
| 9 | 57.4 | 128.30 | 128.13 | 130.83 | 127.61 | 64.91 | 65.08 | 62.38 | 65.60 | 56.05 | 55.95 | 54.83 | 57.73 |
| 10 | 49.6 | 132.84 | 132.60 | 133.00 | 134.56 | 60.37 | 60.61 | 60.21 | 58.65 | 51.73 | 51.73 | 52.77 | 51.11 |
| 11 | 20.5 | 169.39 | 169.62 | 170.24 | 169.08 | 23.83 | 23.59 | 22.97 | 24.13 | 16.95 | 16.67 | 17.32 | 18.27 |
| 12 | 170.4 | 11.23 | 11.45 | 13.48 | 13.69 | 181.98 | 181.76 | 179.74 | 179.52 | 167.47 | 166.42 | 166.52 | 166.14 |
| 13 | 57.9 | 125.55 | 125.70 | 128.47 | 128.78 | 67.66 | 67.52 | 64.74 | 64.43 | 58.67 | 58.26 | 57.08 | 56.61 |
| 14 | 26.5 | 158.86 | 159.61 | 161.17 | 160.82 | 34.35 | 33.60 | 32.05 | 32.39 | 26.96 | 26.15 | 25.96 | 26.13 |
| 15 | 21.0 | 164.33 | 163.93 | 163.88 | 163.93 | 28.88 | 29.28 | 29.33 | 29.28 | 21.76 | 22.06 | 23.38 | 23.17 |
| 16 | 46.6 | 139.53 | 139.72 | 137.85 | 137.85 | 53.68 | 53.49 | 55.36 | 55.36 | 45.36 | 44.99 | 48.15 | 47.99 |
| 18 | 167.5 | 11.79 | 10.28 | 16.55 | 16.78 | 181.42 | 182.93 | 176.66 | 176.43 | 166.94 | 167.53 | 163.60 | 163.20 |
| 20 | 53.5 | 130.36 | 129.21 | 128.55 | 128.78 | 62.85 | 64.01 | 64.66 | 64.43 | 54.09 | 54.94 | 57.00 | 56.61 |
| 21 | 29.3 | 155.16 | 155.31 | 155.42 | 155.50 | 38.06 | 37.90 | 37.79 | 37.71 | 30.49 | 30.22 | 31.43 | 31.19 |
| 23 | 25.1 | 159.46 | 160.41 | 159.06 | 159.24 | 33.76 | 32.80 | 34.15 | 33.97 | 26.40 | 25.40 | 27.96 | 27.63 |
| 28 | 46.7 | 138.06 | 137.95 | 137.65 | 137.79 | 55.15 | 55.26 | 55.56 | 55.42 | 46.76 | 46.66 | 48.34 | 48.04 |
| 30 | 166.8 | 14.46 | 14.19 | 12.51 | 12.52 | 178.75 | 179.02 | 180.70 | 180.69 | 164.40 | 163.82 | 167.44 | 167.25 |
| 31 | 52.5 | 130.48 | 130.33 | 131.68 | 131.28 | 62.73 | 62.88 | 61.53 | 61.93 | 53.97 | 53.87 | 54.02 | 54.23 |
| 32 | 70.3 | 115.90 | 116.21 | 119.63 | 119.77 | 77.31 | 77.00 | 73.58 | 73.44 | 67.85 | 67.24 | 65.49 | 65.19 |
| 33 | 18.2 | 169.73 | 169.77 | 169.44 | 169.33 | 23.48 | 23.44 | 23.77 | 23.88 | 16.62 | 16.54 | 18.08 | 18.03 |
| 34 | 159.2 | 9.70 | 9.93 | 9.57 | 9.12 | 183.51 | 183.28 | 183.65 | 184.09 | 168.93 | 167.86 | 170.25 | 170.49 |
| 35 | 22.9 | 12.40 | 11.75 | 12.90 | 12.89 | 28.88 | 28.80 | 28.73 | 28.71 | 21.76 | 21.61 | 22.81 | 22.62 |
| $R^2$ | | | | | | 0.9980 | 0.9980 | 0.9950 | 0.9950 | | | | |
| Slope (a) | | | | | | 1.0507 | 1.0563 | 1.0507 | 1.0508 | | | | |
| Intercept (b) | | | | | | 6.0190 | 5.9745 | 4.7681 | 4.9886 | | | | |
| CMAE | | | | | | | | | | 1.0251 | 1.0762 | 1.7739 | 1.6974 |

[a] the experimental chemical shifts abtained in DMSO-$d_6$. [b] the boltzmann averaged isotropic shielding constant were obtained by GIAO NMR computation at the mPW1PW91/6-311+G(d,p)//M062x/6-311+G(d,p) level of theory with PCM model in DMSO. [c] $\delta_u = \sigma^0 - \sigma^x$, where $\sigma^0$ (=193.21) is the shielding constant for the carbon nuclei in tetramethylsilane (TMS) calculated at the same level of theory used for $\sigma^x$. [d] $\delta_s = (\delta_u - b)/a$, where a and b were obtained from linear regression $\delta_u = a\delta_{exp} + b$. The corrected mean absolute error (CMAE) was defined as $\Sigma_n|\delta_s - \delta_{exp}|/n$

#### Table S7. Experimental and calculated $^1$H NMR chemical shifts of 1a – 1d

| No. | $\delta_{exp}$ [a] | $\sigma^x$ (shielding constants)[b] | | | | $\delta_u$ (unscaled shifts)[c] | | | | $\delta_s$ (scaled shifts)[d] | | | |
|---|---|---|---|---|---|---|---|---|---|---|---|---|---|
| | | **1a** | **1b** | **1c** | **1d** | **1a** | **1b** | **1c** | **1d** | **1a** | **1b** | **1c** | **1d** |
| 1 | 3.28 | 28.890 | 28.830 | 26.908 | 26.923 | 2.962 | 3.022 | 4.944 | 4.929 | 3.021 | 3.054 | 5.426 | 5.313 |
| 2 | 2.32 | 29.320 | 29.144 | 29.743 | 29.748 | 2.532 | 2.708 | 2.109 | 2.104 | 2.538 | 2.701 | 2.219 | 2.205 |
| 3 | 0.94 | 30.717 | 30.794 | 30.958 | 30.965 | 1.135 | 1.058 | 0.894 | 0.887 | 0.968 | 0.846 | 0.845 | 0.866 |
| 4 | 0.94 | 30.717 | 30.794 | 30.958 | 30.965 | 1.135 | 1.058 | 0.894 | 0.887 | 0.968 | 0.846 | 0.845 | 0.866 |
| 5 | 0.94 | 30.717 | 30.794 | 30.958 | 30.965 | 1.135 | 1.058 | 0.894 | 0.887 | 0.968 | 0.846 | 0.845 | 0.866 |
| 6 | 0.76 | 30.949 | 30.875 | 31.099 | 31.118 | 0.903 | 0.977 | 0.753 | 0.734 | 0.707 | 0.755 | 0.685 | 0.697 |
| 7 | 0.76 | 30.949 | 30.875 | 31.099 | 31.118 | 0.903 | 0.977 | 0.753 | 0.734 | 0.707 | 0.755 | 0.685 | 0.697 |
| 8 | 0.76 | 30.949 | 30.875 | 31.099 | 31.118 | 0.903 | 0.977 | 0.753 | 0.734 | 0.707 | 0.755 | 0.685 | 0.697 |
| 9 | 3.07 | 28.600 | 28.203 | 29.389 | 29.365 | 3.252 | 3.649 | 2.463 | 2.487 | 3.347 | 3.759 | 2.619 | 2.626 |
| 10 | 3.07 | 28.600 | 28.203 | 29.389 | 29.365 | 3.252 | 3.649 | 2.463 | 2.487 | 3.347 | 3.759 | 2.619 | 2.626 |
| 11 | 3.07 | 28.600 | 28.203 | 29.389 | 29.365 | 3.252 | 3.649 | 2.463 | 2.487 | 3.347 | 3.759 | 2.619 | 2.626 |
| 12 | 5.22 | 27.173 | 27.266 | 27.899 | 27.554 | 4.679 | 4.586 | 3.953 | 4.298 | 4.950 | 4.812 | 4.305 | 4.619 |
| 13 | 2.56 | 28.701 | 28.767 | 29.007 | 29.304 | 3.151 | 3.085 | 2.845 | 2.548 | 3.233 | 3.125 | 3.052 | 2.693 |
| 14 | 2.51 | 29.134 | 29.390 | 29.559 | 29.357 | 2.718 | 2.462 | 2.293 | 2.495 | 2.747 | 2.425 | 2.427 | 2.635 |
| 15 | 1.46 | 30.677 | 30.625 | 30.567 | 30.700 | 1.175 | 1.227 | 1.285 | 1.152 | 1.013 | 1.037 | 1.287 | 1.157 |
| 16 | 1.46 | 30.677 | 30.625 | 30.567 | 30.700 | 1.175 | 1.227 | 1.285 | 1.152 | 1.013 | 1.037 | 1.287 | 1.157 |
| 17 | 1.46 | 30.677 | 30.625 | 30.567 | 30.700 | 1.175 | 1.227 | 1.285 | 1.152 | 1.013 | 1.037 | 1.287 | 1.157 |
| 18 | 5.10 | 27.389 | 27.618 | 27.390 | 27.279 | 4.463 | 4.234 | 4.462 | 4.573 | 4.707 | 4.417 | 4.881 | 4.921 |
| 19 | 1.74 | 30.016 | 29.996 | 30.339 | 30.311 | 1.836 | 1.856 | 1.513 | 1.541 | 1.756 | 1.743 | 1.545 | 1.586 |
| 20 | 2.09 | 29.869 | 29.870 | 29.743 | 29.748 | 1.983 | 1.982 | 2.109 | 2.104 | 1.921 | 1.885 | 2.219 | 2.205 |
| 21 | 1.64 | 29.934 | 30.123 | 29.743 | 29.628 | 1.918 | 1.729 | 2.109 | 2.224 | 1.848 | 1.601 | 2.219 | 2.337 |
| 22 | 1.55 | 30.264 | 30.245 | 30.416 | 30.401 | 1.588 | 1.607 | 1.436 | 1.451 | 1.477 | 1.463 | 1.458 | 1.487 |
| 23 | 3.09 | 28.701 | 28.715 | 29.007 | 28.967 | 3.151 | 3.137 | 2.845 | 2.885 | 3.233 | 3.183 | 3.052 | 3.064 |
| 24 | 2.56 | 29.030 | 29.002 | 29.169 | 29.151 | 2.822 | 2.850 | 2.683 | 2.701 | 2.864 | 2.861 | 2.868 | 2.862 |
| 25 | 4.88 | 27.277 | 27.266 | 27.309 | 27.220 | 4.575 | 4.586 | 4.543 | 4.632 | 4.833 | 4.812 | 4.973 | 4.986 |
| 26 | 1.93 | 29.869 | 29.870 | 29.743 | 29.748 | 1.983 | 1.982 | 2.109 | 2.104 | 1.921 | 1.885 | 2.219 | 2.205 |
| 27 | 1.84 | 30.100 | 30.061 | 29.956 | 29.948 | 1.752 | 1.791 | 1.896 | 1.904 | 1.661 | 1.670 | 1.978 | 1.985 |
| 28 | 2.33 | 29.320 | 29.555 | 29.304 | 29.304 | 2.532 | 2.297 | 2.548 | 2.548 | 2.538 | 2.239 | 2.716 | 2.693 |
| 29 | 1.93 | 29.869 | 29.870 | 29.743 | 29.748 | 1.983 | 1.982 | 2.109 | 2.104 | 1.921 | 1.885 | 2.219 | 2.205 |
| 30 | 3.66 | 28.284 | 28.176 | 28.038 | 27.987 | 3.568 | 3.676 | 3.814 | 3.865 | 3.702 | 3.789 | 4.148 | 4.142 |
| 31 | 3.61 | 28.284 | 28.176 | 27.899 | 27.987 | 3.568 | 3.676 | 3.953 | 3.865 | 3.702 | 3.789 | 4.305 | 4.142 |
| 32 | 4.71 | 27.645 | 27.618 | 27.534 | 27.621 | 4.207 | 4.234 | 4.318 | 4.231 | 4.420 | 4.417 | 4.718 | 4.545 |
| 33 | 4.72 | 27.087 | 27.063 | 26.246 | 26.304 | 4.765 | 4.789 | 5.606 | 5.548 | 5.047 | 5.041 | 6.175 | 5.994 |
| 34 | 1.3 | 30.490 | 30.441 | 30.466 | 30.459 | 1.362 | 1.411 | 1.386 | 1.393 | 1.223 | 1.243 | 1.401 | 1.423 |
| 35 | 1.3 | 30.490 | 30.441 | 30.466 | 30.459 | 1.362 | 1.411 | 1.386 | 1.393 | 1.223 | 1.243 | 1.401 | 1.423 |
| 36 | 1.3 | 30.490 | 30.441 | 30.466 | 30.459 | 1.362 | 1.411 | 1.386 | 1.393 | 1.223 | 1.243 | 1.401 | 1.423 |
| 37 | 3.27 | 29.967 | 29.954 | 29.926 | 29.943 | 1.885 | 1.898 | 1.926 | 1.909 | 1.811 | 1.791 | 2.012 | 1.991 |
| 38 | 3.27 | 29.967 | 29.954 | 29.926 | 29.943 | 1.885 | 1.898 | 1.926 | 1.909 | 1.811 | 1.791 | 2.012 | 1.991 |
| 39 | 3.27 | 29.967 | 29.954 | 29.926 | 29.943 | 1.885 | 1.898 | 1.926 | 1.909 | 1.811 | 1.791 | 2.012 | 1.991 |
| $R^2$ | | | | | | 0.9579 | 0.9284 | 0.8090 | 0.8741 | | | | |
| Slope (a) | | | | | | 0.8900 | 0.8895 | 0.8839 | 0.9090 | | | | |
| Intercept | | | | | | 0.2734 | 0..3053 | 0.1477 | 0.0997 | | | | |
| CMAE | | | | | | | | | | 0.2543 | 0.2878 | 0.3545 | 0.3413 |

[a] the experimental chemical shifts abtained in DMSO-$d_6$. [b] the boltzmann averaged isotropic shielding constant were obtained by GIAO NMR computation at the mPW1PW91/6-311+G(d,p)//M062x/6-311+G(d,p) level of theory with PCM model in DMSO-$d_6$. [c] $\delta_u = \sigma^0 - \sigma^x$, where $\sigma^0$ (=31.852) is the shielding constant for the carbon nuclei in tetramethylsilane (TMS) calculated at the same level of theory used for $\sigma^x$. [d] $\delta_s = (\delta_u - b)/a$, where a and b were obtained from linear regression $\delta_u = a\delta_{exp} + b$. The corrected mean absolute error (CMAE) was defined as $\Sigma_n|\delta_s - \delta_{exp}|/n$.

Table S8. DP4+ Probabilities Computed for 1a – 1d

| | **1a** | **1b** | **1c** | **1d** |
|---|---|---|---|---|
| DP4+ (H data) | 97.34% | 2.66% | 0.00% | 0.09% |
| DP4+ (C data) | 99.68% | 0.32% | 0.00% | 0.00% |
| DP4+ (all data) | 99.99% | 0.01% | 0.00% | 0.00% |

| Functional | | Solvent? | | Basis Set | | Type of Data |
|---|---|---|---|---|---|---|
| mPW1PW91 | | PCM | | 6-311+G(d,p) | | Shielding Tensors |
| | | DP4+ | 99.99% | 0.01% | 0.00% | 0.00% | – |
| Nuclei | sp2? | Experimental | Isomer 1 | Isomer 2 | Isomer 3 | Isomer 4 | Isomer 5 |
| C | x | 167.8 | 12.404 | 11.747 | 12.897 | 12.892 | |
| C | | 69.3 | 115.146 | 112.604 | 124.296 | 123.945 | |
| C | | 27 | 158.196 | 158.515 | 157.561 | 157.534 | |
| C | | 21.1 | 166.769 | 166.416 | 168.154 | 168.100 | |
| C | | 18.6 | 166.946 | 167.228 | 169.84 | 169.785 | |
| C | | 39.4 | 146.002 | 144.432 | 153.995 | 154.260 | |
| C | x | 167 | 13.685 | 11.143 | 13.144 | 12.264 | |
| C | | 52.5 | 129.635 | 128.658 | 127.535 | 126.719 | |
| C | | 57.4 | 128.299 | 128.132 | 130.827 | 127.609 | |
| C | | 49.6 | 132.84 | 132.596 | 132.996 | 134.564 | |
| C | | 20.5 | 169.385 | 169.623 | 170.239 | 169.076 | |
| C | x | 170.4 | 11.231 | 11.447 | 13.475 | 13.687 | |
| C | | 57.9 | 125.549 | 125.695 | 128.469 | 128.781 | |
| C | | 26.5 | 158.86 | 159.609 | 161.165 | 160.816 | |
| C | | 21 | 164.326 | 163.929 | 163.88 | 163.927 | |
| C | | 46.6 | 139.531 | 139.717 | 137.846 | 137.846 | |
| C | x | 167.5 | 11.79 | 10.277 | 16.549 | 16.778 | |
| C | | 53.5 | 130.363 | 129.205 | 128.552 | 128.781 | |
| C | | 29.3 | 155.155 | 155.312 | 155.417 | 155.498 | |
| C | | 25.1 | 159.455 | 160.408 | 159.063 | 159.242 | |
| C | | 46.7 | 138.064 | 137.948 | 137.65 | 137.794 | |
| C | x | 166.8 | 14.459 | 14.191 | 12.511 | 12.524 | |
| C | | 52.5 | 130.484 | 130.329 | 131.681 | 131.282 | |
| C | | 70.3 | 115.903 | 116.208 | 119.627 | 119.767 | |
| C | | 18.2 | 169.726 | 169.766 | 169.443 | 169.327 | |
| C | x | 159.2 | 9.7 | 9.929 | 9.565 | 9.118 | |
| C | | 22.9 | 164.326 | 164.408 | 164.480 | 164.501 | |
| H | | 3.3 | 28.890 | 28.830 | 26.908 | 26.923 | |
| H | | 2.3 | 29.320 | 29.144 | 29.743 | 29.748 | |
| H | | 0.9 | 30.717 | 30.794 | 30.958 | 30.965 | |
| H | | 0.9 | 30.717 | 30.794 | 30.958 | 30.965 | |
| H | | 0.9 | 30.717 | 30.794 | 30.958 | 30.965 | |
| H | | 0.8 | 30.949 | 30.875 | 31.099 | 31.118 | |
| H | | 0.8 | 30.949 | 30.875 | 31.099 | 31.118 | |
| H | | 0.8 | 30.949 | 30.875 | 31.099 | 31.118 | |
| H | | 3.1 | 28.600 | 28.203 | 29.389 | 29.365 | |
| H | | 3.1 | 28.600 | 28.203 | 29.389 | 29.365 | |
| H | | 3.1 | 28.600 | 28.203 | 29.389 | 29.365 | |
| H | | 5.2 | 27.173 | 27.266 | 27.899 | 27.554 | |
| H | | 2.6 | 28.701 | 28.767 | 29.007 | 29.304 | |
| H | | 2.5 | 29.134 | 29.390 | 29.559 | 29.357 | |
| H | | 1.5 | 30.677 | 30.625 | 30.567 | 30.700 | |
| H | | 1.5 | 30.677 | 30.625 | 30.567 | 30.700 | |
| H | | 1.5 | 30.677 | 30.625 | 30.567 | 30.700 | |
| H | | 5.1 | 27.389 | 27.618 | 27.39 | 27.279 | |
| H | | 1.74 | 30.016 | 29.996 | 30.339 | 30.311 | |
| H | | 2.09 | 29.869 | 29.870 | 29.743 | 29.748 | |
| H | | 1.64 | 29.934 | 30.123 | 29.743 | 29.628 | |
| H | | 1.55 | 30.264 | 30.245 | 30.416 | 30.401 | |
| H | | 3.09 | 28.701 | 28.715 | 29.007 | 28.967 | |
| H | | 2.56 | 29.030 | 29.002 | 29.169 | 29.151 | |
| H | | 4.88 | 27.277 | 27.266 | 27.309 | 27.220 | |
| H | | 1.93 | 29.869 | 29.870 | 29.743 | 29.748 | |
| H | | 1.84 | 30.100 | 30.061 | 29.956 | 29.948 | |
| H | | 2.33 | 29.320 | 29.555 | 29.304 | 29.304 | |
| H | | 1.93 | 29.869 | 29.870 | 29.743 | 29.748 | |
| H | | 3.66 | 28.284 | 28.176 | 28.038 | 27.987 | |
| H | | 3.61 | 28.284 | 28.176 | 27.899 | 27.987 | |
| H | | 4.71 | 27.645 | 27.618 | 27.534 | 27.621 | |
| H | | 4.72 | 27.087 | 27.063 | 26.246 | 26.304 | |
| H | | 1.30 | 30.490 | 30.441 | 30.466 | 30.459 | |
| H | | 1.30 | 30.490 | 30.441 | 30.466 | 30.459 | |
| H | | 1.30 | 30.490 | 30.441 | 30.466 | 30.459 | |
| H | | 3.27 | 29.967 | 29.954 | 29.926 | 29.943 | |
| H | | 3.27 | 29.967 | 29.954 | 29.926 | 29.943 | |
| H | | 3.27 | 29.967 | 29.954 | 29.926 | 29.943 | |

Table S9. Lowest-energy conformers optimized at the M062X/6-311+G(d, p) Level of 1a in DMSO with Relative Energies < 2.5 kcal/mol

| **1a_C1** | ΔG 0.00 kcal/mol<br>Boltzman pop. 97.88% |
|---|---|
| 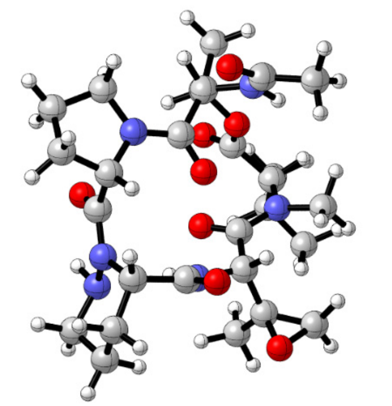 | |

Table S10. Atomic coordinates for the lowest-energy conformers of 1a (1a_C1)

| Atoms | 1a_C1 | | |
|---|---|---|---|
| | x | y | z |
| C | -0.4070 | -3.1692 | 0.2266 |
| C | -1.3504 | -2.8593 | -0.9422 |
| O | -2.1271 | -1.7319 | -0.5001 |
| C | -2.6850 | -0.9314 | -1.4175 |
| O | 0.5427 | -1.0717 | 0.9547 |
| C | 0.7023 | -2.1108 | 0.3245 |
| C | -3.4363 | 0.2002 | -0.7089 |
| N | -2.5880 | 0.7499 | 0.3544 |
| O | 1.8382 | 1.6004 | 2.5456 |
| O | 2.3321 | -0.4792 | -2.3274 |
| C | -1.3647 | 1.1932 | -0.0079 |
| C | -0.5217 | 1.9862 | 1.0024 |
| N | 0.8417 | 1.9740 | 0.5331 |
| C | 1.8837 | 1.6682 | 1.3248 |
| C | 2.7420 | -0.2789 | -1.1885 |
| C | 2.9650 | -1.4559 | -0.2379 |
| N | 1.8430 | -2.3839 | -0.3383 |
| C | 3.2215 | 1.4100 | 0.6115 |
| N | 3.0731 | 0.9750 | -0.7790 |
| O | -2.6313 | -1.1298 | -2.6002 |
| C | -3.1182 | 0.7180 | 1.7122 |
| C | -4.0073 | 1.2631 | -1.6614 |
| C | -4.5630 | 2.4438 | -0.8610 |
| C | -5.1220 | 0.6669 | -2.5256 |
| O | -0.9544 | 1.0635 | -1.1567 |
| C | 4.1639 | -2.3043 | -0.6981 |
| C | 3.5441 | -3.2984 | -1.6862 |
| C | 2.1825 | -3.6190 | -1.0662 |
| C | 4.1547 | 2.6297 | 0.6904 |
| C | 3.7764 | 3.7092 | -0.3197 |
| C | 3.6935 | 3.0828 | -1.7047 |
| N | 2.6819 | 2.0201 | -1.6709 |
| C | -1.0525 | 3.4200 | 1.0666 |
| C | -1.0789 | 4.1763 | -0.2381 |
| C | -1.9228 | 3.8246 | 2.1815 |
| O | -0.5359 | 4.1617 | 2.1644 |
| C | -2.2600 | -4.0247 | -1.2868 |
| N | -1.1528 | -3.2369 | 1.4618 |
| C | -0.5375 | -3.6761 | 2.5890 |
| C | -1.3125 | -3.5266 | 3.8734 |
| O | 0.5847 | -4.1653 | 2.5532 |
| H | 0.0412 | -4.1476 | 0.0511 |
| H | -0.7649 | -2.5532 | -1.8131 |
| H | -4.2787 | -0.2836 | -0.2021 |
| H | -0.5294 | 1.5490 | 1.9997 |
| H | 0.9863 | 1.9728 | -0.4735 |
| H | 3.0561 | -1.1368 | 0.7959 |
| H | 3.6939 | 0.6094 | 1.1729 |
| H | -2.3242 | 0.8377 | 2.4429 |
| H | -3.8700 | 1.4976 | 1.8608 |
| H | -3.5815 | -0.2545 | 1.8855 |
| H | -3.1949 | 1.6105 | -2.3054 |
| H | -4.9593 | 3.1998 | -1.5418 |
| H | -3.8062 | 2.9191 | -0.2335 |
| H | -5.3828 | 2.1146 | -0.2138 |

| | | | |
|---|---|---|---|
| H | -5.5266 | 1.4350 | -3.1881 |
| H | -4.7688 | -0.1614 | -3.1374 |
| H | -5.9405 | 0.3101 | -1.8919 |
| H | 4.5705 | -2.8289 | 0.1686 |
| H | 4.9555 | -1.6968 | -1.1366 |
| H | 3.4042 | -2.8225 | -2.6564 |
| H | 4.1496 | -4.1944 | -1.8153 |
| H | 2.2518 | -4.4561 | -0.3648 |
| H | 1.4309 | -3.8501 | -1.8224 |
| H | 5.1708 | 2.2818 | 0.4838 |
| H | 4.1338 | 3.0068 | 1.7141 |
| H | 2.8069 | 4.1497 | -0.0667 |
| H | 4.5204 | 4.5074 | -0.3100 |
| H | 4.6747 | 2.6908 | -2.0034 |
| H | 3.3619 | 3.8051 | -2.4511 |
| H | 2.5846 | 1.5696 | -2.5776 |
| H | -1.4283 | 5.1944 | -0.0656 |
| H | -0.0732 | 4.2177 | -0.6649 |
| H | -1.7368 | 3.6942 | -0.9639 |
| H | -2.2021 | 3.1101 | 2.9477 |
| H | -2.6023 | 4.6583 | 2.0358 |
| H | -2.9509 | -3.7379 | -2.0793 |
| H | -1.6594 | -4.8665 | -1.6362 |
| H | -2.8281 | -4.3368 | -0.4088 |
| H | -1.9628 | -2.6393 | 1.5494 |
| H | -2.3573 | -3.2680 | 3.7071 |
| H | -1.2488 | -4.4591 | 4.4331 |
| H | -0.8399 | -2.7422 | 4.4678 |

#### Table S11. Lowest-energy conformers optimized at the M062X/6-311+G(d, p) level of 1b in DMSO with relative energies < 2.5 kcal/mol

| **1b_C1** ΔG 0.00 kcal/mol Boltzman pop. 95.69% | **1b_C2** ΔG 2.19 kcal/mol Boltzman pop. 2.37% | **1b_C3** ΔG 2.32 kcal/mol Boltzman pop. 1.92% |
|---|---|---|
| 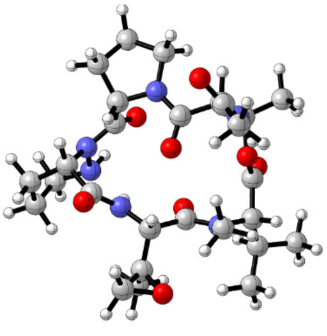 | 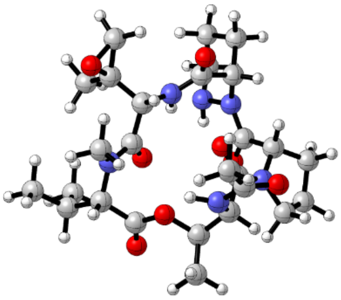 | 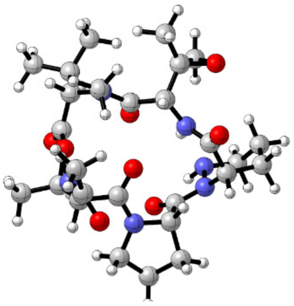 |

#### Table S12. Atomic coordinates for the lowest-energy conformers of 1b (1b_C1–1b_C3)

| Atoms | 1b_C1 | | | 1b_C2 | | | 1b_C3 | | |
|---|---|---|---|---|---|---|---|---|---|
| | x | y | z | x | y | z | x | y | z |
| C | 1.9617 | 2.5820 | 0.0554 | 0.8585 | -3.0098 | 0.3804 | -2.6225 | -1.9737 | 0.0615 |
| C | 2.5465 | 1.8183 | -1.1389 | -0.1170 | -3.1614 | -0.7925 | -3.0168 | -1.0463 | -1.0933 |
| O | 2.7267 | 0.4779 | -0.6504 | -1.2483 | -2.3370 | -0.4513 | -2.7890 | 0.2807 | -0.5828 |
| C | 2.7431 | -0.5442 | -1.5163 | -2.0905 | -1.9819 | -1.4323 | -2.5633 | 1.2809 | -1.4436 |
| O | 0.2819 | 1.0946 | 0.9321 | 1.0771 | -0.6875 | 1.0095 | -0.5780 | -1.0352 | 0.9241 |
| C | 0.5179 | 2.1212 | 0.3034 | 1.5348 | -1.6304 | 0.3758 | -1.1008 | -1.9303 | 0.2684 |
| C | 2.8573 | -1.8489 | -0.7217 | -3.2902 | -1.2365 | -0.8540 | -2.2705 | 2.5656 | -0.6603 |
| N | 1.8453 | -1.8550 | 0.3422 | -2.8828 | -0.2516 | 0.1549 | -1.3140 | 2.2767 | 0.4164 |
| O | -2.5839 | -0.5586 | 2.4828 | 0.9089 | 2.1790 | 2.5427 | 2.5249 | -0.2549 | 2.5293 |
| O | -1.6720 | 1.4633 | -2.2288 | 2.4264 | 0.4925 | -2.4010 | 1.1792 | -1.7489 | -2.2626 |
| C | 0.5541 | -1.7540 | -0.0440 | -1.8242 | 0.5450 | -0.1221 | -0.0745 | 1.8904 | 0.0336 |
| C | -0.5330 | -1.8519 | 1.0329 | -1.3384 | 1.5380 | 0.9528 | 0.9962 | 1.6598 | 1.1101 |
| N | -1.7224 | -1.2383 | 0.4941 | -0.0661 | 2.0430 | 0.4967 | 1.9270 | 0.7136 | 0.5608 |
| C | -2.5967 | -0.5658 | 1.2578 | 0.9830 | 2.1906 | 1.3202 | 2.5644 | -0.1993 | 1.3065 |
| C | -2.2264 | 1.4691 | -1.1377 | 2.6892 | 0.7694 | -1.2335 | 1.7044 | -1.9628 | -1.1780 |
| C | -1.8456 | 2.5010 | -0.0701 | 3.3962 | -0.2616 | -0.3547 | 1.0609 | -2.9254 | -0.1742 |
| N | -0.4513 | 2.8795 | -0.2431 | 2.6489 | -1.5193 | -0.3740 | -0.3859 | -2.8979 | -0.3354 |
| C | -3.6504 | 0.2645 | 0.5056 | 2.3565 | 2.4090 | 0.6649 | 3.3717 | -1.2484 | 0.5239 |
| N | -3.2415 | 0.6013 | -0.8600 | 2.4235 | 2.0064 | -0.7401 | 2.9100 | -1.4080 | -0.8591 |
| O | 2.7037 | -0.4139 | -2.7088 | -1.9595 | -2.3121 | -2.5798 | -2.6136 | 1.1625 | -2.6374 |
| C | 2.2644 | -1.5744 | 1.7201 | -3.4469 | -0.4438 | 1.4908 | -1.8323 | 1.9580 | 1.7534 |
| C | 2.8704 | -3.1099 | -1.5982 | -4.2719 | -0.7265 | -1.9212 | -1.9038 | 3.7366 | -1.5902 |
| C | 2.7676 | -4.3604 | -0.7237 | -3.7159 | 0.3386 | -2.8652 | -1.1733 | 4.8455 | -0.8357 |
| C | 4.1628 | -3.1569 | -2.4185 | -5.5493 | -0.2261 | -1.2443 | -3.1862 | 4.2787 | -2.2280 |
| O | 0.2273 | -1.6698 | -1.2237 | -1.2708 | 0.5200 | -1.2146 | 0.2446 | 1.7969 | -1.1458 |
| C | -2.5940 | 3.8248 | -0.2995 | 4.7569 | -0.6490 | -0.9627 | 1.4190 | -4.3831 | -0.5112 |
| C | -1.7272 | 4.5277 | -1.3504 | 4.4349 | -1.8423 | -1.8683 | 0.3752 | -4.7570 | -1.5694 |
| C | -0.2948 | 4.1684 | -0.9413 | 3.3522 | -2.5941 | -1.0944 | -0.8989 | -4.0592 | -1.0838 |
| C | -5.0366 | -0.3967 | 0.5118 | 2.8177 | 3.8681 | 0.8118 | 4.8799 | -0.9619 | 0.5646 |
| C | -5.1524 | -1.5062 | -0.5287 | 2.1021 | 4.7852 | -0.1771 | 5.2806 | 0.1388 | -0.4124 |
| C | -4.7351 | -0.9539 | -1.8842 | 2.2627 | 4.2251 | -1.5840 | 4.7645 | -0.2267 | -1.7970 |
| N | -3.3484 | -0.4885 | -1.7824 | 1.6982 | 2.8694 | -1.6144 | 3.3042 | -0.3397 | -1.7289 |
| C | -0.7978 | -3.3219 | 1.3694 | -2.3105 | 2.7048 | 1.1611 | 1.6759 | 3.0093 | 1.4196 |
| C | -1.2230 | -4.2097 | 0.2289 | -2.9047 | 3.3120 | -0.0809 | 0.9179 | 3.9649 | 2.3033 |
| C | -1.1888 | -3.6380 | 2.7478 | -2.1193 | 3.5078 | 2.3743 | 2.6767 | 3.5106 | 0.4690 |
| O | 0.1400 | -3.8923 | 2.2773 | -3.2008 | 2.5757 | 2.2655 | 3.0623 | 2.9121 | 1.7076 |
| C | 3.8675 | 2.3893 | -1.6211 | -0.5511 | -4.6032 | -0.9937 | -4.4640 | -1.2091 | -1.5209 |
| N | 2.7747 | 2.3592 | 1.2294 | 0.1695 | -3.2309 | 1.6292 | -3.3076 | -1.5750 | 1.2702 |
| C | 2.5000 | 3.0413 | 2.3691 | 0.8877 | -3.3558 | 2.7749 | -3.1780 | -2.3304 | 2.3901 |
| C | 3.2557 | 2.6153 | 3.6018 | 0.0976 | -3.3643 | 4.0588 | -3.7362 | -1.7406 | 3.6600 |
| O | 1.6796 | 3.9509 | 2.3830 | 2.1078 | -3.4564 | 2.7551 | -2.6299 | -3.4251 | 2.3575 |
| H | 1.9796 | 3.6474 | -0.1746 | 1.6259 | -3.7789 | 0.2809 | -2.9350 | -2.9875 | -0.1907 |
| H | 1.8138 | 1.7968 | -1.9502 | 0.3352 | -2.7601 | -1.7028 | -2.3427 | -1.2117 | -1.9378 |
| H | 3.8245 | -1.7927 | -0.2130 | -3.8207 | -2.0145 | -0.2923 | -3.2063 | 2.8264 | -0.1575 |
| H | -0.2476 | -1.3149 | 1.9372 | -1.1820 | 1.0307 | 1.9074 | 0.5789 | 1.2392 | 2.0255 |
| H | -1.8249 | -1.2243 | -0.5182 | 0.0993 | 2.0415 | -0.5048 | 2.0517 | 0.7091 | -0.4489 |
| H | -1.9737 | 2.1056 | 0.9344 | 3.4957 | 0.0702 | 0.6740 | 1.2979 | -2.6550 | 0.8516 |
| H | -3.7399 | 1.1924 | 1.0669 | 3.0343 | 1.7847 | 1.2384 | 3.2120 | -2.1912 | 1.0424 |
| H | 1.7349 | -2.2196 | 2.4175 | -3.2172 | 0.3951 | 2.1385 | -2.8696 | 2.2811 | 1.8048 |
| H | 3.3276 | -1.7881 | 1.8055 | -4.5319 | -0.5165 | 1.4102 | -1.7783 | 0.8813 | 1.9306 |
| H | 2.0821 | -0.5230 | 1.9608 | -3.0635 | -1.3691 | 1.9311 | -1.2811 | 2.4889 | 2.5277 |
| H | 2.0152 | -3.0664 | -2.2756 | -4.5211 | -1.6110 | -2.5169 | -1.2527 | 3.3550 | -2.3772 |
| H | 2.8249 | -5.2559 | -1.3460 | -4.4657 | 0.5635 | -3.6279 | -1.0619 | 5.7184 | -1.4826 |
| H | 1.8361 | -4.3951 | -0.1571 | -2.8023 | 0.0067 | -3.3556 | -0.1756 | 4.5333 | -0.5199 |
| H | 3.5961 | -4.3942 | -0.0080 | -3.4952 | 1.2628 | -2.3281 | -1.7360 | 5.1522 | 0.0520 |
| H | 4.1655 | -4.0418 | -3.0584 | -6.2811 | 0.0730 | -1.9970 | -2.9445 | 5.0403 | -2.9720 |
| H | 4.2768 | -2.2784 | -3.0532 | -6.0049 | -0.9962 | -0.6160 | -3.7541 | 3.4898 | -2.7245 |

| | | | | | | | | | |
|---|---|---|---|---|---|---|---|---|---|
| H | 5.0311 | -3.2229 | -1.7549 | -5.3385 | 0.6472 | -0.6191 | -3.8253 | 4.7399 | -1.4683 |
| H | -2.5996 | 4.3881 | 0.6356 | 5.4231 | -0.9539 | -0.1534 | 1.2914 | -4.9901 | 0.3871 |
| H | -3.6244 | 3.6712 | -0.6202 | 5.2197 | 0.1813 | -1.4953 | 2.4463 | -4.4890 | -0.8597 |
| H | -1.9400 | 4.1250 | -2.3411 | 4.0322 | -1.4918 | -2.8180 | 0.6673 | -4.3573 | -2.5409 |
| H | -1.8829 | 5.6052 | -1.3711 | 5.3039 | -2.4706 | -2.0596 | 0.2366 | -5.8332 | -1.6615 |
| H | 0.1225 | 4.9099 | -0.2545 | 3.7903 | -3.2975 | -0.3790 | -1.4764 | -4.7034 | -0.4147 |
| H | 0.3635 | 4.0701 | -1.8060 | 2.6749 | -3.1367 | -1.7548 | -1.5341 | -3.7437 | -1.9129 |
| H | -5.7743 | 0.3801 | 0.2925 | 3.8946 | 3.8979 | 0.6238 | 5.4015 | -1.8864 | 0.3013 |
| H | -5.2309 | -0.7629 | 1.5212 | 2.6490 | 4.1768 | 1.8444 | 5.1463 | -0.7108 | 1.5926 |
| H | -4.5023 | -2.3478 | -0.2694 | 1.0359 | 4.8481 | 0.0636 | 4.8488 | 1.0978 | -0.1074 |
| H | -6.1786 | -1.8750 | -0.5688 | 2.5176 | 5.7928 | -0.1238 | 6.3662 | 0.2480 | -0.4293 |
| H | -5.4100 | -0.1427 | -2.1878 | 3.3228 | 4.2205 | -1.8701 | 5.2249 | -1.1634 | -2.1378 |
| H | -4.7477 | -1.7272 | -2.6527 | 1.7049 | 4.8122 | -2.3141 | 4.9805 | 0.5557 | -2.5250 |
| H | -3.0151 | -0.1179 | -2.6687 | 1.7942 | 2.4498 | -2.5362 | 2.9071 | -0.5764 | -2.6345 |
| H | -1.4005 | -5.2210 | 0.5954 | -3.4517 | 4.2218 | 0.1664 | -0.0380 | 4.2430 | 1.8553 |
| H | -2.1428 | -3.8311 | -0.2216 | -2.1174 | 3.5560 | -0.7988 | 0.7318 | 3.5087 | 3.2791 |
| H | -0.4546 | -4.2437 | -0.5469 | -3.5950 | 2.6066 | -0.5502 | 1.5073 | 4.8704 | 2.4501 |
| H | -1.7812 | -4.5275 | 2.9345 | -2.4171 | 4.5510 | 2.3626 | 2.8483 | 4.5799 | 0.4006 |
| H | -1.2737 | -2.8329 | 3.4695 | -1.3405 | 3.2245 | 3.0737 | 2.9186 | 2.9240 | -0.4124 |
| H | 4.5935 | 2.4154 | -0.8068 | -0.9871 | -4.9993 | -0.0751 | -5.1350 | -1.0448 | -0.6763 |
| H | 4.2616 | 1.7793 | -2.4339 | -1.2822 | -4.6695 | -1.7983 | -4.7006 | -0.4970 | -2.3112 |
| H | 3.7145 | 3.4051 | -1.9894 | 0.3158 | -5.2101 | -1.2613 | -4.6206 | -2.2191 | -1.9036 |
| H | 3.2843 | 1.4885 | 1.2887 | -0.7907 | -2.9219 | 1.6882 | -3.5643 | -0.6021 | 1.3648 |
| H | 4.0663 | 1.9226 | 3.3804 | -0.9712 | -3.4937 | 3.8947 | -2.8985 | -1.4187 | 4.2823 |
| H | 3.6539 | 3.5019 | 4.0943 | 0.4718 | -4.1639 | 4.6970 | -4.3894 | -0.8888 | 3.4757 |
| H | 2.5518 | 2.1363 | 4.2852 | 0.2681 | -2.4148 | 4.5701 | -4.2811 | -2.5142 | 4.1996 |

Table S13. Lowest-energy conformers optimized at the M062X/6-311+G(d, p) level of 1c in DMSO with relative energies < 2.5 kcal/mol

| **1c_C1** ΔG 0.00 kcal/mol Boltzman pop. 92.70% | **1c_C2** ΔG 1.82 kcal/mol Boltzman pop. 4.30% | **1c_C3** ΔG 2.12 kcal/mol Boltzman pop. 2.57% |
|---|---|---|
| 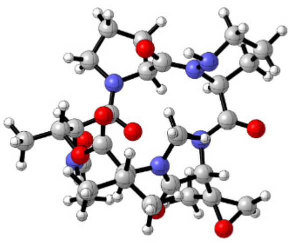 | 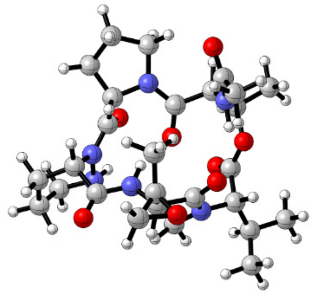 | 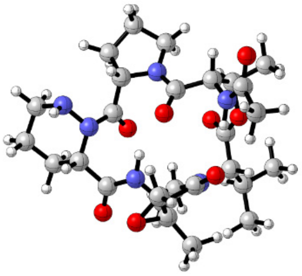 |

Table S14. Atomic coordinates for the lowest-energy conformers of 1c (1c_C1–1c_C3)

| Atoms | 1c_C1 | | | 1c_C2 | | | 1c_C3 | | |
|---|---|---|---|---|---|---|---|---|---|
| | x | y | z | x | y | z | x | y | z |
| C | 2.6199 | 1.9187 | 0.3222 | 1.7906 | 2.3436 | -0.5452 | 1.7205 | 2.3084 | 0.7997 |
| C | 3.2340 | 1.3595 | -0.9680 | 2.6697 | 1.6470 | -1.6003 | 2.5700 | 2.2233 | -0.4835 |
| O | 3.0362 | -0.0701 | -0.9384 | 2.9625 | 0.3302 | -1.0957 | 2.8235 | 0.8214 | -0.6844 |
| C | 2.1351 | -0.6131 | -1.7699 | 2.4248 | -0.7498 | -1.6734 | 2.3133 | 0.1863 | -1.7411 |
| O | 0.6168 | 0.6225 | 0.5207 | 0.1988 | 0.5889 | -0.5502 | -0.2162 | 0.9466 | 1.2273 |
| C | 1.0969 | 1.7418 | 0.3158 | 0.3596 | 1.8144 | -0.5913 | 0.2524 | 1.8990 | 0.6030 |
| C | 1.8700 | -2.0794 | -1.4424 | 2.5967 | -1.9759 | -0.7777 | 2.5235 | -1.3162 | -1.5881 |
| N | 0.5169 | -2.1737 | -0.8854 | 1.3051 | -2.2270 | -0.1209 | 1.2441 | -1.8774 | -1.1294 |
| O | -3.7637 | -1.7541 | 0.8139 | -3.1049 | -2.2529 | 1.7317 | -2.7126 | -3.0665 | -0.5848 |
| O | -1.2380 | 2.1929 | -2.1299 | -1.9325 | 1.1980 | -2.6647 | -1.7216 | 0.3537 | -1.4176 |
| C | 0.3942 | -2.4136 | 0.4494 | 1.0851 | -1.6063 | 1.0688 | 1.0330 | -1.9494 | 0.2099 |
| C | -1.0240 | -2.3323 | 1.0518 | -0.3076 | -1.7905 | 1.7138 | -0.2730 | -2.6122 | 0.6879 |
| N | -1.6145 | -1.0338 | 0.7905 | -1.3112 | -1.1548 | 0.8821 | -1.3726 | -1.6753 | 0.6028 |
| C | -2.9457 | -0.8588 | 0.6719 | -2.6273 | -1.4041 | 0.9936 | -2.5227 | -1.9919 | -0.0319 |
| C | -1.6854 | 1.9489 | -1.0125 | -2.3035 | 1.0432 | -1.5067 | -2.3021 | 0.9745 | -0.5477 |
| C | -1.1246 | 2.6741 | 0.2124 | -2.0399 | 2.1304 | -0.4638 | -1.9633 | 2.4221 | -0.2053 |
| N | 0.3263 | 2.8084 | 0.0878 | -0.6732 | 2.6588 | -0.6041 | -0.5086 | 2.6334 | -0.2246 |
| C | -3.3529 | 0.6016 | 0.3754 | -3.5151 | -0.4350 | 0.1807 | -3.6691 | -0.9893 | 0.0753 |
| N | -2.6863 | 1.0535 | -0.8429 | -2.9126 | -0.0996 | -1.1058 | -3.2589 | 0.4031 | 0.2571 |
| O | 1.5989 | -0.0159 | -2.6666 | 1.8733 | -0.7585 | -2.7424 | 1.7332 | 0.7196 | -2.6525 |
| C | -0.6049 | -1.8467 | -1.7632 | 0.2323 | -2.8154 | -0.9232 | 0.2324 | -2.1449 | -2.1528 |
| C | 2.1095 | -2.9648 | -2.6738 | 3.1562 | -3.1691 | -1.5581 | 3.0616 | -1.9699 | -2.8630 |
| C | 1.6817 | -4.4012 | -2.3842 | 3.2069 | -4.4125 | -0.6725 | 3.3095 | -3.4592 | -2.6224 |
| C | 3.5899 | -2.8927 | -3.0542 | 4.5538 | -2.8077 | -2.0680 | 4.3607 | -1.2731 | -3.2753 |
| O | 1.3446 | -2.7073 | 1.1595 | 1.9450 | -0.9457 | 1.6403 | 1.8461 | -1.5355 | 1.0305 |
| C | -1.6112 | 4.1338 | 0.2619 | -2.9194 | 3.3693 | -0.6689 | -2.4865 | 3.4414 | -1.2204 |
| C | -0.5483 | 4.8953 | -0.5378 | -2.0398 | 4.5075 | -0.1457 | -1.4722 | 4.5828 | -1.1060 |
| C | 0.7571 | 4.1962 | -0.1556 | -0.6471 | 4.1342 | -0.6524 | -0.1423 | 3.8405 | -0.9840 |
| C | -4.8675 | 0.7849 | 0.2345 | -4.9379 | -0.9640 | -0.0268 | -4.7225 | -1.0869 | -1.0237 |
| C | -5.4021 | 0.2286 | -1.0860 | -4.9922 | -2.0508 | -1.0984 | -5.8920 | -0.1770 | -0.6303 |
| C | -4.5572 | 0.7375 | -2.2464 | -4.3058 | -1.5529 | -2.3629 | -5.4065 | 1.2349 | -0.2960 |
| N | -3.1573 | 0.3624 | -1.9869 | -2.9249 | -1.1873 | -2.0132 | -4.2927 | 1.2713 | 0.6713 |
| C | -1.0074 | -2.6315 | 2.5456 | -0.3205 | -1.2020 | 3.1184 | -0.0840 | -3.1652 | 2.0992 |
| C | -0.3295 | -1.6353 | 3.4513 | -0.3349 | 0.3038 | 3.2281 | 0.8971 | -4.3029 | 2.2083 |
| C | -2.1023 | -3.4633 | 3.0649 | -0.9100 | -2.0112 | 4.1921 | -0.4075 | -2.3324 | 3.2575 |
| O | -0.7987 | -4.0064 | 2.8414 | 0.5064 | -1.9172 | 4.0273 | -1.3033 | -3.3538 | 2.8086 |
| C | 4.7217 | 1.6263 | -1.0824 | 3.9874 | 2.3667 | -1.8038 | 3.9041 | 2.9246 | -0.3308 |
| N | 3.1769 | 1.2354 | 1.4697 | 2.2968 | 2.0567 | 0.7864 | 2.2932 | 1.4766 | 1.8282 |
| C | 2.9113 | 1.7016 | 2.7174 | 2.2350 | 2.9616 | 1.7943 | 2.8843 | 1.9736 | 2.9364 |
| C | 3.3907 | 0.8533 | 3.8672 | 2.6843 | 2.4551 | 3.1435 | 3.3684 | 0.9446 | 3.9295 |
| O | 2.3264 | 2.7644 | 2.8857 | 1.8676 | 4.1180 | 1.6202 | 3.0315 | 3.1765 | 3.1295 |
| H | 2.8756 | 2.9758 | 0.3957 | 1.8345 | 3.4192 | -0.7049 | 1.7411 | 3.3406 | 1.1597 |
| H | 2.6993 | 1.7707 | -1.8245 | 2.1191 | 1.5529 | -2.5368 | 2.0282 | 2.6032 | -1.3469 |
| H | 2.5517 | -2.3849 | -0.6510 | 3.2971 | -1.7180 | 0.0156 | 3.2408 | -1.4673 | -0.7815 |
| H | -1.6439 | -3.0965 | 0.5703 | -0.5224 | -2.8617 | 1.7882 | -0.5173 | -3.4562 | 0.0398 |
| H | -0.9759 | -0.2491 | 0.6600 | -0.9842 | -0.4368 | 0.2369 | -1.2131 | -0.7218 | 0.9201 |
| H | -1.3381 | 2.1387 | 1.1351 | -2.1347 | 1.7429 | 0.5521 | -2.3354 | 2.6416 | 0.7956 |
| H | -3.0095 | 1.2091 | 1.2107 | -3.5847 | 0.4756 | 0.7766 | -4.1466 | -1.2632 | 1.0269 |
| H | -1.5151 | -2.3351 | -1.4233 | 0.6571 | -3.5613 | -1.5905 | 0.1231 | -1.2607 | -2.7819 |
| H | -0.7728 | -0.7693 | -1.8102 | -0.4914 | -3.3167 | -0.2846 | 0.5263 | -2.9962 | -2.7690 |
| H | -0.3835 | -2.2118 | -2.7648 | -0.2818 | -2.0541 | -1.5147 | -0.7311 | -2.3517 | -1.7040 |
| H | 1.5269 | -2.5718 | -3.5122 | 2.5225 | -3.3583 | -2.4290 | 2.3324 | -1.8393 | -3.6677 |
| H | 1.8996 | -5.0399 | -3.2425 | 3.6401 | -5.2497 | -1.2234 | 3.6504 | -3.9390 | -3.5418 |
| H | 0.6132 | -4.4696 | -2.1702 | 2.2164 | -4.7102 | -0.3231 | 2.4139 | -3.9796 | -2.2777 |
| H | 2.2271 | -4.7942 | -1.5206 | 3.8320 | -4.2253 | 0.2059 | 4.0868 | -3.5913 | -1.8638 |
| H | 3.7811 | -3.4932 | -3.9453 | 4.9695 | -3.6358 | -2.6447 | 4.7789 | -1.7547 | -4.1609 |
| H | 3.9067 | -1.8683 | -3.2692 | 4.5361 | -1.9269 | -2.7151 | 4.2011 | -0.2188 | -3.5124 |

| | | | | | | | | | |
|---|---|---|---|---|---|---|---|---|---|
| H | 4.2137 | -3.2798 | -2.2432 | 5.2282 | -2.6051 | -1.2309 | 5.1028 | -1.3388 | -2.4739 |
| H | -1.6186 | 4.4653 | 1.3018 | -3.8733 | 3.2777 | -0.1511 | -3.5084 | 3.7412 | -0.9922 |
| H | -2.6174 | 4.2439 | -0.1427 | -3.1071 | 3.4980 | -1.7377 | -2.4575 | 3.0089 | -2.2246 |
| H | -0.7285 | 4.7768 | -1.6065 | -2.3560 | 5.4861 | -0.5035 | -1.4867 | 5.2561 | -1.9616 |
| H | -0.5244 | 5.9574 | -0.2988 | -2.0453 | 4.5167 | 0.9467 | -1.6597 | 5.1626 | -0.1995 |
| H | 1.1875 | 4.6107 | 0.7605 | 0.1503 | 4.5212 | -0.0200 | 0.6237 | 4.4178 | -0.4657 |
| H | 1.4974 | 4.2383 | -0.9544 | -0.4992 | 4.4633 | -1.6843 | 0.2297 | 3.5577 | -1.9729 |
| H | -5.0666 | 1.8595 | 0.2867 | -5.5603 | -0.1178 | -0.3315 | -5.0486 | -2.1208 | -1.1346 |
| H | -5.3573 | 0.3111 | 1.0857 | -5.3103 | -1.3310 | 0.9292 | -4.2777 | -0.7663 | -1.9709 |
| H | -5.3615 | -0.8624 | -1.0706 | -4.4784 | -2.9498 | -0.7460 | -6.6266 | -0.1248 | -1.4366 |
| H | -6.4431 | 0.5286 | -1.2202 | -6.0295 | -2.3161 | -1.3102 | -6.3973 | -0.6013 | 0.2443 |
| H | -4.6597 | 1.8260 | -2.3540 | -4.8501 | -0.6967 | -2.7840 | -6.2104 | 1.8437 | 0.1206 |
| H | -4.8468 | 0.2632 | -3.1848 | -4.2447 | -2.3367 | -3.1187 | -5.0513 | 1.7311 | -1.2054 |
| H | -2.5585 | 0.6697 | -2.7499 | -2.4228 | -0.8552 | -2.8331 | -4.5985 | 0.9737 | 1.5942 |
| H | -0.3945 | -1.9787 | 4.4846 | -0.0512 | 0.5948 | 4.2406 | 0.5843 | -5.1293 | 1.5646 |
| H | 0.7184 | -1.5189 | 3.1763 | 0.3590 | 0.7559 | 2.5175 | 1.8938 | -3.9802 | 1.9050 |
| H | -0.8255 | -0.6651 | 3.3767 | -1.3390 | 0.6846 | 3.0287 | 0.9403 | -4.6596 | 3.2376 |
| H | -2.8723 | -3.8151 | 2.3875 | -1.3412 | -2.9782 | 3.9576 | -0.7621 | -1.3172 | 3.1102 |
| H | -2.3814 | -3.3659 | 4.1090 | -1.3009 | -1.5067 | 5.0700 | 0.0842 | -2.5369 | 4.2029 |
| H | 5.1000 | 1.1972 | -2.0107 | 4.6146 | 1.8047 | -2.4965 | 4.5085 | 2.7738 | -1.2260 |
| H | 4.9027 | 2.7022 | -1.0948 | 3.8089 | 3.3595 | -2.2192 | 3.7450 | 3.9953 | -0.1908 |
| H | 5.2622 | 1.1873 | -0.2429 | 4.5129 | 2.4689 | -0.8522 | 4.4434 | 2.5319 | 0.5332 |
| H | 3.4303 | 0.2633 | 1.3539 | 2.4381 | 1.0736 | 1.0091 | 2.1617 | 0.4740 | 1.7227 |
| H | 2.5270 | 0.5706 | 4.4706 | 1.9603 | 2.7724 | 3.8940 | 4.4452 | 1.0642 | 4.0563 |
| H | 4.0522 | 1.4567 | 4.4897 | 3.6441 | 2.9170 | 3.3823 | 2.8937 | 1.1430 | 4.8911 |
| H | 3.9164 | -0.0432 | 3.5426 | 2.7905 | 1.3706 | 3.1685 | 3.1505 | -0.0770 | 3.6201 |

Table S15. Lowest-energy conformers optimized at the M062X/6-311+G(d, p) level of 1d in DMSO with relative energies < 2.5 kcal/mol

| **1d_C1** ΔG 0.00 kcal/mol Boltzman pop. 93.75% | **1d_C2** ΔG 1.62 kcal/mol Boltzman pop. 6.09% |
|---|---|
| 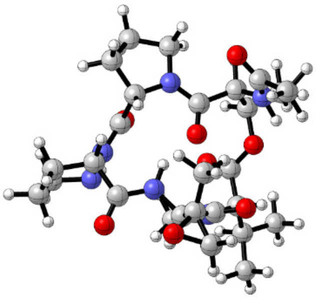 | 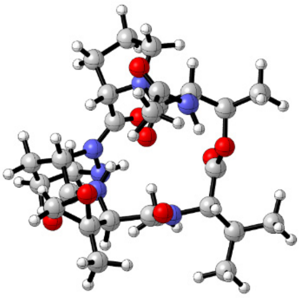 |

Table S16. Atomic coordinates for the lowest-energy conformers of 1d (1d_C1–1d_C2)

| Atoms | **1d_C1** | | | **1d_C2** | | |
|---|---|---|---|---|---|---|
| | x | y | z | x | y | z |
| C | 2.4718 | 2.0716 | 0.2140 | 2.3391 | 2.2131 | 0.4510 |
| C | 3.0550 | 1.5482 | -1.1065 | 3.0751 | 1.8145 | -0.8362 |
| O | 2.9678 | 0.1084 | -1.0590 | 3.0592 | 0.3723 | -0.8971 |
| C | 2.0658 | -0.5147 | -1.8316 | 2.1986 | -0.2300 | -1.7315 |
| O | 0.5763 | 0.6449 | 0.5112 | 0.5084 | 0.6772 | 0.5186 |
| C | 0.9659 | 1.7934 | 0.2802 | 0.8503 | 1.8530 | 0.3639 |
| C | 1.9394 | -1.9931 | -1.4753 | 2.1410 | -1.7366 | -1.4965 |
| N | 0.6300 | -2.1867 | -0.8432 | 0.8221 | -2.0624 | -0.9439 |
| O | -3.5840 | -2.0657 | 0.7826 | -3.4558 | -2.2925 | 0.6543 |
| O | -1.4468 | 2.1183 | -2.1144 | -1.5216 | 2.0730 | -2.1256 |
| C | 0.5990 | -2.3430 | 0.5079 | 0.7387 | -2.2952 | 0.3962 |
| C | -0.7837 | -2.3381 | 1.1980 | -0.6791 | -2.4908 | 0.9845 |
| N | -1.5170 | -1.1286 | 0.8752 | -1.4381 | -1.2724 | 0.7930 |
| C | -2.8524 | -1.0934 | 0.6929 | -2.7718 | -1.2829 | 0.5938 |
| C | -1.8513 | 1.8166 | -0.9948 | -1.9173 | 1.7333 | -1.0141 |
| C | -1.3207 | 2.5623 | 0.2326 | -1.4666 | 2.5053 | 0.2265 |
| N | 0.1138 | 2.8012 | 0.0761 | -0.0404 | 2.8203 | 0.1316 |
| C | -3.3883 | 0.3255 | 0.3938 | -3.3682 | 0.1182 | 0.3355 |
| N | -2.7699 | 0.8367 | -0.8265 | -2.7649 | 0.6866 | -0.8670 |
| O | 1.4316 | 0.0252 | -2.7001 | 1.5560 | 0.3471 | -2.5689 |
| C | -0.5620 | -2.0193 | -1.6714 | -0.3403 | -1.8801 | -1.8105 |
| C | 2.1864 | -2.8757 | -2.7064 | 2.4881 | -2.4973 | -2.7848 |
| C | 1.9030 | -4.3383 | -2.3737 | 2.2773 | -3.9966 | -2.5897 |
| C | 3.6319 | -2.6831 | -3.1701 | 3.9372 | -2.1874 | -3.1667 |
| O | 1.6086 | -2.5027 | 1.1794 | 1.7213 | -2.3626 | 1.1190 |
| C | -1.9068 | 3.9832 | 0.3125 | -2.1321 | 3.8930 | 0.2847 |
| C | -0.9254 | 4.8253 | -0.5106 | -1.1475 | 4.7998 | -0.4623 |
| C | 0.4381 | 4.2151 | -0.1809 | 0.2202 | 4.2586 | -0.0455 |
| C | -4.9142 | 0.3775 | 0.2636 | -4.8939 | 0.1057 | 0.1939 |
| C | -5.4112 | -0.2090 | -1.0583 | -5.3529 | -0.4724 | -1.1457 |
| C | -4.6238 | 0.3852 | -2.2183 | -4.5790 | 0.1758 | -2.2860 |
| N | -3.1953 | 0.1253 | -1.9760 | -3.1430 | -0.0269 | -2.0315 |
| C | -0.6025 | -2.4896 | 2.7025 | -0.5926 | -2.9281 | 2.4435 |
| C | -0.1195 | -1.2732 | 3.4475 | -0.0395 | -4.3131 | 2.6713 |
| C | -0.3957 | -3.8384 | 3.2298 | -1.5201 | -2.3634 | 3.4323 |
| O | -1.6685 | -3.1985 | 3.3382 | -0.1656 | -1.9174 | 3.3478 |
| C | 4.5107 | 1.9232 | -1.3014 | 4.5229 | 2.2623 | -0.8537 |
| N | 3.1274 | 1.4253 | 1.3308 | 2.9232 | 1.5167 | 1.5791 |
| C | 2.8480 | 1.8394 | 2.5941 | 2.4296 | 1.7567 | 2.8251 |
| C | 3.4517 | 1.0312 | 3.7141 | 2.8486 | 0.7841 | 3.8964 |
| O | 2.1665 | 2.8365 | 2.7966 | 1.6756 | 2.6973 | 3.0353 |
| H | 2.6606 | 3.1437 | 0.2744 | 2.4581 | 3.2866 | 0.6011 |
| H | 2.4461 | 1.9109 | -1.9352 | 2.5355 | 2.2091 | -1.6974 |
| H | 2.6848 | -2.2301 | -0.7186 | 2.8696 | -1.9932 | -0.7299 |
| H | -1.3677 | -3.1937 | 0.8437 | -1.1861 | -3.2956 | 0.4417 |
| H | -0.9605 | -0.2847 | 0.7414 | -0.9128 | -0.4026 | 0.7021 |
| H | -1.4754 | 2.0021 | 1.1525 | -1.6267 | 1.9369 | 1.1400 |
| H | -3.0952 | 0.9631 | 1.2259 | -3.1074 | 0.7412 | 1.1890 |
| H | -1.4012 | -2.5689 | -1.2502 | -0.6583 | -0.8362 | -1.8266 |
| H | -0.8348 | -0.9678 | -1.7714 | -0.0767 | -2.1834 | -2.8221 |
| H | -0.3610 | -2.4265 | -2.6610 | -1.1696 | -2.5024 | -1.4830 |
| H | 1.5275 | -2.5495 | -3.5164 | 1.8446 | -2.1391 | -3.5937 |
| H | 2.1269 | -4.9714 | -3.2345 | 2.5752 | -4.5390 | -3.4892 |
| H | 0.8588 | -4.4962 | -2.0962 | 1.2335 | -4.2353 | -2.3767 |
| H | 2.5293 | -4.6664 | -1.5383 | 2.8856 | -4.3607 | -1.7559 |
| H | 3.8262 | -3.2834 | -4.0606 | 4.2007 | -2.6993 | -4.0938 |
| H | 3.8450 | -1.6399 | -3.4194 | 4.0990 | -1.1169 | -3.3204 |

| | | | | | | |
|---|---|---|---|---|---|---|
| H | 4.3310 | -2.9970 | -2.3894 | 4.6221 | -2.5271 | -2.3842 |
| H | -1.9055 | 4.3036 | 1.3559 | -2.2155 | 4.1981 | 1.3293 |
| H | -2.9302 | 4.0257 | -0.0607 | -3.1303 | 3.8868 | -0.1535 |
| H | -1.1327 | 4.7070 | -1.5741 | -1.2753 | 4.6833 | -1.5388 |
| H | -0.9671 | 5.8836 | -0.2577 | -1.2645 | 5.8508 | -0.2025 |
| H | 0.8756 | 4.6574 | 0.7187 | 0.5539 | 4.6908 | 0.9027 |
| H | 1.1417 | 4.3073 | -1.0082 | 0.9826 | 4.4239 | -0.8067 |
| H | -5.2052 | 1.4303 | 0.3281 | -5.2314 | 1.1429 | 0.2796 |
| H | -5.3545 | -0.1447 | 1.1136 | -5.3178 | -0.4548 | 1.0279 |
| H | -5.2759 | -1.2923 | -1.0562 | -5.1737 | -1.5491 | -1.1642 |
| H | -6.4754 | 0.0020 | -1.1801 | -6.4235 | -0.3025 | -1.2748 |
| H | -4.8184 | 1.4627 | -2.3088 | -4.8181 | 1.2457 | -2.3597 |
| H | -4.8813 | -0.0985 | -3.1612 | -4.8061 | -0.3010 | -3.2402 |
| H | -2.6320 | 0.4925 | -2.7395 | -2.5883 | 0.3760 | -2.7832 |
| H | 0.8001 | -0.8981 | 2.9954 | 0.0266 | -4.5114 | 3.7416 |
| H | 0.0670 | -1.5222 | 4.4927 | 0.9518 | -4.4140 | 2.2302 |
| H | -0.8751 | -0.4851 | 3.4030 | -0.7047 | -5.0548 | 2.2227 |
| H | 0.1414 | -3.9603 | 4.1646 | -1.7525 | -2.9556 | 4.3118 |
| H | -0.3652 | -4.6824 | 2.5470 | -2.2611 | -1.6296 | 3.1395 |
| H | 4.8711 | 1.5142 | -2.2457 | 4.9948 | 1.9399 | -1.7822 |
| H | 4.6108 | 3.0092 | -1.3292 | 4.5720 | 3.3507 | -0.7953 |
| H | 5.1253 | 1.5331 | -0.4889 | 5.0718 | 1.8393 | -0.0114 |
| H | 3.4598 | 0.4807 | 1.1893 | 3.3036 | 0.5968 | 1.3943 |
| H | 2.6782 | 0.8281 | 4.4545 | 2.8545 | 1.2904 | 4.8595 |
| H | 4.2267 | 1.6330 | 4.1921 | 3.8271 | 0.3492 | 3.6951 |
| H | 3.8901 | 0.0949 | 3.3718 | 2.1092 | -0.0210 | 3.9284 |

**Determination of the structure of aglomycin B (2)**

Aglomycin B (**2**) was obtained as a white powder. Its molecular formula of $C_{10}H_8ClN_2O_2S$ was deduced from the (+)-HRESIMS ion at *m/z* 254.9985 $[M+H]^+$, with nine double bond equivalents (DBEs). The $^1H$ and $^{13}C$ NMR data (Table S18) recorded in DMSO-$d_6$ indicated the presence of an ortho-trisubstituted phenyl group [$\delta_C$ 115.1 – 142.4, $\delta_H$ 7.40, d (7.8); 6.67, dt (1.8, 7.8); 7.64, d (8.4)] and a 2,4-disubstituted thiazole ring ($\delta_C$ 167.7, 148.6, 126.2; $\delta_H$ 8.39, s). Analysis of the $^1H$-$^1H$ COSY spectrum revealed one independent spin coupling systems H-4′/H-5′/H-6′ (Figure S27). The HMBC correlations from H-5 to C-2, C-4 and C-1″, H-4′ to C-2″ and C-3″, H-6′ to C-2, C-1″ and C-2″ and $NH_2$ to C-1″ and C-3″, combined with characteristic isotope distributions of the chlorine atom in MS spectrum (Figure S28), the structure of **2** was deduced. (Figure S27).

Table S17. $^1H$ and $^{13}C$ NMR data for compound 4 in DMSO-$d_6$.

| No. | $\delta_C$, type | $\delta_H$, multi. (*J* in Hz) | $^1H$-$^1H$ COSY | HMBC |
|---|---|---|---|---|
| 2 | 167.7, C | | | |
| 4 | 148.6, C | | | |
| 5 | 126.2, CH | 8.38, s | | 1'', 4, 2 |
| 1' | 115.1, C | | | |
| 2' | 142.4, C | | | |
| 3' | 119.2, C | | | |
| 4' | 131.0, CH | 7.40, dd (1.8, 7.8) | 5' | 1', 2', 3', 6' |
| 5' | 116.1, CH | 6.67, dt (1.8, 7.8) | 4', 6' | 1', 2', 3', 4', 6' |
| 6' | 128.0, CH | 7.64, dd (1.8, 7.8) | 5' | 2, 1', 2', 3', 4' |
| 1'' | 162.3, C | | | |
| 2'-$NH_2$ | | 7.32, s | | 1', 3' |

$^1H$ and $^{13}C$ NMR data were recorded at 600 and 150 MHz, respectively. The assignments were based on 2D NMR ($^1H$-$^1H$ COSY, HSQC, HMBC and ROESY) experiments.

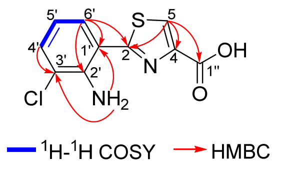

**Figure S27. The structure and the key 2D NMR correlations for aglomycin B (2).**

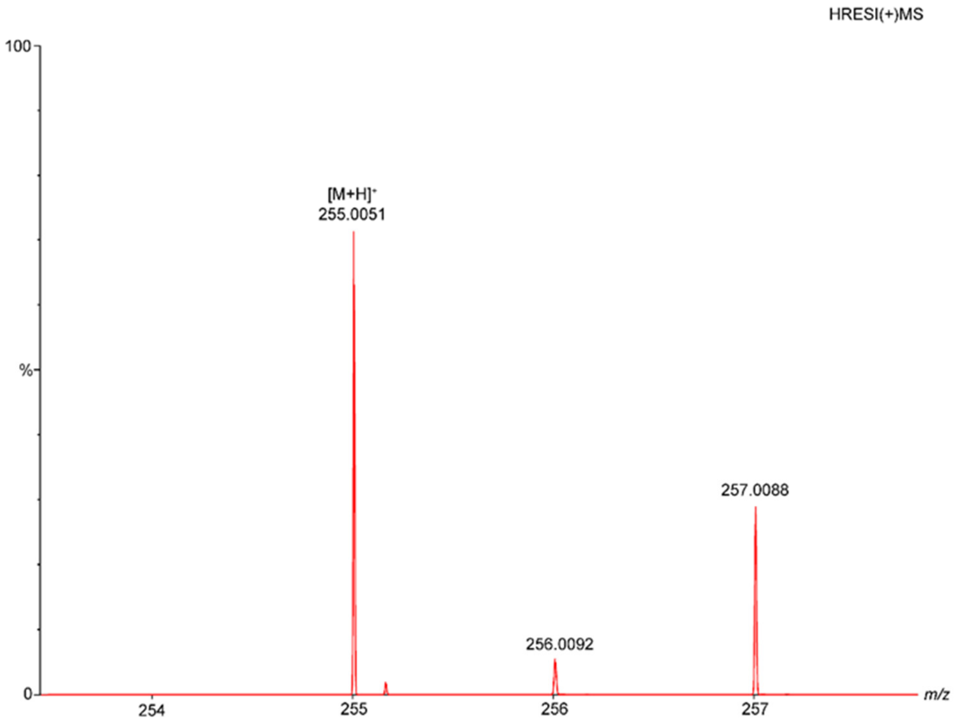

**Figure S28. HRESI(+)MS spectrum of compound 2**

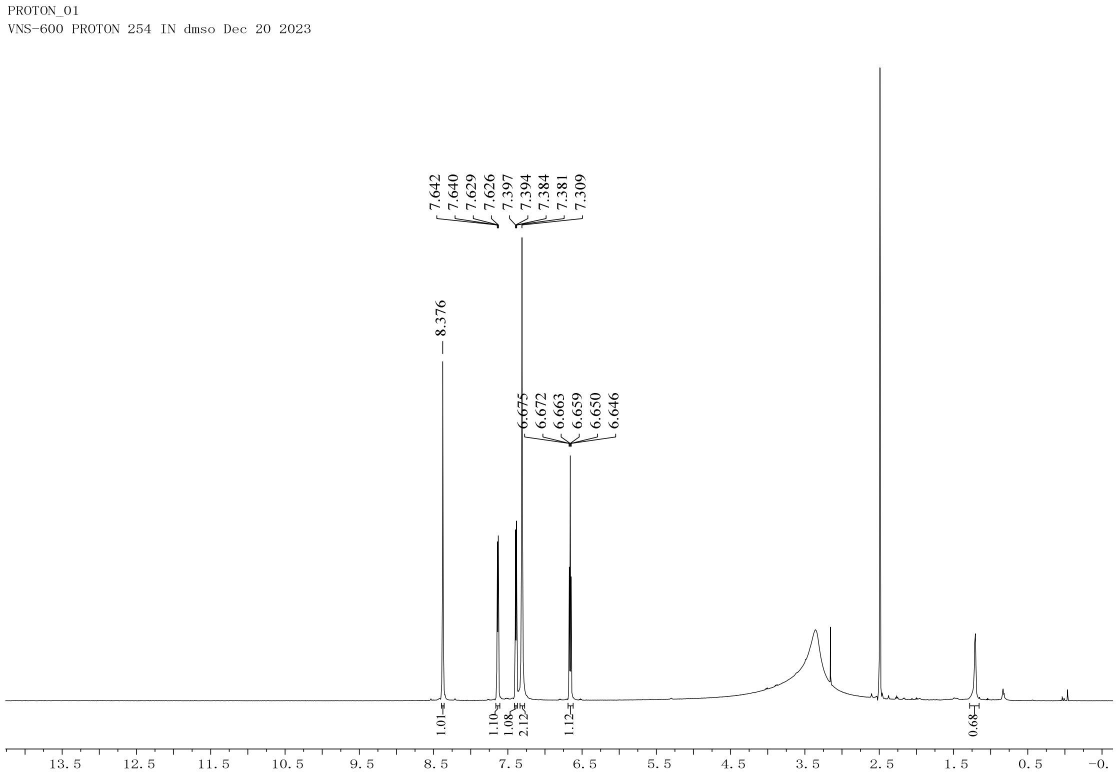

**Figure S29. $^{1}$H NMR spectrum of compound 2 in DMSO-$d_6$ (600 MHz).**

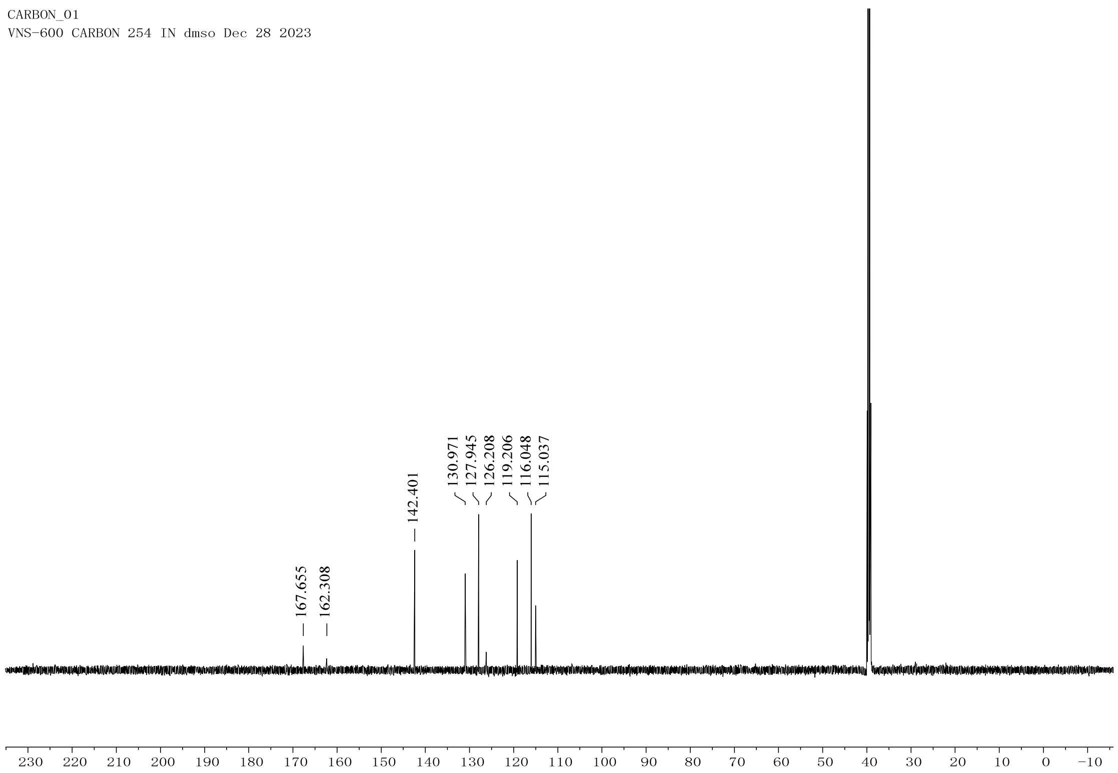

**Figure S30.** $^{13}C$ **NMR spectrum of compound 2 in DMSO-** $d_6$ **(150 MHz).**

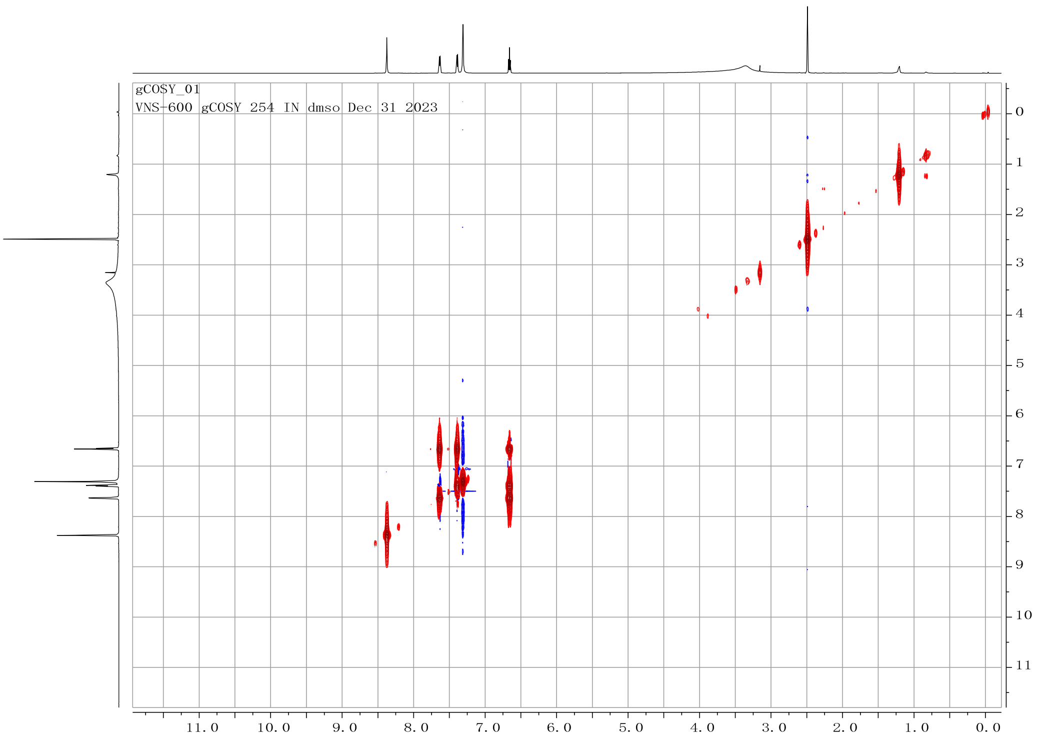

**Figure S31.** $^{1}H$-$^{1}H$ **COSY spectrum of compound 2 in DMSO-** $d_6$

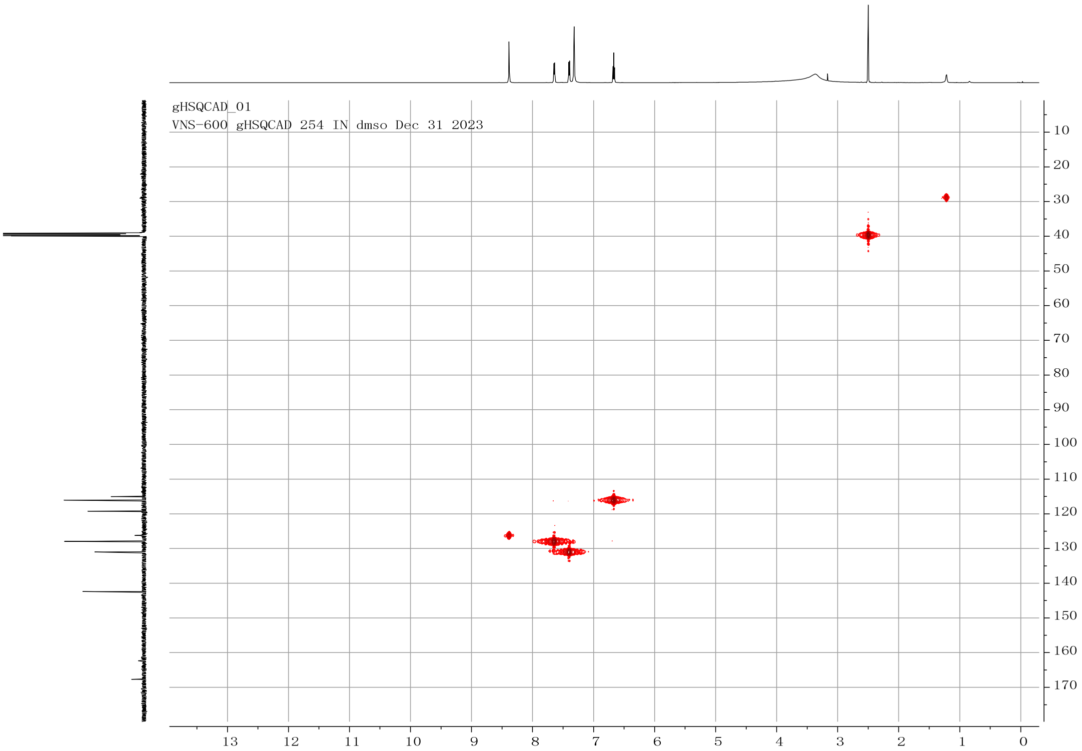

**Figure S32. HSQC spectrum of compound 2 in DMSO-$d_6$**

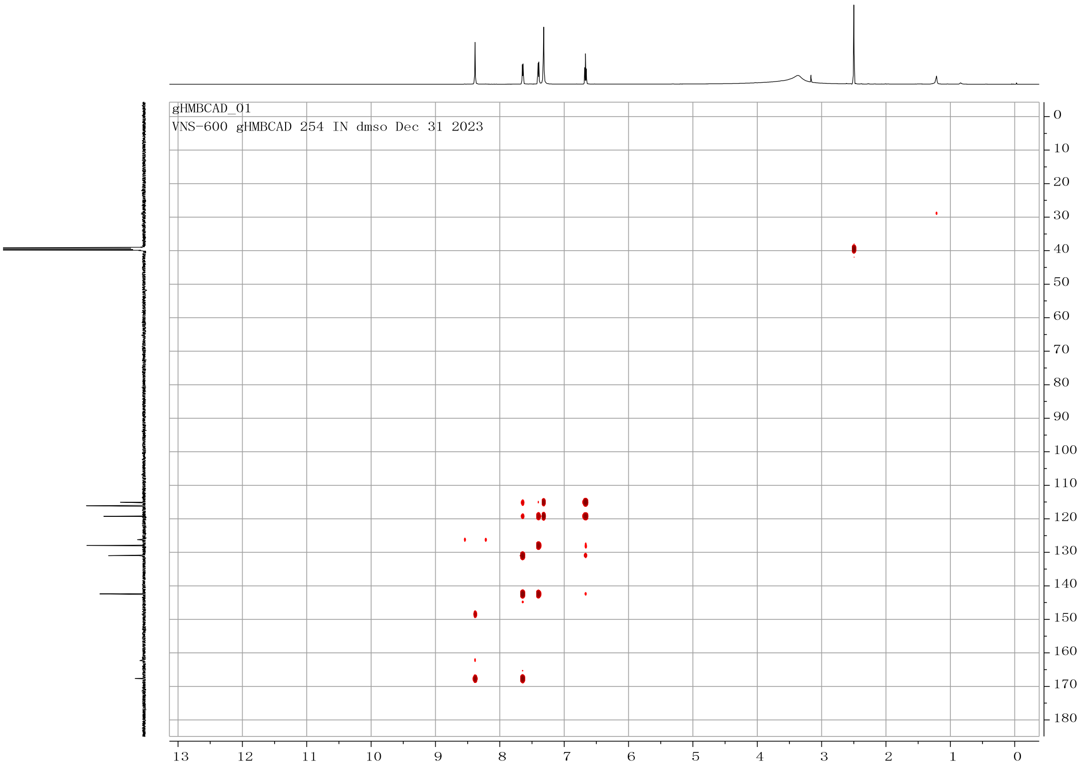

**Figure S33. HMBC spectrum of compound 2 in DMSO-$d_6$**

**Structural characterization of aglomycins C and D**

Additionally, due to the low yields of other aglomycin analogues, their structures were only tentatively identified through MS (Figure S34). HRESI(+)MS of aglomycin C (**3**) revealed a sodium adduct ion at *m/z* 779.2731, 16 mass units less than that of **1**. Its $MS^2$ spectrum displayed diagnostic fragment ions at *m/z* 666, 569, 457 and 360, corresponding to sequential neutral losses of 113, 97, 112, and 97 Da, respectively. This fragmentation pattern closely resembled that of **1**, except for the second neutral loss (97 Da in **3** vs. 113 Da in **1**), suggestive of the replacement of epoxy-$Val^4$ (loss of 113 Da) in **1** with either dehydro-$Val^4$ or $Pro^4$ (loss of 97 Da) in **3**. However, based on the following biosynthetic assembly line analysis and the substrate specificity of nonribosomal peptide synthetase (NRPS) A domains, dehydro-$Val^4$ appears to be the most plausible modification in compound **3**, rather than $Pro^4$. Similarly, aglomycin D (**4**) ($[M+Na]^+$ at *m/z* 813.2811) was assumed to be a hydrolysis product of **1**, bearing a 3,4-dihydroxy-Val (diHO-$Val^4$) residue in place of the epoxy-$Val^4$ found in **1**.

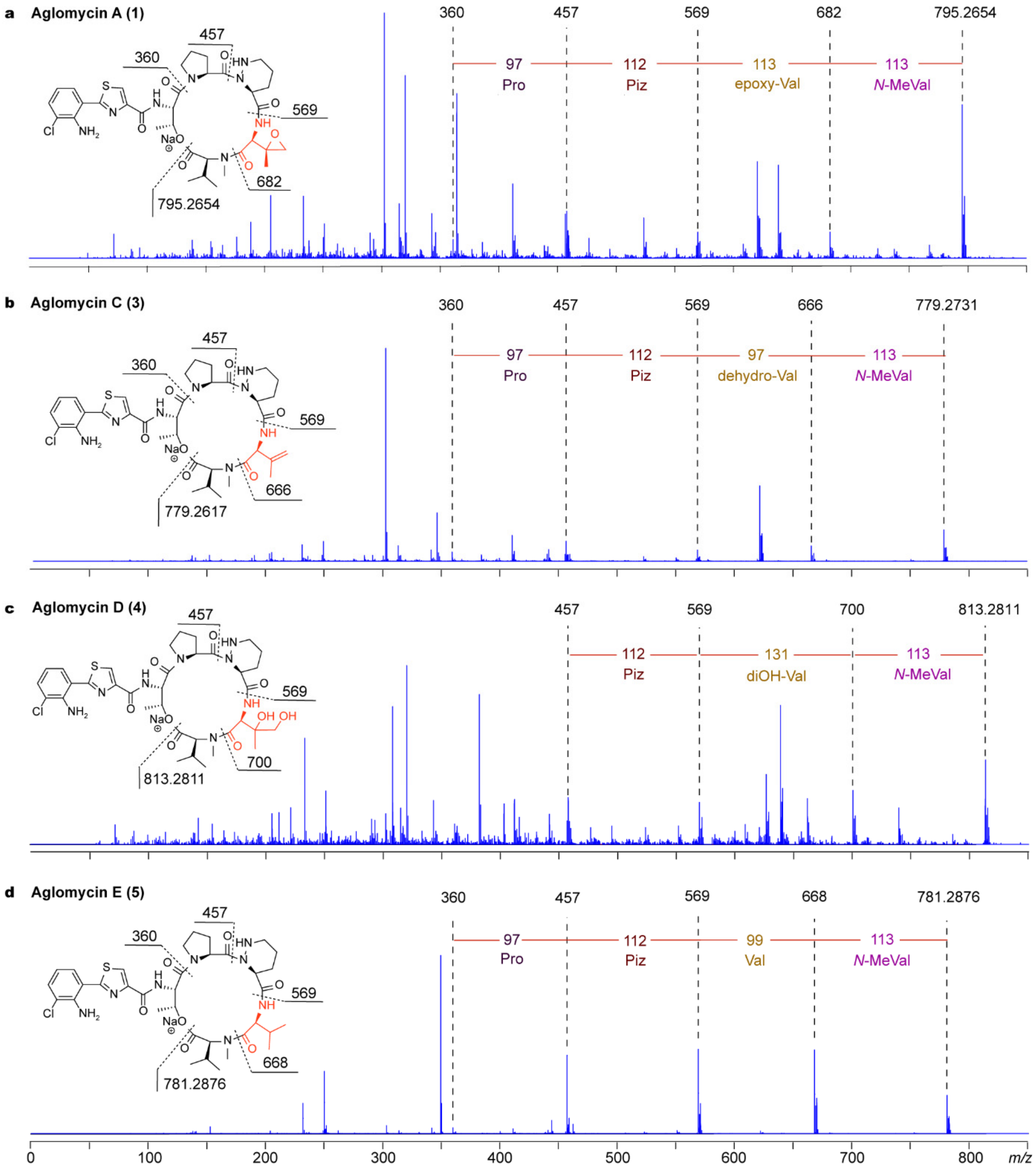

**Figure S34. Structural identification of aglomycins A, C, D and E through MS$^2$ spectra analysis of the sodium adducts. a**, Aglomycin A (**1**), with *m/z* 795.2654, produced diagnostic fragment ions at *m/z* 682, 569, 457 and 360 with a mass difference of 113-113-112-97, allowing to build up the fragment of (*N*-MeVal)-(epoxy-Val)-Piz. **b**, Aglomycin C (**3**), with *m/z* 779.2731, yielded characteristic fragment ions at *m/z* 666, 569, 457 and 360 with a mass difference of 113-97-112-97, enabling the deduction of the fragment of (*N*-MeVal)-(dehydro-Val)-Piz-Pro. **c**, Aglomycin D (**4**), with *m/z* 813.2811, generated diagnostic fragment ions at *m/z* 700, 569 and 457 with a mass difference of 113-131-112, supporting the proposed fragment of (*N*-MeVal)-(diOH-Val)-Piz. **d**, Aglomycin E (**5**), observed at *m/z* 781.2876, displayed fragment ions at *m/z* 668, 569, 457 and 360, showing neutral losses of 113-99-112-97, from which the fragment (*N*-MeVal)-Val-Piz-Pro was deduced.

**Characterization of aglomycins A-E**

Aglomycin A (**1**): white powder; $[\alpha]_D^{20}$ +3.03 (*c* 0.001, MeOH); UV (MeOH) $\lambda_{max}$ (log $\varepsilon$): 286 (0.51), 366 (0.51) nm; $^1$H-NMR (DMSO-$d_6$, 600 MHz) and $^{13}$C NMR (DMSO-$d_6$, 150 MHz), Table S5; HRESIMS: *m/z* 773.2849 $[M+H]^+$ (Calcd for $C_{35}H_{45}ClN_8O_8S$, 773.2848, 0.1 ppm).

Aglomycin B (**2**): white powder; UV (MeOH) $\lambda_{max}$ (log $\varepsilon$): 286 (0.28), 366 (0.24) nm; $^1$H-NMR (DMSO-$d_6$, 600 MHz) and $^{13}$C NMR (DMSO-$d_6$, 150 MHz), Table S17; HRESIMS: *m/z* 254.9985 $[M+H]^+$ (Calcd for $C_{10}H_8ClN_2O_2S$, 254.9995, 3.9 ppm).

Aglomycin C (**3**): HRESIMS: *m/z* 779.2731 $[M+Na]^+$ (Calcd for $C_{35}H_{45}ClN_8O_7SNa$, 779.2718, 1.6 ppm); diagnostic fragment ions MS$^2$ spectrum: *m/z* 666, 569, 457, and 360.

Aglomycin D (**4**): HRESIMS: *m/z* 791.2958 $[M+H]^+$ (Calcd for $C_{35}H_{48}ClN_8O_9S$, 791.2953, 0.6 ppm); diagnostic fragment ions MS$^2$ spectrum: *m/z* 700, 569, and 457.

Aglomycin E (**5**): HRESIMS: *m/z* 781.2876 $[M+Na]^+$ (Calcd for $C_{35}H_{47}ClN_8O_7SNa$, 781.2875, 0.1 ppm); diagnostic fragment ions MS$^2$ spectrum: *m/z* 668, 569, 457 and 360.

#### Identification of algomycin biosynthesis gene cluster

The complete genome of strain 11-23 was sequenced using PacBio HiFi technology, assembled with Hifiasm, circularized using Circlator v1.5.5, and polished with Illumina data using Pilon v1.22. Secondary metabolite biosynthetic gene clusters (BGCs) were analyzed with AntiSMASH[45]. The complete aglomycin BGC has been deposited into GenBank under accession number PV571899. The identities and similarities of the encoded proteins to their homologs were examined using BlastP.

To confirm the BGC, genes *aglA*, *aglB*, and *aglG* were inactivated by homologous recombination. Upstream and downstream homologous fragments were amplified by PCR, sequenced, and cloned into the temperature-sensitive plasmid pKC1139 to create in-frame deletion plasmids. These plasmids were introduced into *Streptomyces* sp. 11-23 by conjugation with *E. coli* ET12567/pUZ8002. After conjugation, conjugants were selected on MS medium with aztreonam and apramycin. Single-crossover recombinants were cultivated at 37 °C, followed by seven rounds of propagation without antibiotics at 28 °C to obtain double-crossover mutants Δ*aglA*, Δ*aglB*, and Δ*aglG*, confirmed by PCR. For gene complementation, coding regions of *aglB* and *aglG* were amplified and cloned into pICLset to obtain complement plasmids, which were introduced into Δ*aglB* and Δ*aglG* strains by conjugation. Correct recombinants were confirmed by PCR. These strains were selected by their apramycin-resistant phenotype and verified by PCR. The metabolites of engineered strained were analyzed using the same method as high-throughput mining of halogenates as described above.

Table S18. Deduced functions of genes in the aglomycin biosynthetic gene cluster (*agl*)

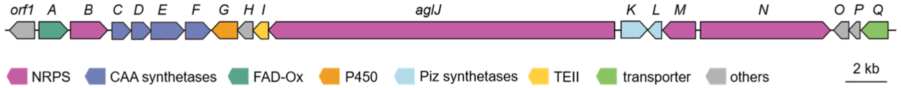

| Genes | Size (aa) | Proposed Function | Protein homologue and origin | ID/SI[a] |
|---|---|---|---|---|
| *orf1* | 460 | putative hypothetical protein | P94572.1, *Bacillus subtilissubsp.* subtilis str. 168 | 33/52 |
| *aglA* | 461 | putative FAD-dependent oxygenase | P43485.2， *Streptomyces lavendulae* | 51/65 |
| *aglB* | 589 | putative non-ribosomal peptide synthetase (A-T) | Q70LM5.1, *Brevibacillus parabrevis* | 44/60 |
| *aglC* | 301 | putative tryptophan 2,3-dioxygenase | A4IT59.1, *Geobacillus thermodenitrificans* NG80-2 | 41/60 |
| *aglD* | 309 | putative kynurenine formamidase | P96402.1， *Mycobacterium tuberculosis* H37Rv | 28/41 |
| *aglE* | 530 | putative tryptophan halogenase | Q8KHZ8.1, Lentzea aerocolonigenes | 63/77 |
| *aglF* | 401 | putative kynureninase | P83788.1, *Pseudomonas fluorescens* | 42/63 |
| *aglG* | 411 | putative cytochrome P450 | Q9L9F9.1, *Streptomyces niveus* | 32/50 |
| *aglH* | 249 | putative hypothetical protein | P74395.2, *Synechocystis* sp. PCC 6803 substr. Kazusa | 29/51 |
| *aglI* | 248 | putative thioesterase (TEII) | P14686.1, *Aneurinibacillus migulanus* | 33/49 |
| *aglJ* | 5525 | putative nonribosomal peptide synthetase (NRPS) | O30409.1, *Brevibacillus parabrevis* | 34/51 |
| *aglK* | 419 | putative L-ornithine N -monooxygenase | Q51548.2, *Pseudomonas aeruginosa* PAO1 | 35/52 |
| *aglL* | 221 | putative N−N bond formation enzyme | WMQ71277.1, *Streptomyces* sp. GB16 | 62/71 |
| *aglM* | 525 | putative AMP-binding ligase (NRPS) | P80436.2, *Streptomyces triostinicus* | 71/81 |
| *aglN* | 2073 | putative nonribosomal peptide synthetase (NRPS) | P48633.1, *Yersinia enterocolitica subsp.* enterocolitica 8081 | 38/52 |
| *aglO* | 243 | putative diacetylchitobiose deacetylase | Q6F4N1.1, *Thermococcus kodakarensis* KOD1 | 29/47 |
| *aglP* | 179 | putative DUF6879 family protein | Q9AW48.1, *Guillardia theta* | 25/44 |
| *aglQ* | 428 | putative MFS transporter | O51798.1, *Cupriavidus pinatubonensis* JMP134 | 29/50 |

[a]ID = Closest Protein Identity (%), SI = Closest Protein Similarity (

Table S19. Key amino acid sequences and possible recognition substrates in A domain of *agl* cluster

| Module | 10 AA signature | Stachelhaus | code match | AA in aglomycin A |
|---|---|---|---|---|
| AglN-M2 | DLFNLSLIWK | Cys | 100% (strong) | Cys |
| AglJ-M3 | DFWNIGMVHK | Thr | 100% (strong) | Thr |
| AglJ-M4 | DVQFCANVVK | Pro | 80% (moderate) | Pro |
| AglJ-M5 | DVFSVAAYAK | Leu | 100% (strong) | Piz |
| AglJ-M7 | DAYWAGIVNK | Val | 70% (weak) | Val |
| AglB-M | DALWLGGTFK | Val | 100% (strong) | Val |
| AglM-M | TAPSQGWLAK | diOH-Bz, Sal | 70% (weak) | 2-Amino-3-chlorobenzoic acid |

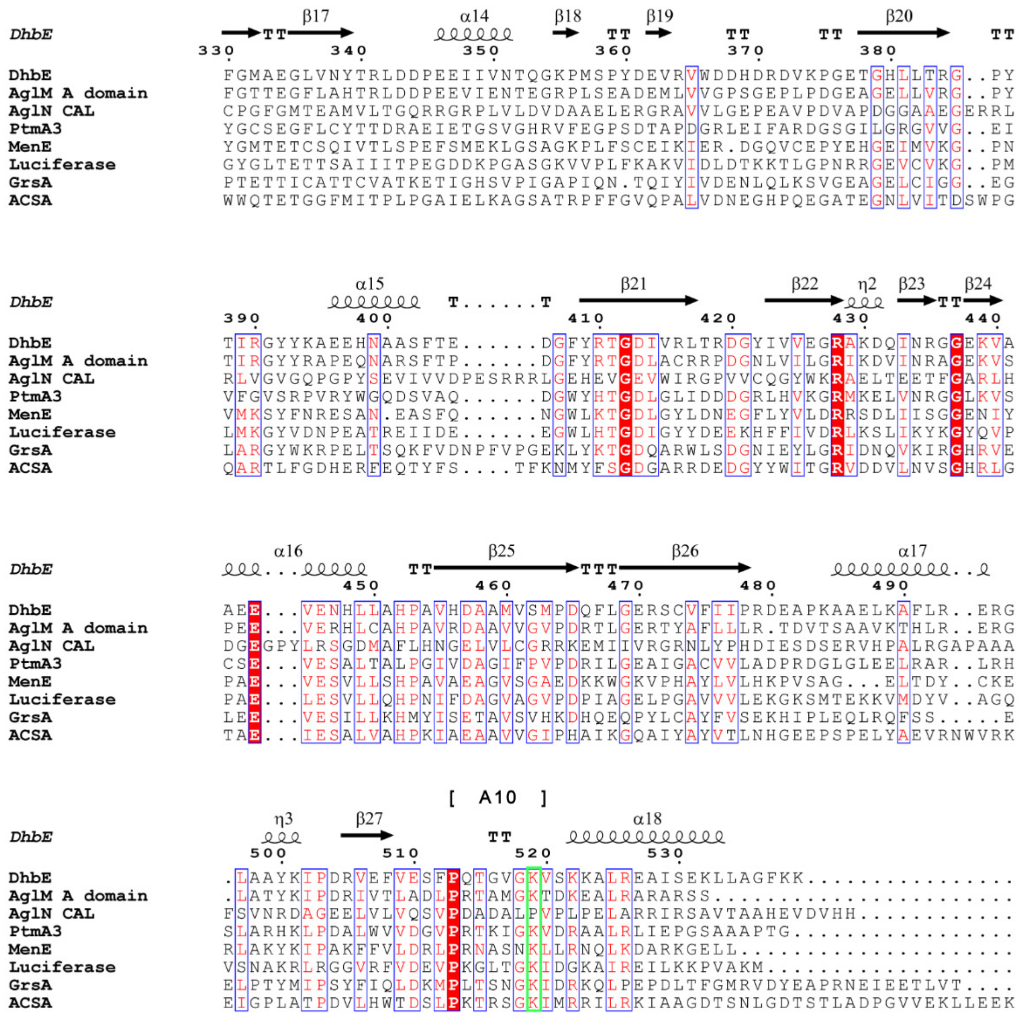

**Figure S35. The amino acid sequence of alignment of CAL, A domain and selected acyl-CoA.** Aligned residues were colored based on the level of conservation (red box with white character showed strict identity, red character showed similarity, and blue frame showed similarity across groups). The corresponding secondary structure of DhbE (PDB entry 1MDF) was depicted above the sequence alignment. The proteins PtmA3 (AIW55578.2), MENE(P23971), luciferase (UUA44467.1), ACSA (Q8ZKF6), and GrsA (CAE7747298.1) are from *Streptomyces platensis*, *Bacillus subtilis*, *Aquatica lateralis*, *Salmonella typhimurium*, and *Brevibacillus brevis*, respectively. Core motif sequences A10 were listed in brackets. The key amino acid residues in the A10 motif were highlighted in green. The alignment was created with ClustalW 2.1 and visualized with ESPript 3.014.

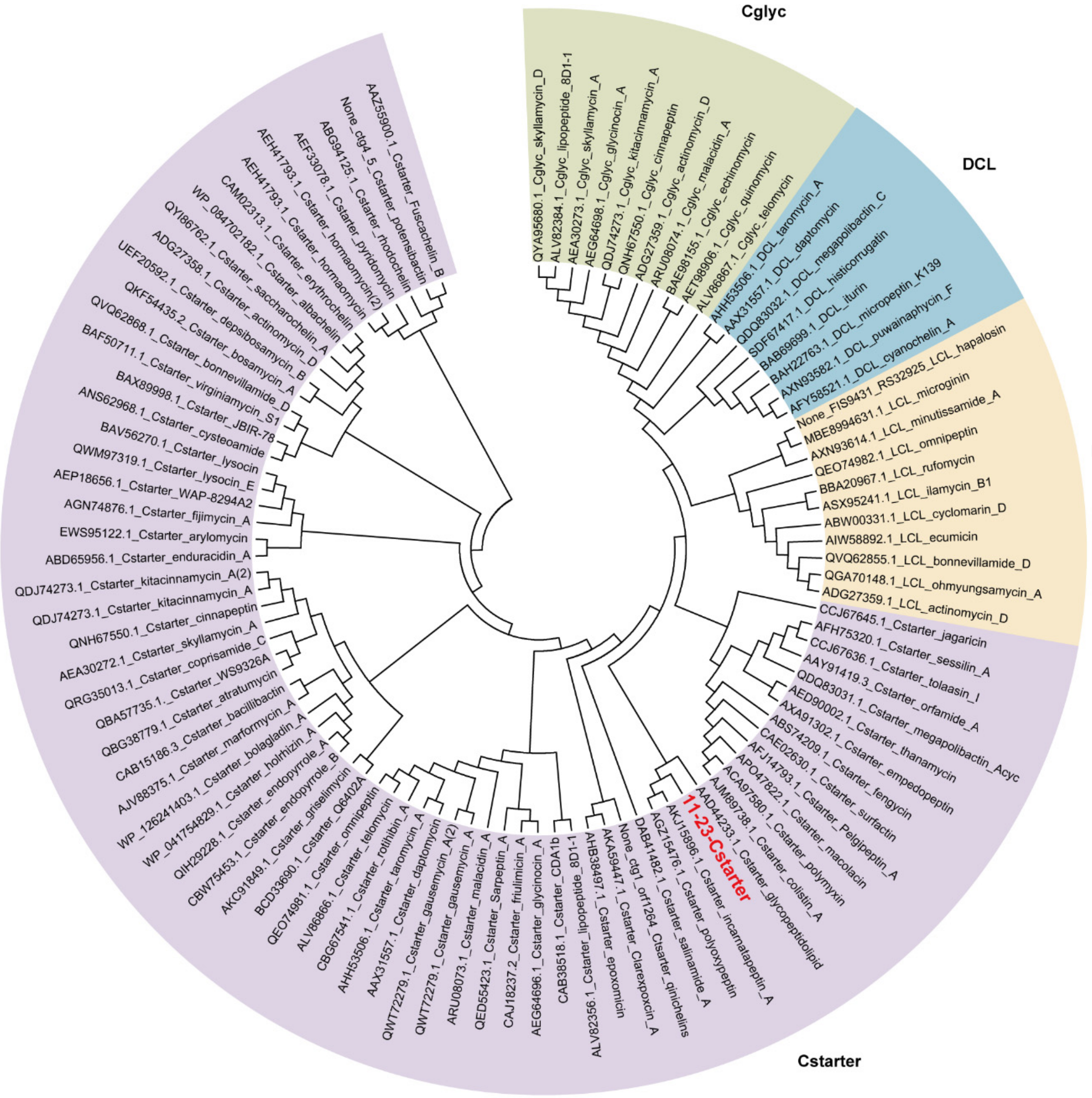

**Figure S36. Phylogenetic tree of condensation (C) domain.** The alignment was created with MUSCLE. The phylogenetic tree was constructed using the Neighbour-Joining (NJ) method. Visualization was conducted with MEGA version 7. In the figure, the C domains are divided into four categories, Cglyc, DCL, LCL, and C$_{starter}$, which are highlighted in green, blue, yellow, and purple, respectively.

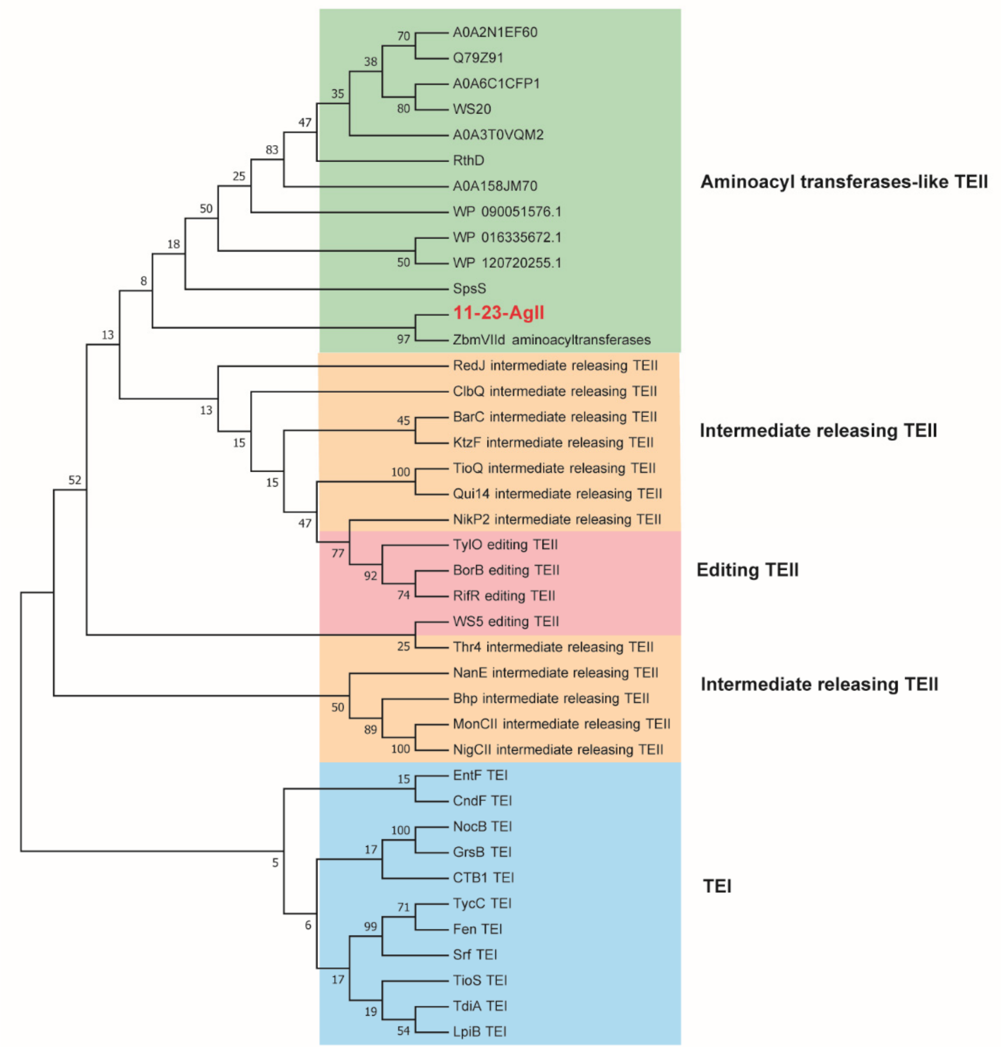

**Figure S37. Phylogenetic tree of thioesterases (TE).** TEI domains, intermediate releasing TEII, editing TEII and aminoacyl transferases-like TEII were highlighted in blue, yellow, pink and green, respectively. The alignment was created with MUSCLE. The phylogenetic tree was constructed using the NJ method. Visualization was conducted with MEGA version 7.

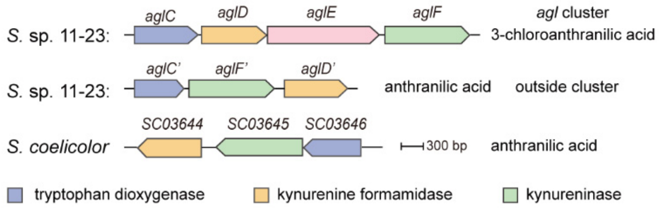

| *S.* sp. 11-23 | AglC | AglD | AglF | AglC' | AglD' | AglF' |
|---|---|---|---|---|---|---|
| *S. coelicolor* | SCO3646 | SCO3644 | SCO3645 | SCO3646 | SCO3644 | SCO3645 |
| Identities (%) | 38.97 | 29.47 | 48.49 | 85.07 | 72.88 | 74.76 |

**Figure S38. Comparison of the triad of AglC/D/F with their paralogs (AglC'/D'/F') and the known *L*-Trp-metabolizing enzymes (SC03646/03644/03645).** The AglC/D/F, predicted to be responsible for 7-Cl-Trp metabolism, exhibit low sequence similarity with their paralogs AglC'/D'/F' which are proposed to be involved in metabolism of *L*-Trp. In contrast, AglC'/D'/F' show high similarity to the known *L*-Trp-metabolizing enzymes SC03646/03644/03645, strongly suggesting their involvement in *L*-Trp metabolism.

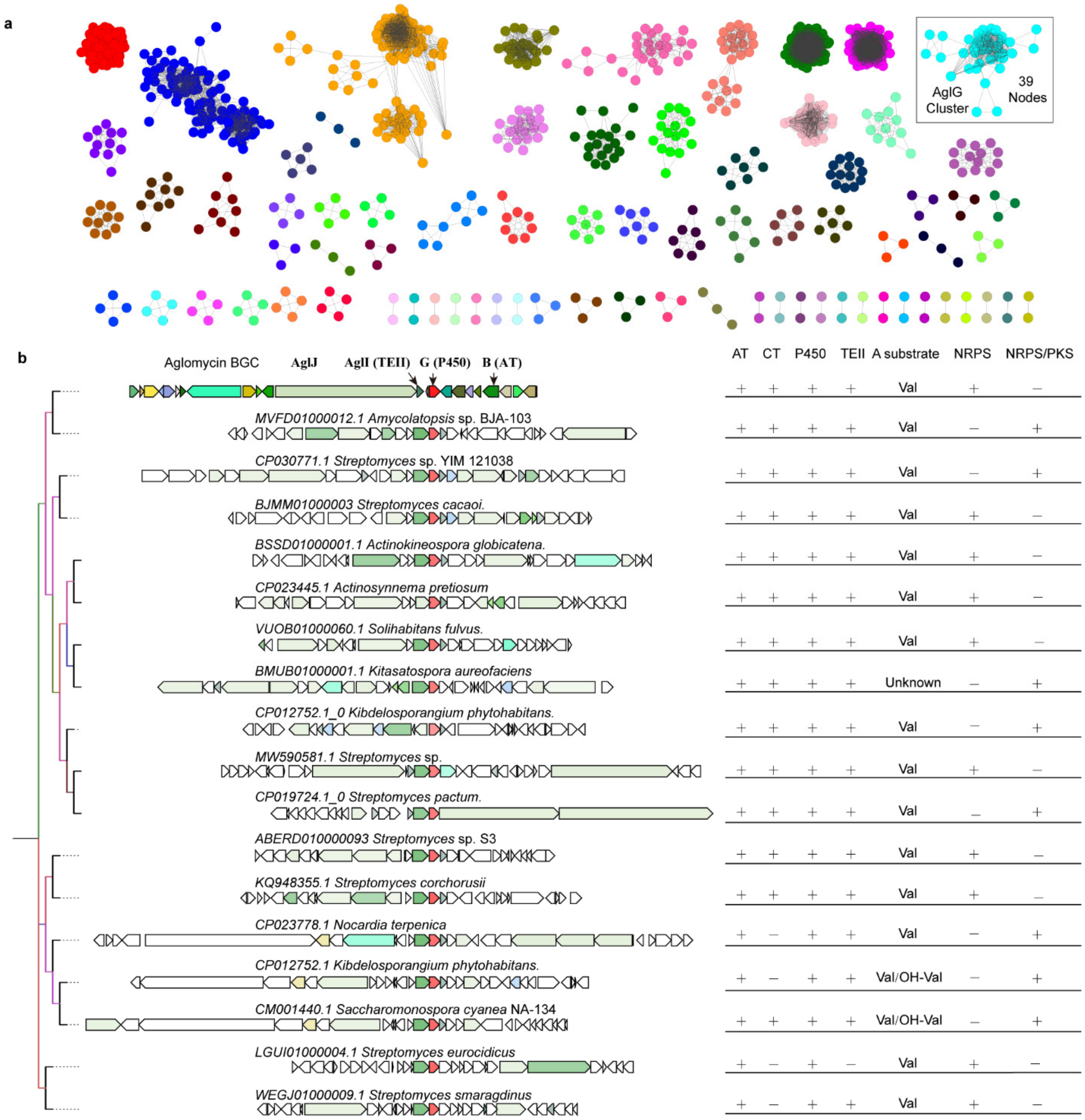

**Figure S39. Genome mining of P450 oxygenases using AglG as a probe. a**, Sequence Similarity Network (SSN) Analysis. The top 1,000 P450 oxygenases were retrieved from the UniProt database (updated to May 2023) using AlgG from the *agl* cluster as a query (E-value < $10^{-5}$). These sequences were used to generate a protein SSN with an alignment score threshold of $10^{-140}$ via the Enzyme Function Initiative-Enzyme Similarity Tool (EFI-EST). A total of 39 sequences clustered together with AlgG, as highlighted in the box. **b**, Genome neighborhood diagrams of the AglG cluster. Genomes corresponding to the AglG cluster were identified using EFI-GNT (Genome Neighborhood Tool) and subsequently analyzed with antiSMASH. CORASON analysis was then performed using an E-value cutoff of $10^{-80}$, with duplicate and edge clusters removed. Eighteen BGCs were identified, with most featuring a conserved gene arrangement comprising an A-T didomain (predicted to selectively activate valine or OH-valine), a P450, a TEII, and an A-less C-T didomain module like the organization seen in the *alg* cluster. Additionally, these genes were found in clusters alongside diverse NRPS or PKS/NRPS-encoding genes. These results suggest that the occurrence of epoxide valine in natural products may be underestimated, and the AglG sequence offers an effective genetic probe for their systematic discovery.

#### The synergistic antibacterial effects of aglomycin A (1) and linezolid

##### MIC determination

The MIC values of aglomycins were determined using the broth microdilution method[46]. *Enterococcus faecium* ATCC 35667, *Staphylococcus aureus* ATCC 29213, *Enterococcus faecalis* (clinical isolated), *Escherichia coli* ATCC 25922, *Klebsiella pneumoniae* ATCC BAA2470, *Acinetobacter baumannii* ATCC 19606, *Pseudomonas aeruginosa* ATCC 27853 and 9 clinical isolates of *Enterococci* were cultured overnight at 37 °C on TSA plates. The bacterial suspensions were diluted to a concentration of $5 \times 10^5$ CFU/mL using fresh MHB medium, and 100 $\mu$L of each suspension was transferred into the wells of a 96-well microplate. Aglomycins, dissolved in DMSO at a concentration of 6.4 $\mu$g/$\mu$L, were added at 2 $\mu$L per well in the first column containing the bacterial suspension. Serial two-fold dilutions were then performed to achieve a concentration range for aglomycins from 0.0625 $\mu$g/mL to 64 $\mu$g/mL. The plates were incubated at 37 °C for 16 – 18 hours, and the MIC was defined as the lowest concentration of aglomycins at which no visible bacterial growth was observed.

##### Cytotoxicity assay

The cytotoxic effects of compounds were determined on various cell lines, including Huh7 (Human hepatocellular carcinoma), HEK293T (human embryonic kidney) and Vero E6 (monkey kidney). Cells were seeded into 96-well plates at a cell density of 10,000 cells per well for Huh7, HEK293T, and Vero E6 cells. After 24 h, the test compounds, with an initial concentration of 64 $\mu$g/mL, were serially diluted twofold for eight dilutions and co-incubated with cells for 24 h. The cytotoxic effects of compounds on various cells were measured by adding 10 $\mu$L cell counting kit-8 (CCK-8) solution (Vazyme Biotech) and incubating for 1 h. The absorbance value was determined at 450 nm in a Victor X5 Plate Reader (PerkinElmer, Waltham, MA, USA). Cell viability was expressed as a percentage of vehicle control, and 50% cytotoxic concentration ($CC_{50}$) was calculated using GraphPad Prism 8.0.

**Checkerboard assay for determining synergistic activity**

*E. faecium*-VRE-1-*van*A was cultured overnight at 37 °C on TSA plates. The bacterial suspension was diluted to a concentration of $5 \times 10^5$ CFU/mL using fresh MHB medium, and 100 $\mu$L of the suspension was added to each well. A checkerboard dilution method was employed to calculate the required final concentration for each well, and the appropriate compounds were added. After thorough mixing, the plates were incubated at 37 °C for 18 hours then analyzed at 600 nm using a Tecan Infinite Spark microplate reader (Tecan, Switzerland). A blank control (MH medium without bacteria) and a positive control (MH medium with bacteria but without antibacterial compounds) were included. Fractional inhibitory concentration (FIC) indices were used to assess the synergistic effects of two antibacterial compounds[47]. The FIC index value was calculated using previously defined formula[48]:

$$FIC\ index = \frac{A}{MIC_a} + \frac{B}{MIC_b}$$

where A and B represent the MICs of each antibiotic when used in combination (within a single well), and $MIC_a$ and $MIC_b$ denote the MICs of each drug individually. The inhibition growth rate was calculated using the following formula:

$$\text{Inhibition Growth Rate (\%)} = \left(\frac{OD_{600}^{\text{positive}} - OD_{600}^{\text{sample}}}{OD_{600}^{\text{positive}} - OD_{600}^{\text{blank}}}\right) \times 100\%$$

where $OD_{600}$ the optical density at 600 nm. The MIC was defined as the minimum concentration required to inhibit 95% of bacteria. The synergistic effect is indicated by an FIC index below 0.5[49].

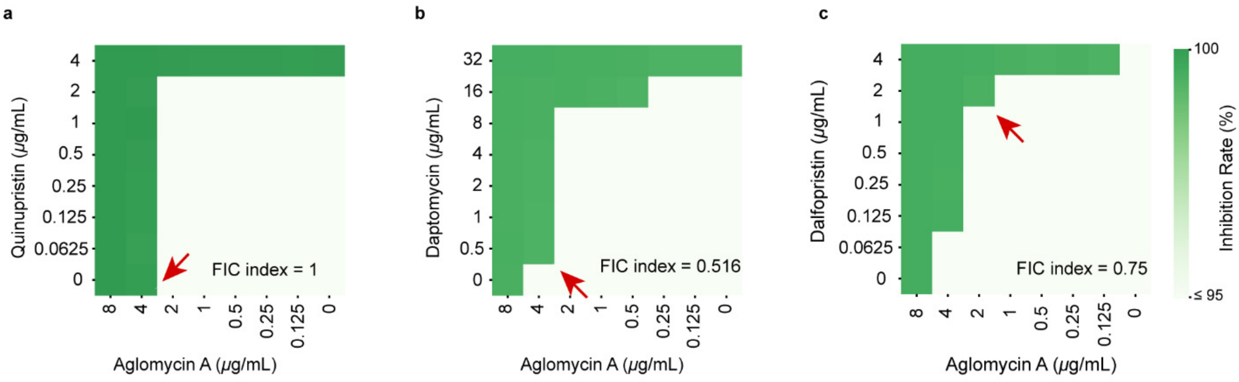

**Figure S40. Checkerboard assay of aglomycin A with quinupristin (a), dalfopristin (b), and daptomycin (c) against *E. faecium*; dark green indicates higher inhibition.**

**Time-dependent killing assay**

We assessed the antimicrobial mode using the time-dependent killing assay. *E. faecium* ATCC 35667 was cultured in TSB medium at 220 rpm and 37 °C to reach the exponential growth phase, then was adjusted to 5 × $10^5$ CFU/mL in MHB medium, with 200 μL added to each well of a 96-well microplate. The stock solution of each compound in DMSO (0.8 or 3.2 mg/mL for aglomycin A and 0.1 or 0.4 mg/mL for linezolid) was added either individually or in combination to the bacterial suspension, achieving final concentrations equivalent to 0.5–8× the MIC of the respective compound. A blank control was established by adding 2 μL of DMSO to the bacterial suspension. The 96-well plate was placed in a 37°C incubator for static incubation, with 20 μL of each sample taken every two hours and diluted 10,000-fold for colony counting.

***In vivo* antimicrobial efficacy assay**

We evaluated the *in vivo* antimicrobial activity using a *G. mellonella* model. First, *E. faecium* ATCC 35667 was inoculated in LB medium and incubated at 37 °C for 24 hours. The culture was centrifuged and washed with saline to collect the bacterial cells. Then the bacterial suspension was adjusted to 8 × $10^9$ CFU/mL using saline. For each group, 20 healthy, milky-white last-instar larvae of *G. mellonella* were selected. A 5 μL aliquot of the bacterial suspension were injected into the hemolymph of each larvae using a Hamilton syringe (7654-01, Hamilton) equipped a 33G RN needle (10 mm, Hamilton). The larvae were then incubated in a dark environment at 37 °C and 50% humidity for one hour. Subsequently, aglomycin A, linezolid, or a control solution of physiological saline containing 9.3% DMSO were separately injected into the infected larvae. Uninfected larvae were also included as a blank control group. The larvae were maintained in an incubator, and their survival was monitored every 24 hours. Larvae that showed no response to needle stimulation were considered dead. The experiment lasted five days, and survival rates were recorded. Kaplan-Meier survival curves were then plotted. Cox regression analysis was performed using R software (version 4.2.2), along with MSTATA software (https://www.mstata.com/).

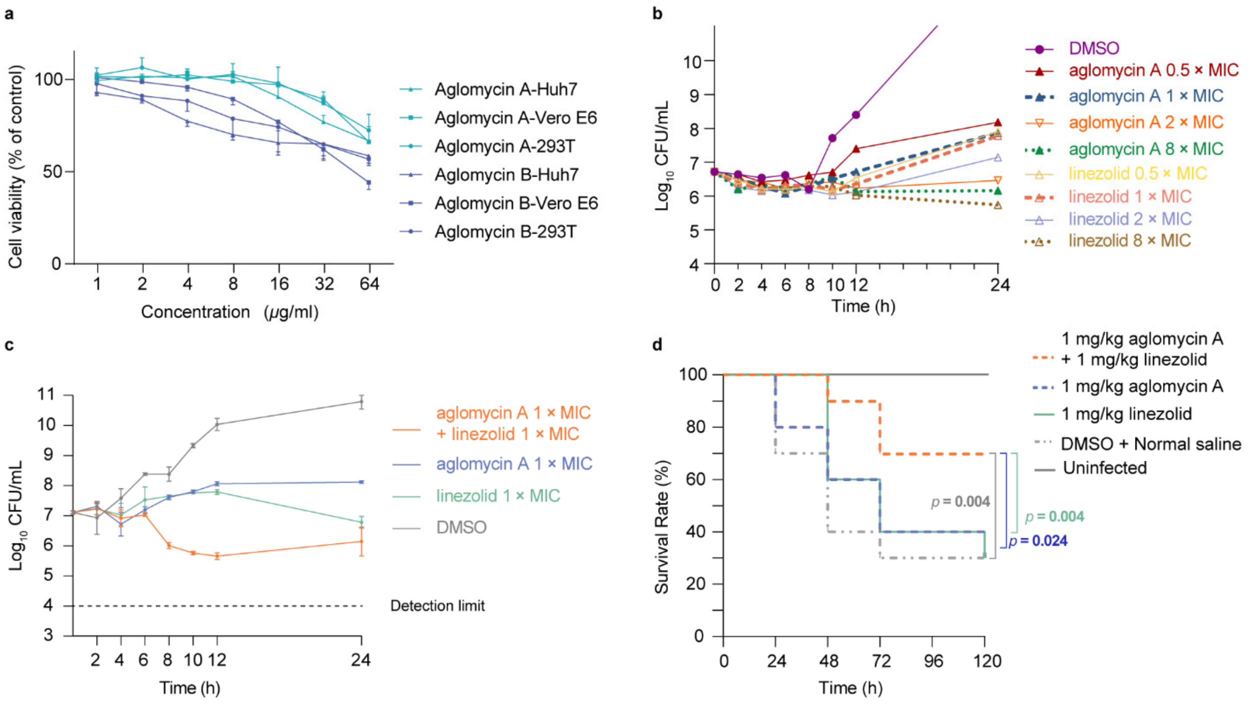

**Figure S41. Cytotoxicity of aglomycins and synergistic effects of aglomycin A and linezolid. a**, Cytotoxicity of aglomycins A and B on Huh-7, Vero E6, and 293T Cells. The $CC_{50}$ values were 159.1 and 106.9 μg/mL; 88.1 and 50.2 μg/mL; 108.9 and 90.3 μg/mL, respectively. Data are mean ± s.d. of three replicates. **b**, Schematic diagram of the time-killing curve of aglomycin A and linezolid alone on *E. faecium* ATCC 35667. Both show time-dependent bacteriostatic activity. **c**, Schematic representation of the time-killing effect of aglomycin A in combination with linezolid alone and in combination against strain *E. faecium* ATCC 35667. The combination exhibits a certain degree of bactericidal effect. **d**, Protective effects of aglomycin A 1 mg/kg and linezolid 1 mg/kg alone and in combination against *E. faecium* ATCC 35667-infected *G. mellonella* larvae. (Each group contained 20 larvae.) Cox regression analysis was performed using R software (version 4.2.2), along with MSTATA software (https://www.mstata.com/).
